# Modeling echolocation as an active pursuit of information via infotaxis

**DOI:** 10.64898/2025.12.31.697206

**Authors:** Wu-Jung Lee, John R. Buck, Peter L. Tyack

## Abstract

Echolocation is a closed-loop active sensing modality in which animals not only choose how they move to acquire information, but also actively modulate incoming sensory (echo) information by shaping the acoustic signals they emit to probe the environment. While many models describe how echolocating animals *react* to prior echoes by adjusting subsequent behavior, few explicitly model how they cognitively *reason* about information embedded in echoes when determining future actions. Here, we extend “infotaxis,” an information-greedy algorithm originally developed for olfactory search, to sonar sensing by formulating an echolocating agent searching for a single target under sensory uncertainty characterized by probabilities of miss and false alarm. Through analytical and computational analyses, we show that the characteristic exploration-exploitation balance of infotaxis also emerges in echolocation, and that the efficiency and reliability of infotaxis search depend strongly on sensory information quality. Compared with a maximum *a posteriori* agent that always directs the beam to the most probable target location, the infotaxis agent consistently completes searches with fewer pings and greater robustness to sensory uncertainty. These results highlight information as a powerful concept for understanding active sensing and developing models for sonar-guided autonomy in both biological and engineered systems.

**AUTHOR SUMMARY:** Echolocating animals such as bats and dolphins actively probe their surroundings by emitting sounds and interpreting the returning echoes. They must continually decide where to direct their active probing to gather useful information. Although many existing theoretical models describe how echolocators react to echoes, few explain how they might reason about uncertain echo information when choosing their next action. In this study, we extend infotaxis, an information-seeking search strategy originally developed for olfactory sensing, to echolocation. We developed a computational model of an echolocating agent searching for a target while accounting for imperfect sensory information, including missed detections and false alarms. At each step of the search, the model evaluates possible sensing actions and selects the one expected to maximize the reduction in uncertainty about the target’s location. Our simulations showed that the echolocating infotaxis agent naturally strikes a balance between exploration of new regions and exploitation of previously observed regions. Moreover, infotaxis searches are more reliable and efficient than a simpler strategy that always focuses on the most likely target location, particularly when sensory uncertainty is high. These results highlight the importance of information-based decision making in active sensing and provide a framework for studying sonar-guided behavior in animals and engineered systems.

## I. INTRODUCTION

Echolocation is a sensory modality in which echolocators emit acoustic signals and extract information from the returning echoes to understand their surroundings. Bats and toothed whales are two prominent animal groups that have evolved specialized anatomical structures and neuro-physiological mechanisms to employ echolocation as their primary sensing modality to navigate and forage (Surlykke *et al*., 2014). Echolocation is a closed-loop feedback system in which the echolocator adjusts its outgoing signals and movements based on information derived from previous echo returns to achieve its behavioral goal (Fig 1A). It is an “active” sensing modality, both in the narrow sense that the echolocator actively transmits acoustic energy to interrogate the environment and in the broad sense that the echolocator actively changes the configuration of its sensing apparatus (e.g. ears), location (e.g., body position with respect to a target), and cognitive strategies (e.g., attention modulation of the incoming echo-acoustic scene) to acquire information (Moss *et al*., 2023; Zweifel and Hartmann, 2020).

**FIG 1:**
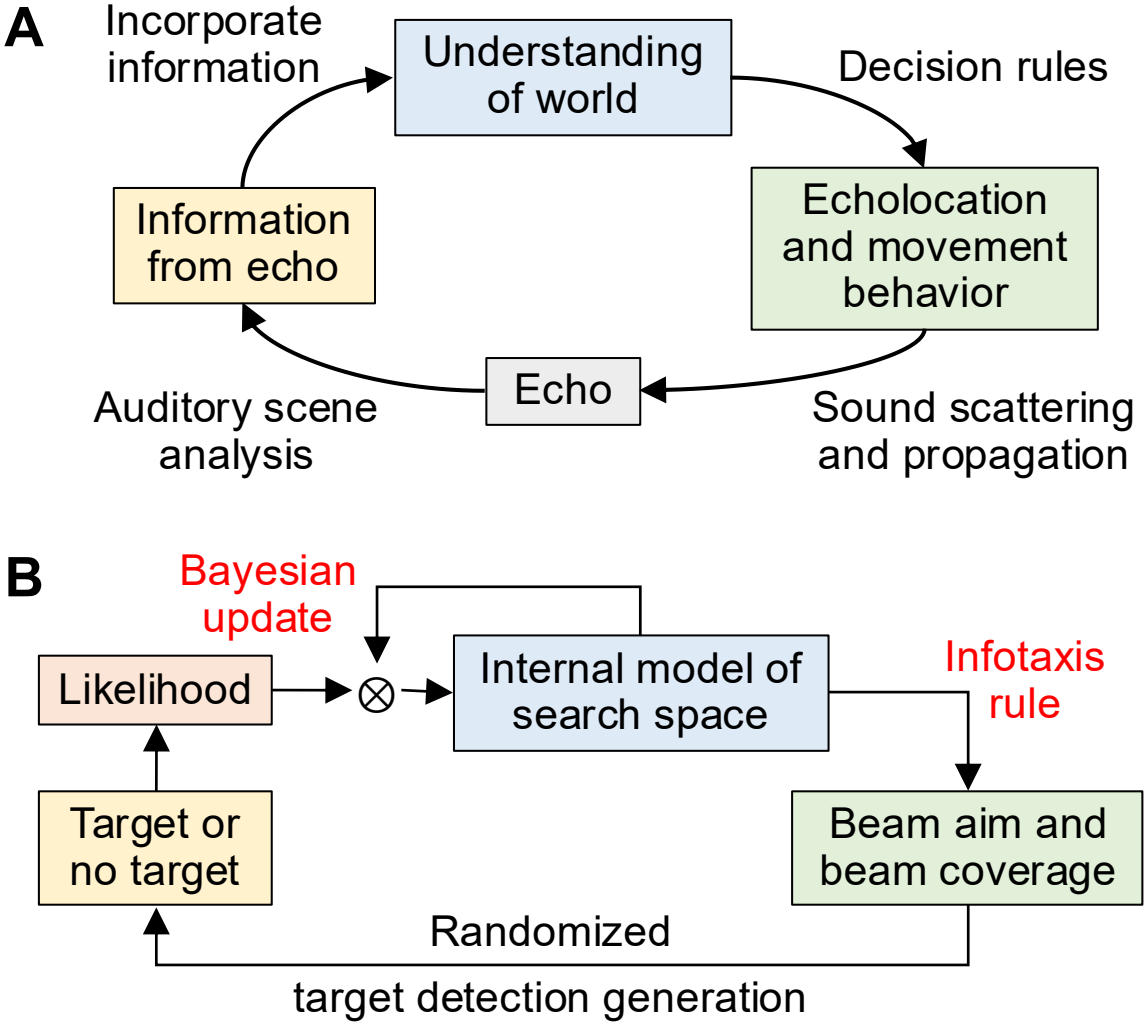
Echolocation as a form of active sensingn. (A) An echolocating agent probes its surroundings by emitting sound and extracting information from the echoes to update its understanding of the world. (B) The active sensing echolocation feedback modeled in this paper. The agent uses the infotaxis rule to determine the next beam aim and beam coverage to locate a target in the search space. New echo information is incorporated into the agent’s internal model of the search space through a likelihood function via Bayesian updates.

Decades of research in both laboratory and field settings have revealed how bats and toothed whales adapt echolocation behaviors to better solve problems as the task and environment change (Moss *et al*., 2023). For example, bats modulate pulse duration and inter-pulse intervals according to their attended range and clutter around the target (e.g., Aytekin *et al*., 2010; Falk *et al*., 2014; Wilkinson *et al*., 2025), adjust flight trajectory and call emission patterns in response to environmental clutter (e.g., Ding *et al*., 2023; Taub and Yovel, 2020; Wheeler *et al*., 2016), and alter the spatial coverage of sonar transmissions in a task-dependent manner, broadening the beam to ensure successful prey interception and narrowing the beam to assist with localization and landing (e.g., Eitan *et al*., 2022; Matsuta *et al*., 2013). Similarly, toothed whales modify click amplitude and inter-click intervals depending on the target range and environmental reverberation (e.g., Christman *et al*., 2024; Johnson *et al*., 2004; Ladegaard and Madsen, 2019; Wisniewska *et al*., 2012), switch from emitting single clicks to packets of clicks to detect distant targets (Finneran, 2013; Ladegaard *et al*., 2019), employ highly directional sonar beams to discriminate closely spaced objects (Malinka *et al*., 2021), broaden the beam to ensure successful prey interception (Wisniewska *et al*., 2015), and adjust their movement trajectories depending on echo ambiguity (Lee *et al*., 2025). Many theoretical models have been developed to describe adaptive changes in acoustic and movement behavior during echolocation. Hand-crafted analytical models have been proposed to describe bats’ movements and sonar parameter adjustments during prey tracking and interception (e.g., Aihara *et al*., 2013; Fujioka *et al*., 2016; Ghose *et al*., 2006; Ghose and Moss, 2006; Mansour *et al*., 2019; Nguyen *et al*., 2021; Nishiumi *et al*., 2024; Salles *et al*., 2020; Sumiya *et al*., 2017). Many studies further integrated auditory perceptual cues from echoes together with spatial information derived from the bat’s self-motion and/or the coordinated movement of its sensing apparatus to model obstacle avoidance (Vanderelst *et al*., 2015), prey capture (Vanderelst and Peremans, 2018), global map construction via repeated flights over acoustical landmarks (Van-derelst and Peremans, 2017), and the coordinated movement of foraging bat pairs (Giuggioli *et al*., 2015). In parallel, information theoretic approaches examined how echolocating animals may exploit frequency-dependent receiving beampatterns and ear motion to enhance echo detection and localization (reviewed in Müller *et al*., 2017), or to improve localization accuracy by directing sonar beams off-axis (Arditi *et al*., 2015; Kloepper *et al*., 2018; Yovel *et al*., 2010). More recently, reinforcement learning and neural network-based approaches were applied to model how bats partition resources in the wild (Goldshtein *et al*., 2020), adjust decision strategy according to environmental dynamics (Naamani *et al*., 2023), avoid obstacles while pursuing food items in computational simulations (Mohan and Vanderelst, 2020; Nguyen and Vanderelst, 2022, 2025), and negotiate obstacles under species-specific constraints in laboratory experiments (Teshima *et al*., 2025).

Despite substantial progress, a clear gap persists between existing modeling approaches and the interpretation of behaviors observed in echolocation experiments. In many models, the echolocator’s behavior is governed by a fixed set of movement and sonar parameter tuning rules that respond directly to sensory stimuli, including echoes from prey, environment features, or conspecific calls. These models emphasize the *reactive* aspects of sensorimotor adjustments in echolocation. The cognitive dimensions of echolocation, where the animal actively *reasons* about the echo-acoustic scene and determines its next set of actions after integrating information from prior echo returns (Fig 1A), remain under-explored, even though such cognitive reasoning is often invoked when interpreting behavioral data. For example, variations in pulse duration and inter-pulse intervals in bats, as well as adjustments of inter-click intervals in toothed whales, are commonly interpreted as evidence that these animals modulate their acoustic attention or field of view in complex environments. While recent deep reinforcement learning (RL) models have begun to address this gap by introducing explicit cognitive architectures to enable agents to learn actionable policies from echo returns and rewards, the interpretability of such models remains challenging (Glanois *et al*., 2024). As new discoveries continue to underscore the remarkable flexibility of animal echolocation behavior, there is a growing need for modeling frameworks that integrate the physical and physiological properties of these biological sonar systems with the associated sensory and cognitive processes to yield mechanistic insights into adaptive sonar-guided actions.

“Infotaxis” has been an influential model for describing the search for odor sources in turbulent environments (Vergassola *et al*., 2007), in which the agent updates an internal “world” model of probable source locations through a sensory model that encodes the likelihood of odor detection within and around a plume. The agent then chooses its next movement by maximizing the expected information gain at each step. While the observable behavior (agent movement following odor detection) appears similar to reactive sensorimotor models, infotaxis explicitly incorporates cognitive reasoning by specifying how sensory information should be used to update the internal world model and select the next action, providing an interpretable bridge between behavioral observations and model parameterization. This information-based formulation has been extended to a variety of odor sensing scenarios, including multi-agent schemes (Hajieghrary *et al*., 2016; Karpas *et al*., 2017; Masson *et al*., 2009). The infotaxis model behavior has also been compared to the behaviors of worms, flies, mosquitoes, and moths in laboratory experiments (e.g., Calhoun *et al*., 2014; Hernandez-Reyes *et al*., 2021; Pang *et al*., 2018). In vision, similar information-based models have been proposed to model eye movements (saccades) during search and categorization tasks and compared with experimental data through summary statistics and on a fixation-by-fixation basis (Jang *et al*., 2021; Najemnik and Geisler, 2005, 2009; Yang *et al*., 2016). Theoretical analyses and simulations have also examined the efficiency and reliability of these algorithms under realistic biological constraints, such as the energetic cost of movement (Chen *et al*., 2020; Loisy and Eloy, 2022).

Here, we built upon these models to frame echolocation as an active pursuit of information, extending the infotaxis algorithm to an active sensing modality that shares both similarities and differences with olfaction and vision. Similar to olfaction and vision, echolocation is a remote sensing modality in which information can be acquired from a distance. However, in olfaction and vision, even though the animals have control over their sensors [e.g., antenna movements and eye fixations (Claverie *et al*., 2023; Kümmerer and Bethge, 2023)] to achieve better sampling and accentuate specific features in the incoming signals, they do not have control over the odor molecules or the light spectrum they receive. In contrast, echolocating animals exert active and direct control over the spectral, temporal, and directional characteristics of their transmit signal, allowing them a greater capability to selectively sample and emphasize environmental or target features most relevant to its behavioral goals.

We first developed a general Bayesian framework for echolocation that incorporates detection parameters specific to active sonar sensing. We then formulated a detailed model of an echolocating agent searching for a single target by steering a narrow sonar beam within a finite frontal search space. An important contribution of this paper is that our sensing model used a binary hypothesis test for echolocation-based target detection with sensory uncertainty captured by the probabilities of miss (*P*_*M*_) and false alarm (*P*_*FA*_), which are important intrinsic characteristics of sonar systems (Van Trees, 1968). This differs from the Poisson point process detection model in the original infotaxis paper, where detection probabilities were based on plume concentrations (Vergassola *et al*., 2007). Despite its simplicity, this modeling scenario generated a variety of search behaviors, demon-strating that the characteristic infotaxis balance between exploration and exploitation in following odor plumes also emerges in echolocation, and that the search trajectories and performance depend fundamentally on sensory information quality. Simulations further showed that infotaxis is more reliable and efficient compared with the maximum *a posteriori* (MAP) rule, under which the agent always directs its sonar beam to the most probable target locations.

The remainder of this paper is organized as follows. Sec. II presents the general formulation of the echolocation infotaxis framework, a detailed formulation of the infotaxis and MAP agents in a single target search scenario, and the corresponding computational implementation. Sec. III describes typical search dynamics of the infotaxis agent, including how entropy decreases as the agent explores the search space, how sensory uncertainty influences infotaxis selection of beam aim and beam footprint size, as well as comparisons of how infotaxis and MAP searches perform. Finally, Sec. IV discusses the broader implications of our results, potential extensions of the model, and its relevance to both biological and engineered sonar systems.

## II. METHODS

### A. General formulation of infotaxis in echolocation

In the infotaxis framework, we use probabilistic representations for quantities in typical echolocation tasks, including target search and discrimination. In a target search task, let *K* be a random variable representing the target location in the search space *A*, 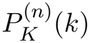 describes the echolocating agent’s understanding of the search scene after the *n*th echolocation transmission (or the *n*th “ping”) as a probabilistic distribution over all possible target locations. Similarly, in a target discrimination task, let *K* be a random variable representing the target identity among all possible categories *A*, 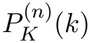 describes the agent’s belief about target identity after the *n*th ping. In both scenarios, 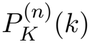 encapsulates the agent’s knowledge about the target—whether its location or identity—given all prior echo observations (evidence) from the first to the *n*th transmission.

In addition, we define *S* as the set of acoustic and movement actions available to the echolocating agent before or during the next ping, such as sonar beam direction and beamwidth, spectral and temporal content of the emitted signal, subsequent body location and head orientation, to name a few. Similarly, we define *X* as the set of echo measurements (features) the agent receives from the previous ping. These features are typically multidimensional, encompassing temporal, spectral, and statistical properties of the echoes that vary depending on the relative position between the echolocating agent and the target, as well as the acoustic characteristics of the environment.

The echolocating agent’s belief of the target location or target identity is iteratively updated using Bayes’ rule upon receiving an echo observation from the last transmission

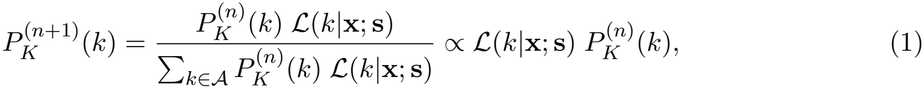

where *ℒ* is the likelihood function, representing the probability of receiving an echo observation with features **X** = **x** given a set of actions **S** = **s**, when the target is at location *k* or the target identity is in category *k*,

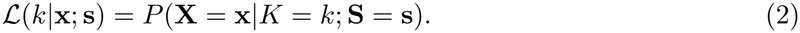

The likelihood function provides a convenient entry point for incorporating the auditory perceptual capabilities of echolocating animals, such as temporal and spectral resolution and directional hearing ambiguity, into the model. These capabilities are typically characterized through behavioral and neurophysiological experiments (e.g., Branstetter *et al*., 2022; Mulsow *et al*., 2024; Nachtigall *et al*., 2000) and can be represented as probabilistic quantities to allow flexible parameterization of sensory and environmental noise influences.

Under infotaxis, the echolocating agent determines its next set of actions by maximizing the *expected* reduction of uncertainty about the key task-related variable *K* (target location or target identity) after receiving echoes from the *next* transmission (Fig 1B). Note this expectation is taken over the agent’s belief after the possible update, not the unknown true probability. Using the Shannon information entropy as a measure of uncertainty (Shannon, 1948), the amount of information contained in the received echo can be quantified by the mutual information between the current and the updated distributions of *K*,

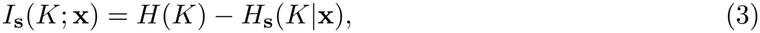

where *H*(*K*) is the entropy of the current distribution 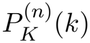 and *H*_**s**_(*K* | **x**) is the entropy of the updated distribution *P*_*K*|**X**;**s**_(*k*|**x**; **s**) given action **s** and echo observation **x**,

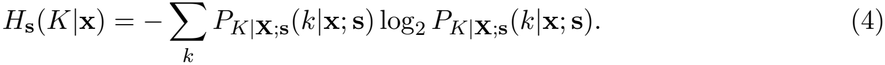

Since the echo observation is not known *a priori*, the echolocating agent operates based on the *expected* uncertainty over all possible echo observations,

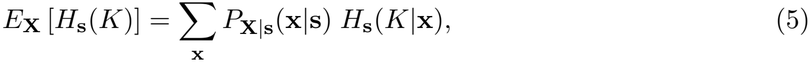

and selects the next set of action **s** that minimizes this expected uncertainty:

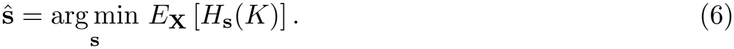

Eq 6 follows directly from Eq 3, since maximizing the mutual information is equivalent to minimizing the expected uncertainty of the distribution.

While the above formulation is general, in the following sections we focus our development in the context of echolocation-based target search.

### B. Infotaxis search for a single target

Building on the general formulation above, in this section we develop a detailed model of an echolocating infotaxis agent searching for a single target in a discrete, two-dimensional search space *A* consisting of *N*_*A*_ cells (Fig 2). This setup conceptually represents an experiment in which an echolocating animal searches for a target on a vertical “screen,” onto which its sonar beam projects a finite footprint. The geometry assumes that the main response axis of the sonar beam is perpendicular to the screen or plane being searched.

**FIG 2:**
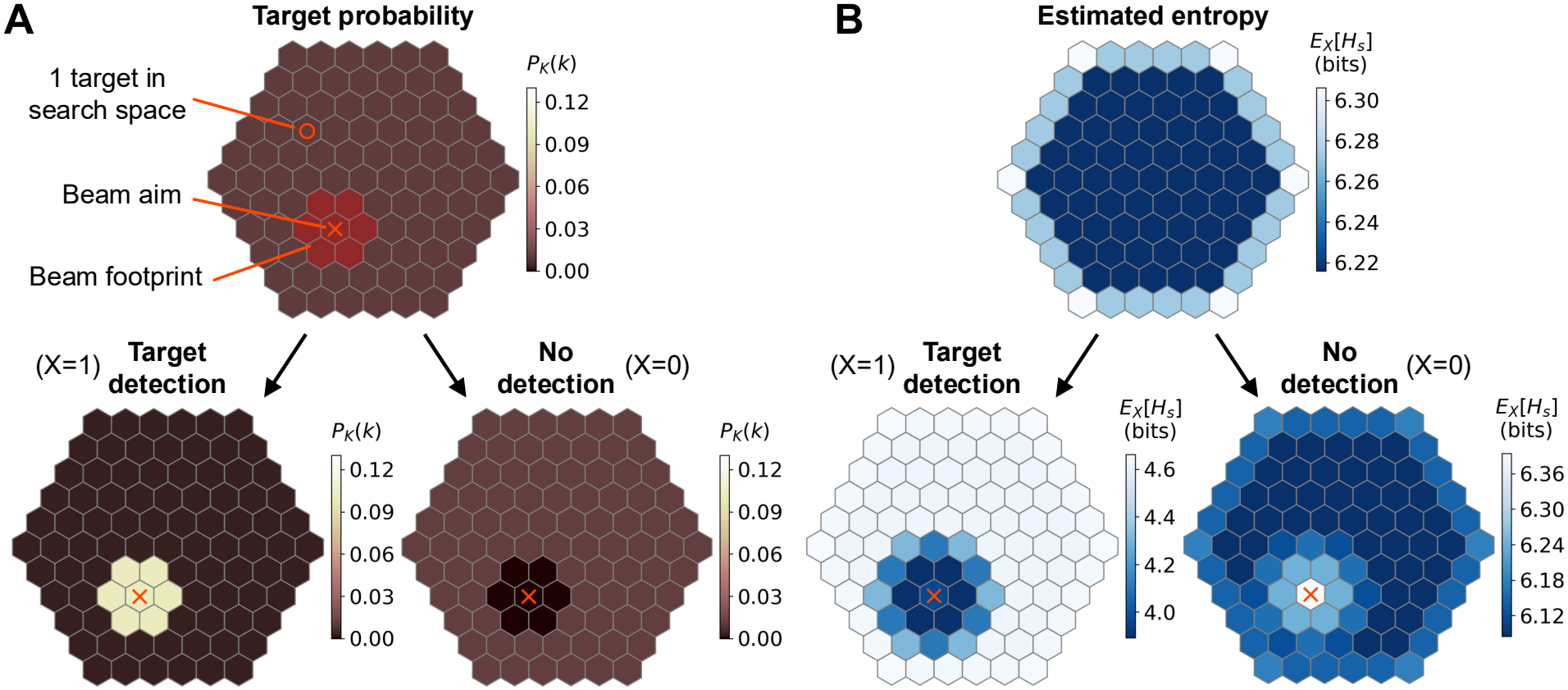
Setup of the echolocation infotaxis model for single target search. Probability distribution of target location 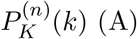 (A) and expected entropy *E*_*X*_[*H*_*s*_] (B) in the beginning of a search (top), after receiving a target detection (lower left), and after not receiving a detection (lower right). *P*_*M*_ = *P*_*FA*_ = 0.005 in this example.

We assume that the echolocating agent can control the beam aim and the size of the beam foot-print, but lacks sufficient directional hearing capabilities to determine where within the footprint a target echo originates. We further assume that auditory scene analysis—the process through which complex sound mixtures are segregated and organized (Bregman, 1990)—and target detection occur automatically, such that the agent receives a binary outcome: a target detection (*X* = 1) or no target detection (*X* = 0) (Fig 1B). Consecutive echo observations are assumed independent across pings given the candidate target location and sensing action. In addition, the agent’s target detection is subject to uncertainty arising from sensory and environmental noises. This uncertainty is characterized by the probability of miss, *P*_*M*_, and the probability of false alarm, *P*_*FA*_. For results presented in the main text of this paper, we use constant and independent *P*_*M*_ and *P*_*FA*_ across cells within the beam to focus our analysis on intrinsic properties of the echolocation infotaxis model. In general, *P*_*M*_ and *P*_*FA*_ can vary across locations within the beam footprint and search space to incorporate the effects of variable signal-to-noise ratio (SNR) due to the directional transmission beampattern and hearing sensitivity, as well as noise interference or environmental clutter. We present preliminary results from beampattern-modulated *P*_*M*_ and *P*_*FA*_ in Section A in S1 Appendix, and note that alternative false alarm models that capture spatially correlated structures given specific sonar system characteristics and the sensing environment could be incorporated in future extensions.

Let **s** = [*θ*_*B*_ *N*_*B*_]^⊤^ denote the echolocation action, where *θ*_*B*_ is the beam aim (beam axis) and *N*_*B*_ is the beam footprint size in cells. Let *ℬ*_**s**_ be the set of cells covered by the beam footprint. When a target detection occurs, the likelihood is

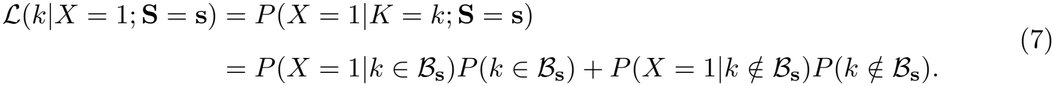

When no target detection occurs, the likelihood is

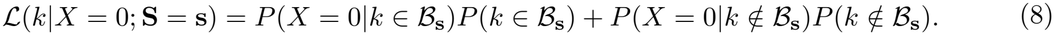

For a given beam footprint, whether or not a particular cell is covered is deterministic. Therefore, *P* (*k* ∈ *ℬ*_**s**_) = 1 when cell *k* is within the beam and *P* (*k* ∈ *ℬ*_**s**_) = 0 when cell *k* is outside of the beam.

Deriving the likelihood functions in terms of the model parameters is simplest for the case when no detection occurs (*X* = 0). First consider the case where the cell to be updated *k* is within the beam footprint:

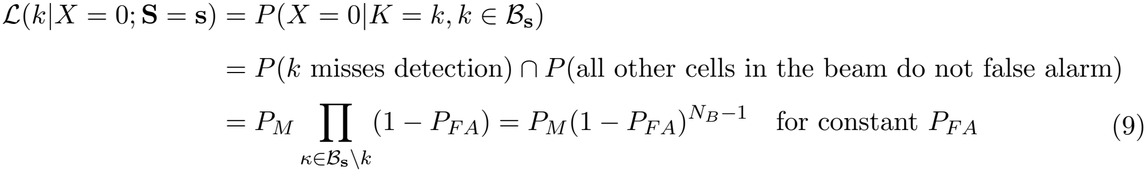

where the set subtraction operation *ℬ*_**s**_ \ *k* denotes the set of cells within the beam footprint except for the cell *k* currently being updated. For cells outside the beam when no detection occurs,

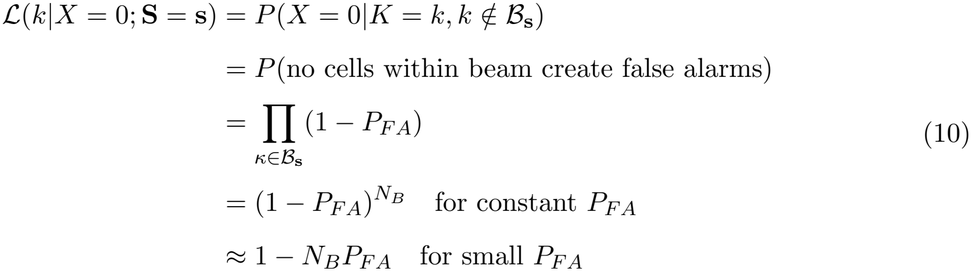

The last step follows from applying the binomial approximation (1 + *y*)^*a*^ ≈ 1 + *ay* for small *y*.

The complementary cases when there is a target detection (*X* = 1) follow directly. When the cells being updated are within the beam,

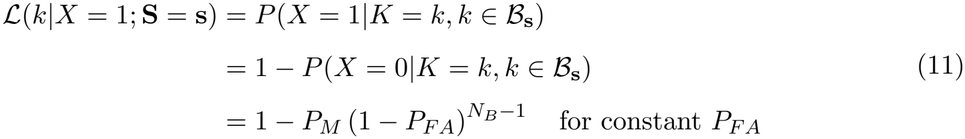

Similarly, for cells outside the beam

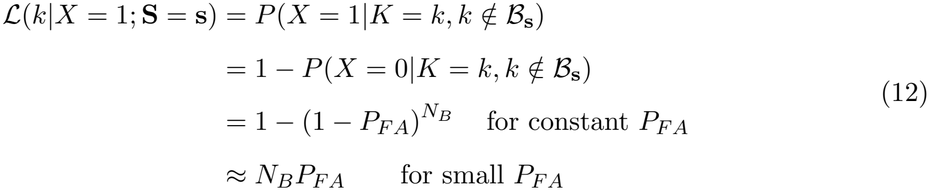

Note the binomial approximation is only used to obtain simplified expressions to aid analytical interpretation. The approximate expressions are not used in the values presented in Sec. III, which are computed from the exact expressions.

Under the infotaxis rule, to determine the next action, we evaluate the expected uncertainty, *E*_*X*_[*H*_**s**_(*K*)] (Eq 5) by computing the conditional entropy *H*_**s**_(*K*|*x*) (Eq 4) and the probability of receiving an echo observation *X* given action **s**,

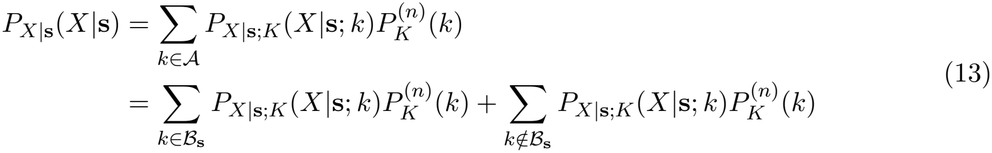

where 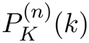 is the agent’s belief of the probability distribution of the target location after ping *n*, and *P*_*X*|**s**;*K*_ (*X*|**s**; *k*) is the probability of receiving echo *X* after action **s** with the target located in cell *k*, which is identical to the likelihood (Eqs 7-8). Here,

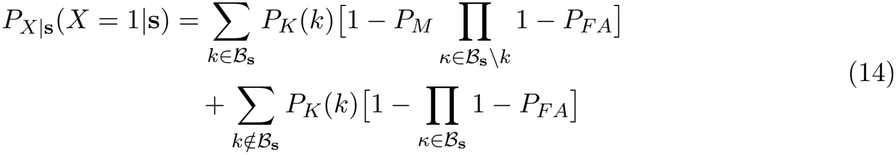

and

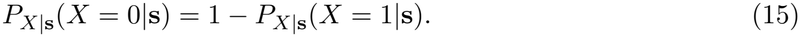

### C. Maximum *a posteriori* (MAP) search for a single target

To gain comparative insights into the properties of echolocation infotaxis, we examine the behavior of another echolocating agent employing a simple MAP rule to select its next beam aim from cell(s) with the highest target probability in the search space:

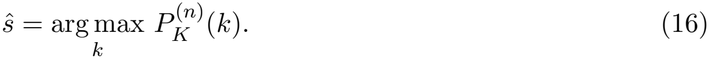

Here, *s* is a scalar, because the MAP does not provide a mechanism to include beam footprint size as a decision variable (i.e., the beam footprint is always fixed within a search). This action selection rule is based solely on the latest probability distribution of target location and, unlike infotaxis, does not account for sensory uncertainty or ambiguity of target location within the beam footprint. In this work, we use the MAP strategy as a simple belief-based exploitation baseline, as it has been employed to model eye fixation selection in visual search for comparison with more sophisticated strategies (e.g., Hoppe and Rothkopf, 2019; Najemnik and Geisler, 2005, 2008).

For all infotaxis and MAP comparisons in this paper, the beam footprint size is fixed in. Apart from the action selection rule, all other aspects of MAP search are identical to those of infotaxis, including the stopping condition (see next section).

### D. Computational implementation

We developed a computational implementation of the single target search scenario described above to systematically examine how an echolocating agent behaves given different action selection rules and search parameters, including sensory uncertainty *P*_*M*_ and *P*_*FA*_, beam footprint size *N*_*B*_, search space size *N*_*A*_, and their relative proportion *α* = *N*_*B*_*/N*_*A*_. In the implementation, the search space is a finite two-dimensional area represented using a hexagonal grid system (Fig 2) defined with radius *r*_*A*_, and the echolocation beam footprint is a circular region with radius *r*_*B*_ centered on the beam aim *θ*_*B*_. Therefore, in figure annotations, *r*_*A*_ and *r*_*B*_ are used in place of *N*_*A*_ and *N*_*B*_, respectively. A hexagonal grid is used, because in this geometry all cells along a rim at a given radius are equidistant from the center beam aim, whereas in rectangular grids, the distances would vary depending on whether the outer cells lie along an edge or at a corner. Under the hexagonal grid setup, the number of cells covered in the area within a given radius *r* is *N* = 1 + 3*r*(*r* + 1).

The agent can operate under either the infotaxis or the MAP rule. Under the infotaxis rule (Eq 6), the agent selects both the next beam aim and beam footprint radius, whereas under the MAP rule (Eq 16) the agent selects only the next beam aim. Two sets of *P*_*M*_ and *P*_*FA*_ values are defined: one represents the agent’s internal model of sensory uncertainty (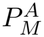 and 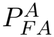), and the other specifies the true values for the sensor in the search space (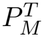 and 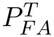). This setup allows investigation of how model performance varies when the agent’s sensory uncecrtinty model differs from true sensory statistics.

By default, the infotaxis model randomly selects the next beam aim from cells with the smallest expected entropy across the entire search space, whereas the MAP model randomly selects the next beam aim from cells with the highest posterior probability. This selection can be optionally restricted to only cells within a user-defined radius from the current beam aim to make the beam aim trajectory more realistic (smoother). All simulations presented in the main text were generated without beam aim movement restriction. See Fig A in S2 Appendix for search examples generated with beam aim movement restriction.

The agent terminates the search when one of two conditions is met: (1) the maximum posterior probability for any cell to contain the target 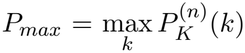 exceeds an arbitrary threshold *p*_*th*_ = 0.95; or (2) the change of *P*_*max*_ across three consecutive pings is smaller than 10^™5^, indicating convergence. A search is deemed successful if, upon termination, the cell with the highest posterior probability coincides with the true target location. Search performance is characterized primarily by the success rate and the number of pings required to complete a search, as well as *P*_*max*_, entropy, and the cumulative beam aim path length at the end of the search. These metrics are compared over 1000 randomized simulations, in which the true target location is randomly assigned for each run. The distributions of a given metric under any two model parameterizations are compared using the Mann-Whitney U tests (Conover, 1999). Although the search space size can be specified arbitrarily, *r*_*A*_ = 5 is used in most figures for clarity and ease of interpretation. Note that the specific values chosen for *p*_*th*_ and *P*_*max*_ convergence threshold do not influence the observed search performance trends (see Sec. III A 3).

The echolocation infotaxis search procedure is summarized in Algorithm 1.

#### Algorithm 1

Echolocation infotaxis search.

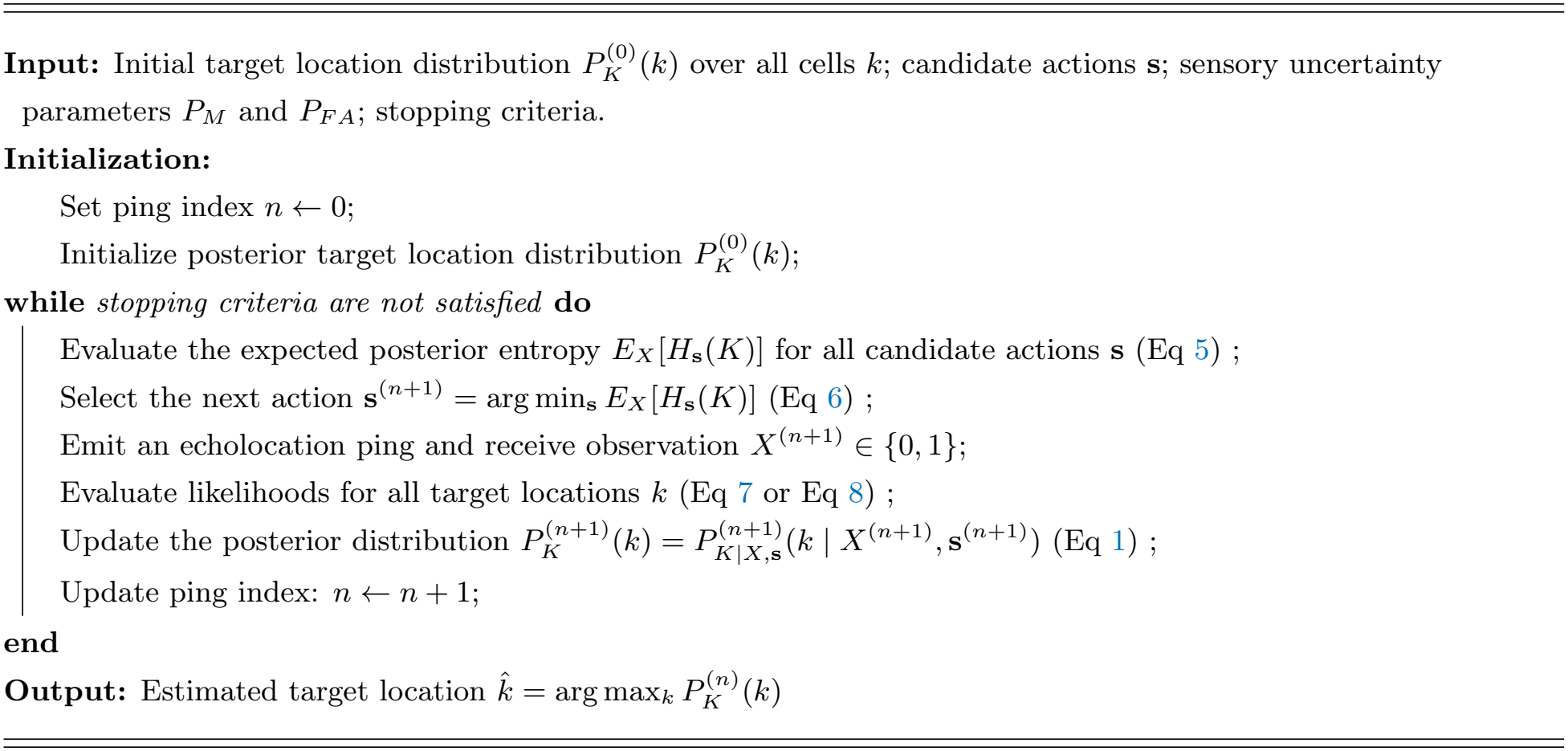

## III. RESULTS

Computational simulations of an active infotaxis agent performing single target search revealed a rich set of behaviors that provide insights into the properties of infotaxis in echolocation. In the following sections, we describe the general search dynamics, explore specific features of the search behavior, and compare the performance of infotaxis and MAP search through simulations.

### A. Infotaxis search behavior

#### 1. Typical search dynamics

Given a fixed beam footprint size and moderate *P*_*FA*_, the infotaxis agent randomly selects beam aims from interior cells of a search space at the beginning of a search (Fig 3A, ping 0). When the search space is moderately large relative to the beam footprint, the agent typically does not receive a target detection during the first few pings and would update the probability distribution of target location by placing higher probability in previously unexplored cells (Fig 3A, pings 1-7). This process continues until a target detection occurs, which sharply alters the distribution of target probability across the search space (Fig 3A, ping 8). The agent now concentrates its search within and around the footprint of the ping that yielded the detection, directing subsequent beam footprints to overlap maximally with the region of high target probability (Fig 3A, pings 9-11) to further narrow down the target location. The search terminates once the target probability of a single cell exceeds the stopping threshold *p*_*th*_ (Fig 3F).

**FIG 3:**
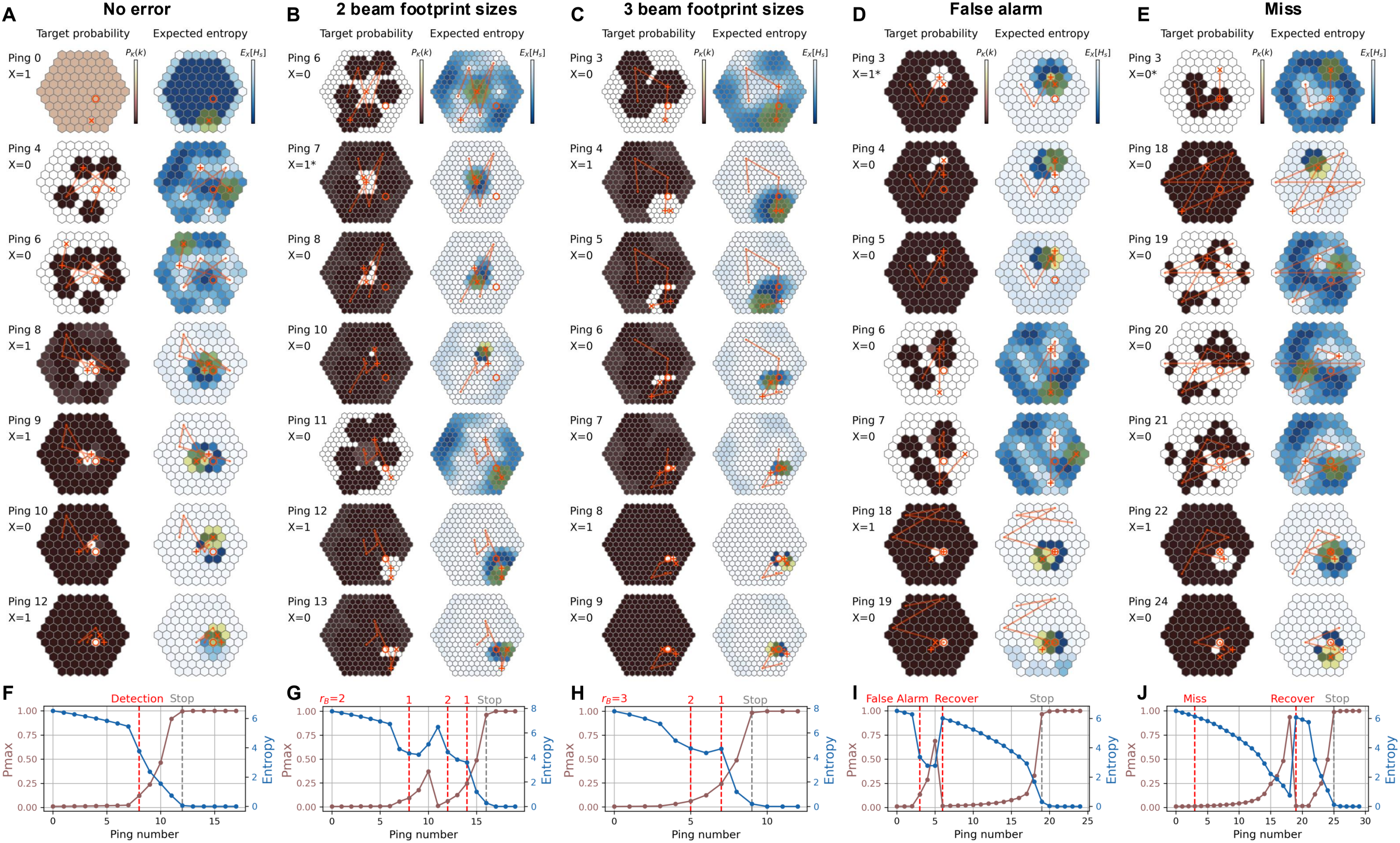
Search dynamics of an echolocating infotaxis agent. (A-E) Posterior distribution of target location (left column) and expected entropy (right column) after receiving echo observation of selected pings in representative searches. Panels A, D, E show searches with *r*_*B*_ = 1 with no detection errors, a false alarm, and a miss early on, respecitvely. Panels B-C show searches with *r*_*B*_ = {1, 2} and *r*_*B*_ = {1, 2, 3}, respectively. Red lines: beam aim trajectory over five pings, including the upcoming ping. Yellow shade: next beam footprint. Circle (◦): true target location. Cross (×): next beam aim. Plus (+): previous beam aim. Color represents a relative scale in the value range spanned in the search space. Under infotaxis, only the relative scale matters because the agent selects the next set of actions that gives the lowest expected entropy among all choices. Asterisk (*) in the ping number annotation indicates a false alarm or a miss. (F-J) Variation of maximum target probability in the search space (*P*_*max*_) and entropy across successive pings, annotated with events occurring at specific pings. Each search terminates when *P*_*max*_ ≥ *p*_*th*_ = 0.95. See Table A in S3 Appendix for all search parameter values, and S1–S5 Videos for the full sequences.

In such a typical search sequence, the initial stage before any target detection represents the “exploration” phase, during which the agent surveys the search space to identify a probable target area. Once a detection occurs, the search transitions into the “exploitation” phase, where the agent prioritizes refining the already identified highly probable target region. As in the classic odor source tracking scenario (Vergassola *et al*., 2007), an echolocating infotaxis agent naturally balances exploration and exploitation while maximizing the information gain at each ping.

The initial search behavior changes dramatically when the relative beam footprint coverage *α* is large or when *P*_*FA*_ is high. Under these conditions, the agent exhibits a distinct boundary- or corner-seeking behavior, often directing its beam toward the edges of the search space (Fig B in S2 Appendix). This occurs because such beam aims reduce the number of cells covered within the footprint, as part of the beam extends outside the search space. This effectively lowers the collective probability of false alarm from its value if the beam aim was in the middle of the grid (≈ *N*_*B*_*P*_*FA*_, see Eq 12), allowing the agent to be more confident about any target detection.

#### 2. Adjustment of beam footprint size

Given choices of beam footprint sizes, the infotaxis agent automatically switches between larger and smaller beam coverages to balance between exploration and exploitation (Fig 3B-3C and 3G-3H). Under moderate *P*_*FA*_, the agent consistently starts the search with the largest beam footprint to survey the search space for the target (Fig 3B and 3G, *r*_*B*_ = 2 for ping 0-6; Fig 3C and 3H, *r*_*B*_ = 3 for ping 0-3). Once a target detection occurs, the agent immediately switches to a smaller Infotaxis in echolocation beam footprint, trying to narrow down the target location within the highly probable target area. The agent’s response depends on the available beam size options. When given two choices, it switches to the smaller footprint (Fig 3B, *r*_*B*_ = 1 for ping 7). When given three choices, it first switches to the intermediate size and then to the smallest once most of the area covered by the ping yielding the first target detection has been explored (Fig 3C, *r*_*B*_ = 2 and *r*_*B*_ = 1 for ping 5 and 7, respectively).

The observed changes of beam footprint size are intuitive: A smaller footprint is more efficient in precisely locating the target within the confined high-probability area covered by the previous ping yielding a target detection. However, if this area is moderately large, selecting an intermediate beam size provides a better balance between exploration and exploitation for refining the target location. In addition, changes of beam footprint size are not unidirectional. In the case with two available beam sizes, once the agent recovers from a false alarm using several pings with the smaller footprint (Fig 3B and 3G, ping 8-11), it switches back to the larger beam footprint to restart exploring the search space (Fig 3B and 3G, ping 12).

#### 3. Impacts of sensory uncertainty

When misses or false alarms occur, the infotaxis agent recovers from these sensory errors in distinctive ways that impact the search efficiency. Following a false alarm (*X* = 1 despite the target being outside the footprint), the agent directs the next few pings to sequentially investigate cells covered by the beam footprint yielding the erroneous detection, and efficiently rules out the possibility that the target is located withing these cells (Fig 3D, ping 4-6). It follows intuitively that the recovery time after a false alarm depends on the size of the beam footprint. Because a target detection—whether true or spurious—causes a sharp contraction of highly probable target region, followed by a gradual redistribution of target probability, a false alarm appears as a distinct “dip” in the actual entropy curve of a search (Fig 3I, between ping 2-6).

In contrast, the infotaxis agent is affected more severely by a miss, often for a longer duration. When a miss occurs (*X* = 0 despite the target being within the beam footprint), the agent assigns low target probability to the cells covered by the beam and moves away from the area, as revisiting it is deemed less informative (Fig 3E, ping 3, low expected entropy around the beam footprint). The agent typically revisits the missed region only after all other parts of the search space have been explored without success (Fig 3E, pings 4-19). Therefore, when a miss occurs early in a search, the resulting entropy curve resembles that of searches with a long exploration phase (Fig 3J). Once the agent realizes that a miss has occurred after failing to detect the target elsewhere, the entropy rises abruptly to the level observed before the miss, reflecting renewed uncertainty and resumption of exploration (Fig 3E and 3J, ping 19; panels A and C of Fig C in S2 Appendix, ping 24). When a miss occurs later in a search, after the high-probability target region has already become spatially confined, it produces only a transient rise in the entropy curve (panels B and D of Fig C in S2 Appendix).

Overall, infotaxis search performance degrades with increasing *P*_*M*_ and *P*_*FA*_, with decreasing success rate and increasing number of pings required to complete a search (Fig 4, with *p*_*th*_ = 0.95, and Fig D in S2 Appendix, with *p*_*th*_ = 0.99). For instance, when *r*_*A*_ = 5 and *r*_*B*_ = 1 (Fig 4A), the median number of pings required to complete a search increases from 10 to 20 as *P*_*M*_ and *P*_*FA*_ rise from 0.001 to 0.05. Under the same conditions but with a larger beam (*r*_*B*_ = 2; Fig 4B), the median increases from 7 to 29. Comparisons between simulations with both *P*_*M*_ and *P*_*FA*_ elevated and those with only *P*_*FA*_ elevated (Fig 4 vs. Fig E in S2 Appendix) indicate that the reduced efficiency primarily stems from the need to mitigate false alarms (observe the systematic increase of ping number). In contrast, comparisons with simulations with only *P*_*M*_ elevated (Fig 4 vs. Fig F in S2 Appendix) show that high *P*_*M*_ mainly increases the number of outlier searches that require exceptionally many pings to complete, likely due to misses occurring early in the search (observe the small number of runs with very high ping number). In addition, across the range of *P*_*M*_ and *P*_*FA*_ investigated, the maximum posterior at the end of the search remains high (Fig G in S2 Appendix), and the entropy of the posterior increases only slightly with higher sensory uncertainty (Fig H in S2 Appendix). Interestingly, the cumulative beam aim path length until search termination remains stable when *r*_*B*_ = 1, but increases substantially when *r*_*B*_ = 2 and sensory uncertainty is high (Fig I in S2 Appendix). This is likely due to the extra pings needed to narrow down the target location within a larger beam footprint, which can be further lengthened if false alarms or misses occur.

**FIG 4:**
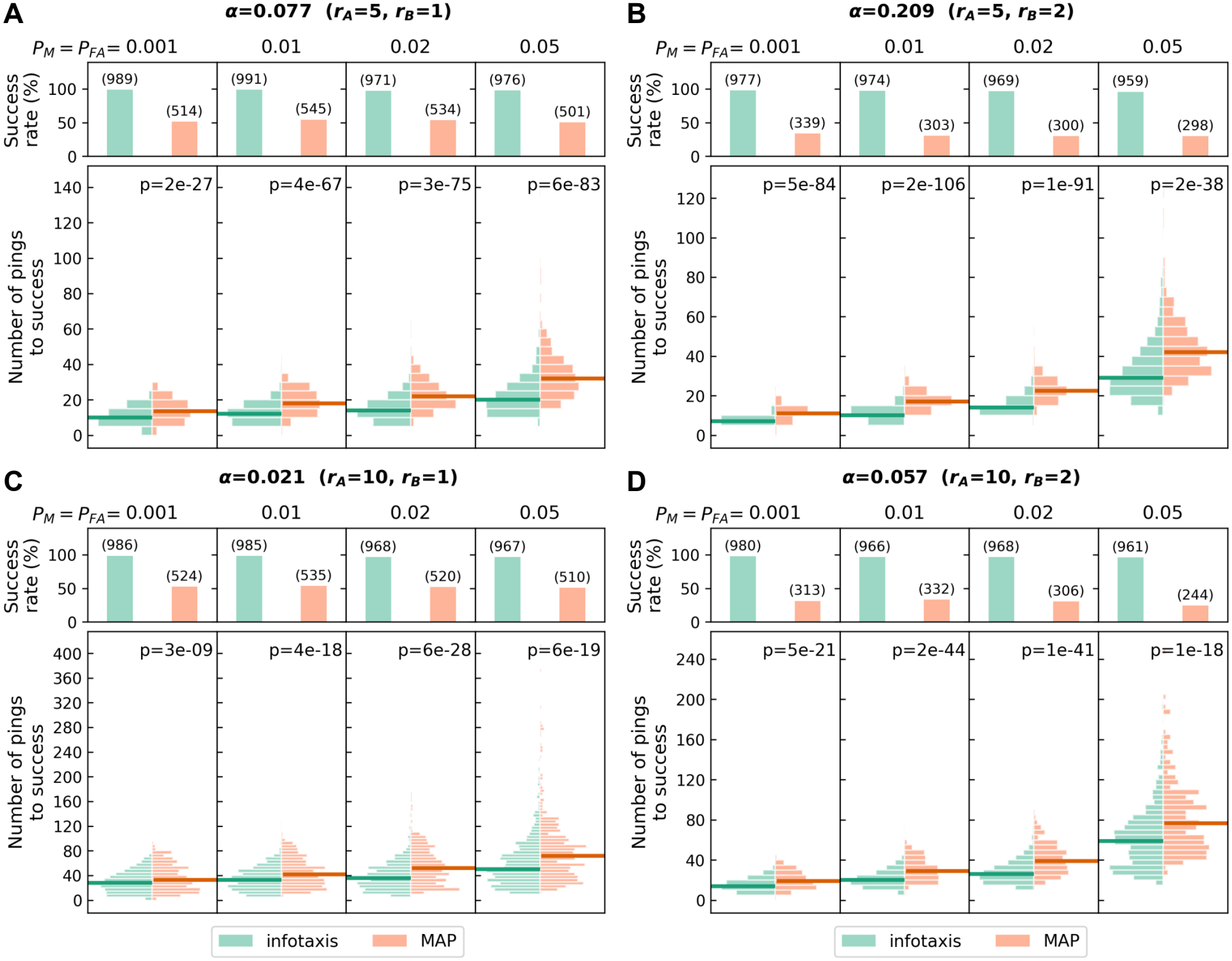
Performance comparison of infotaxis and MAP searches. Success rate (top) and number of echolocation pings required (bottom) to locate the target for infotaxis (green on the left) and MAP (orange on the right) searches under different parameter combinations. The infotaxis agent consistently outperforms the MAP agent, achieving a higher success rate with fewer pings. The numbers of successful runs out of 1000 runs are indicated in the parentheses. A search is successful when the cell with the highest posterior probability at search termination coincides with the true target location. *r*_*A*_ and *r*_*B*_ denote the radii of the search space and the beam footprint, respectively, and *α* is the relative size of the beam footprint to the search space. The reported *p*-values are from Mann-Whitney U tests under the alternative hypothesis that the infotaxis search requires fewer pings to complete than the MAP search. Note that the beam footprint radius *r*_*B*_ is fixed for both the infotaxis and MAP agents. See Sec. II D for definitions of all quantities and details on the stopping condition.

When the infotaxis agent’s assumed sensory uncertainty (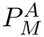 and 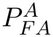) mismatches the true sensory statistics of the search space (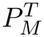 and 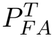), its search performance declines in different ways depending on whether sensory uncertainty is overestimated or underestimated. When the agent underestimates the sensory uncertainty, the localization success rate declines sharply as the mismatch widens (Fig 5, purple bars on all top panels). Detailed investigation revealed that the agent often mistakenly takes repeated false alarms as true detections because of the assumed lower sensory uncertainty, causing it to prematurely terminate the search at an incorrect target location (Fig J in S2 Appendix). Paradoxically, the number of pings required to complete a search decreases with increasing mismatch (Fig 5, purple histograms on all bottom panels). This is because when the agent happens to encounter the target, its underestimated sensory uncertainty causes the target probability distribution to converge much faster than under the true (higher) sensory uncertainty. Across the range of sensory uncertainty investigated, *P*_*max*_ and entropy at the end of the search are mostly comparable between the matched and underestimated cases for successful runs (Figs K and L in S2 Appendix). The cumulative beam aim path length decreases with increasing true sensory uncertainty for the underestimated cases (Fig M in S2 Appendix). However, note the rapidly dropping success rate.

**FIG 5:**
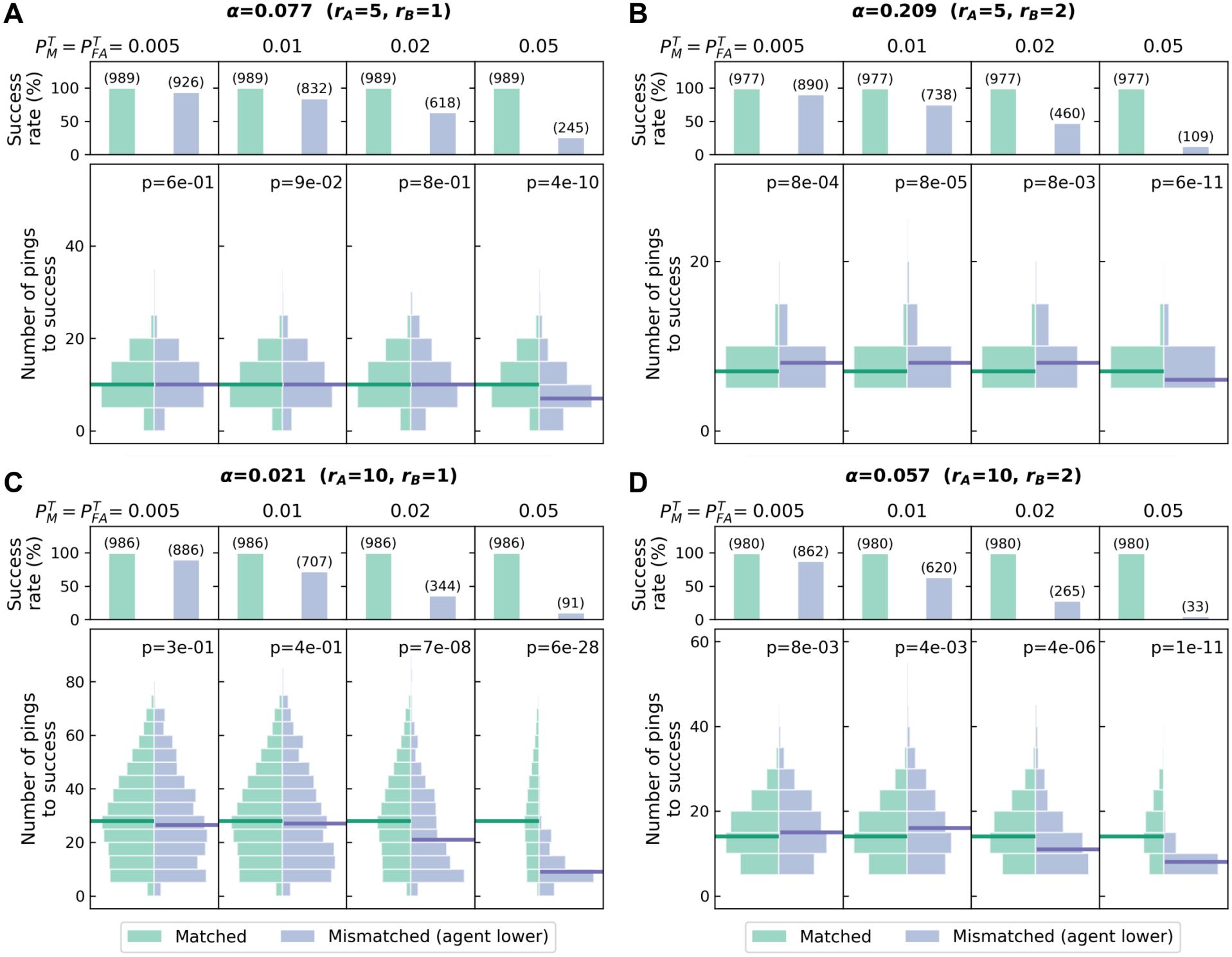
Impacts on infotaxis search performance when the agent underestimates sensory uncertainty. Success rate (top) and number of pings required (bottom) to locate the target when the agent’s assumed *P*_*M*_ and *P*_*FA*_ match (green on the left) and mismatch (purple on the right) the true values. Here, the agent assumes 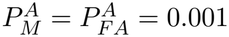, but the true values 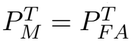 vary across a range of values indicated on the top of each panel. The number of successful runs out of 1000 runs are indicated in the parentheses. A search is successful when the cell with the highest posterior probability at search termination coincides with the true target location. Note that the success rate drops sharply as the mismatch increases, even though the searches may terminate earlier. The reported *p*-values are from Mann-Whitney U tests under the alternative hypothesis that the number of pings required to locate the target are the same under the matched and mismatched conditions. See Sec. II D for definitions of all quantities and details on the stopping condition.

When the agent overestimates sensory uncertainty, both the localization success rate and the number of pings required to complete a search increase (Fig 6, purple bars and histograms on all top and bottom panels). This is intuitive, as with higher 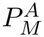 and 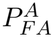 each observation produces a smaller update to *P*_*K*_(*k*), which lengthens the search. Due to the slower posterior updates in these cases, while *P*_*max*_ at the end of the search remains high, the values are lower than those in the matched case (Fig N in S2 Appendix), with correspondingly higher entropy (Fig O in S2 Appendix). Note that these slightly higher entropy values are still much lower than those under most of the MAP searches (Fig H in S2 Appendix, also see Sec. III D). The cumulative beam aim path length is higher with higher assumed sensory statistics and when *r*_*B*_ = 2 (Fig P in S2 Appendix), due to the extra pings needed to narrow down the target location within a larger beam footprint, especially when false alarms and misses are more likely to occur.

**FIG 6:**
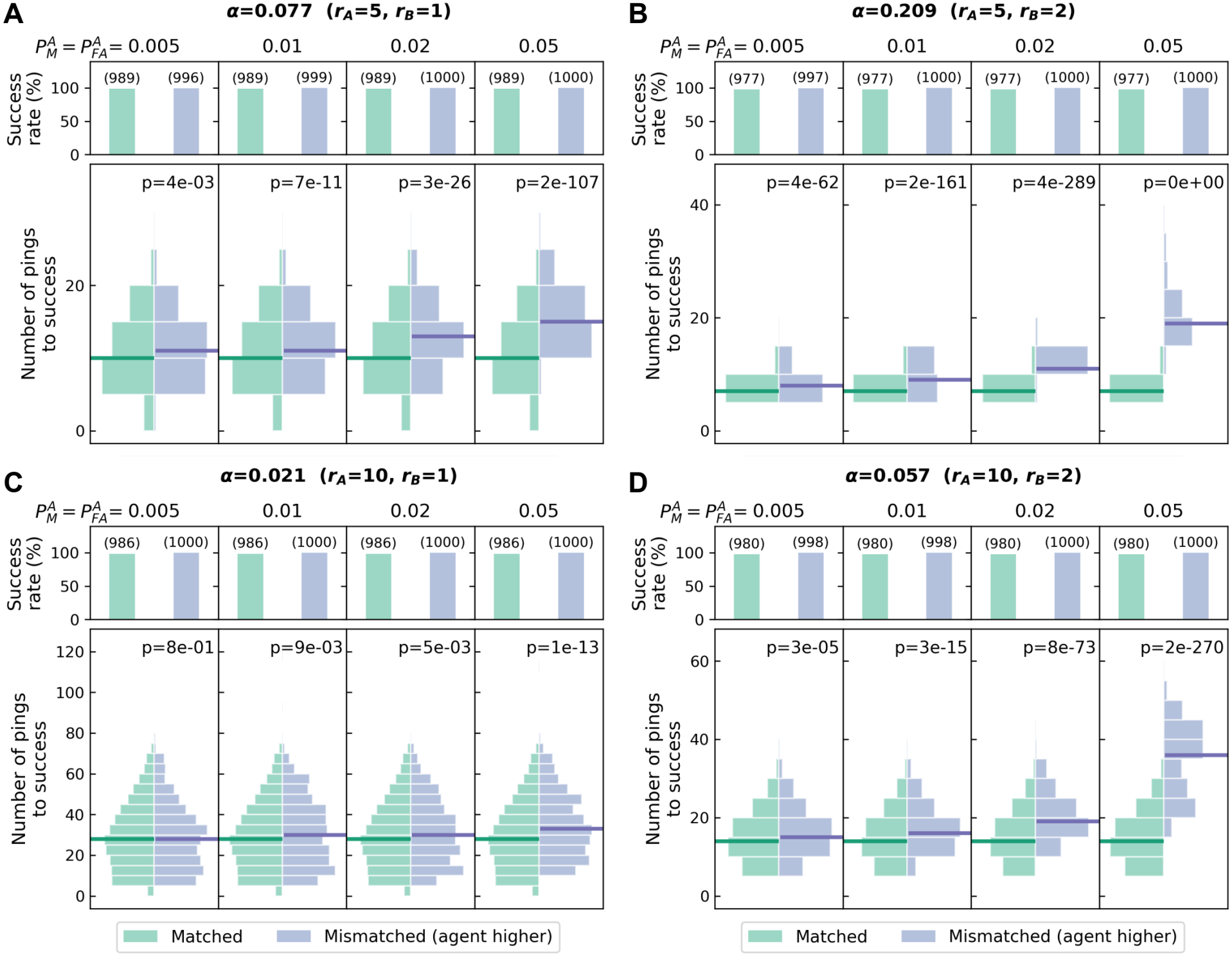
Impacts on infotaxis search performance when the agent overestimates sensory uncertainty. All plot and annotation details are identical to those in Fig 5, except for here the true sensory statistics are fixed at 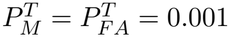 but the agent’s assumed 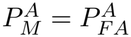 varies across each panel. Both the number of pings required to locate the target and the success rate increase with increasing mismatch, as the agent becomes more cautious.

### B. Entropy decreases in the exploration phase

During the exploration phase before the first target detection, the entropy decreases mono- tonically as the infotaxis agent rules out regions not containing the target (Fig 3F, ping 1-7). Intuitively, the beam footprint size relative to the search space (*α*) and sensory uncertainty (*P*_*M*_ and *P*_*FA*_) would influence this update. Below, we explore the connection between these parameters and the trend of entropy decrease up until the first target detection *X* = 1.

Assume 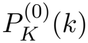 is uniform over the search space in the beginning of a search, and let 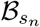 denote the set of cells covered by the beam footprint of the *n*th ping. Using Eq 1, the updated scene after observing *X* = 0 with ping *n* + 1 is

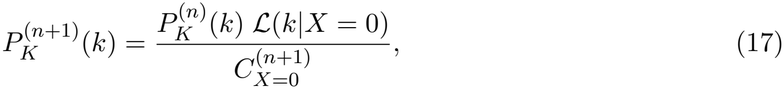

where *ℒ*(*k*|*X* = 0) follows Eq 9 or Eq 10 depending on if cell *k* lies within 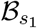, and 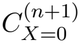 is the normalization factor determined by all values in the updated map *P* 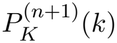

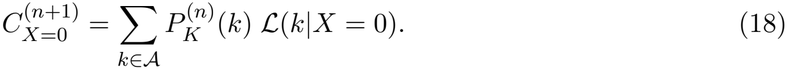

After ping 1, when no target is detected,

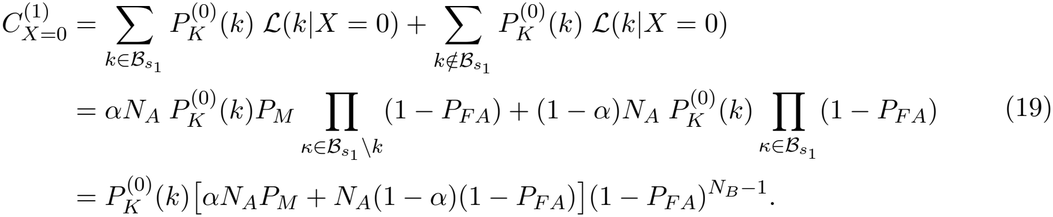

Recall that *α* = *N*_*B*_*/N*_*A*_ represents the relative size of the beam footprint to the search space. Substituting Eqs 9 and 10 for the likelihood in the numerator of Eq 17 makes it clear that the 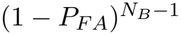 in 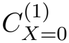 cancels in both cases, leaving

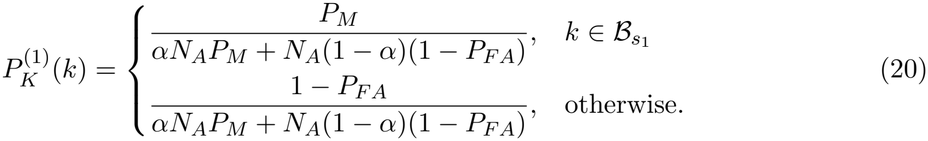

Using the approximation 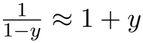, where 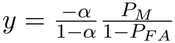, we find

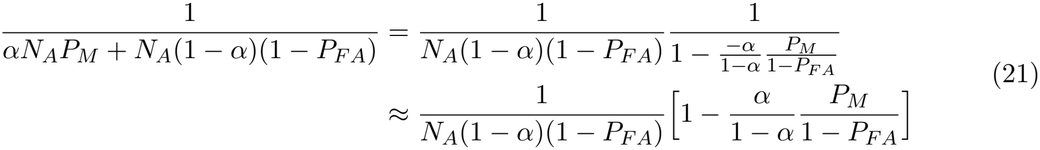

and obtain

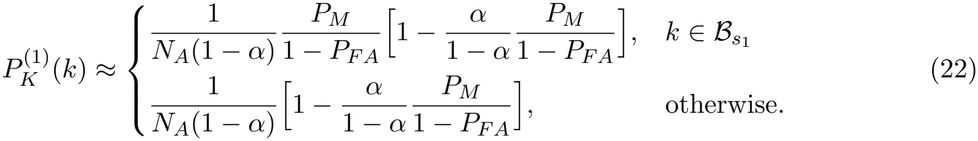

Here, cells within 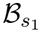 are updated with low target probabilities, especially when *P*_*M*_ is low. In fact, when *P*_*M*_ → 0, the probability of target is uniform across the unobserved cells at 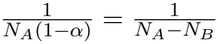.

Intuitively, since the agent has already gathered information about cells covered by the beam footprint, the next beam aim would be directed such that the new footprint does not overlap with the previous one (Fig 7A). Therefore,

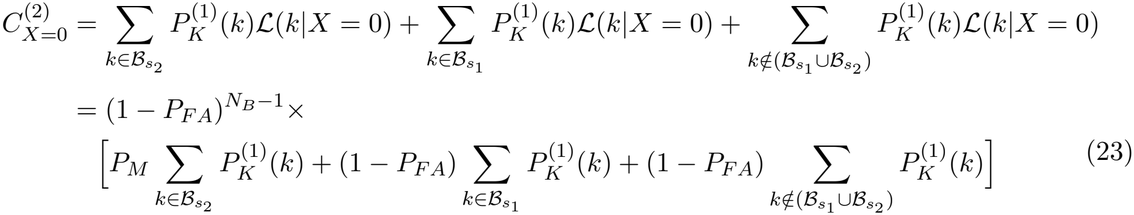

**FIG 7:**
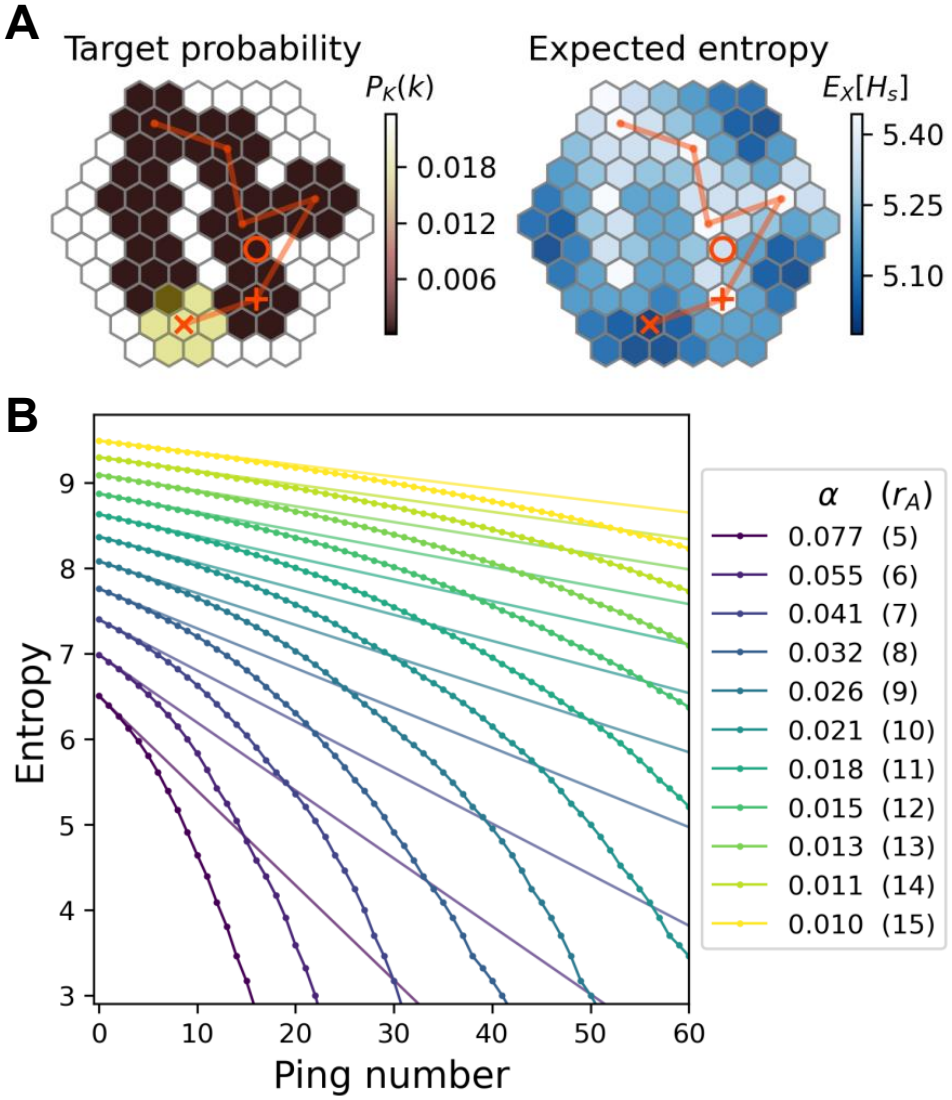
Entropy variation during infotaxis searches. (A) Effects of consecutive non-overlapping pings with no target detection (*X* = 0) on the probability distribution of target location (left) and expected entropy (right). See Fig 3 caption for definitions of all symbols. *P*_*M*_ = *P*_*FA*_ = 0.005 in this example. (B) The slope of entropy decrease varies systematically depending on the relative size of the beam footprint to the search space (*α*). Lines with markers show actual entropy decrease from simulations, and lines without markers show the analytical approximation (Eq 28). Numbers in the parenthesis indicate the search space radius *r*_*A*_. The beam footprint radius is fixed at *r*_*B*_ = 1. See Table A in S3 Appendix for all search parameter values.

Again, the 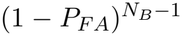 terms in the likelihood and normalization terms of Eq 17 cancel, and using the same approximation in Eq 21, we obtain

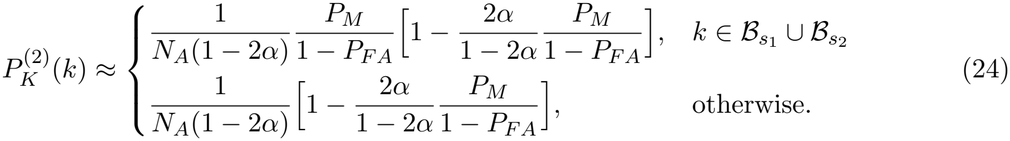

Following the same reasoning, when no target is detected, the agent progressively samples unvisited regions of the search space, moving the beam in a manner that maximizes coverage while minimizing redundant sampling of previously interrogated areas (Fig 7A). We find that,

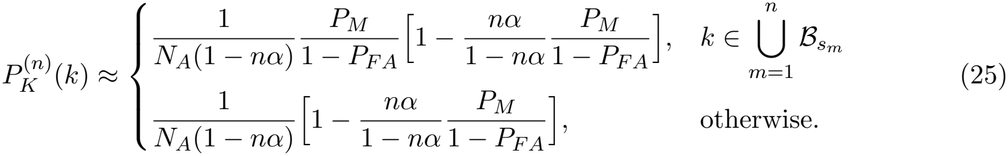

This approximated form of 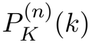 is intuitive: In the limit *P*_*M*_ → 0, when no target detection is received across multiple pings, the probability of target present in cells covered by all preceding non-overlapping beam footprints would be redistributed to unexplored cells in the search space:

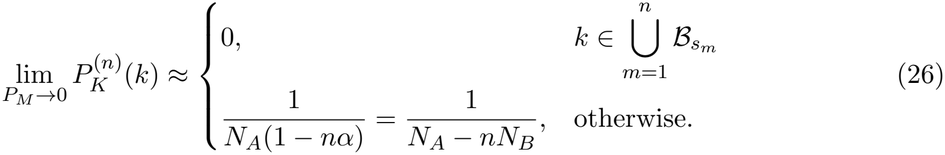

The accuracy of these approximations deteriorates with increasing *α* and *P*_*M*_, as the approximation in Eq 21 becomes less valid.

When both *P*_*M*_ and *α* are small, let 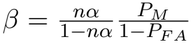, we can then show that the entropy after the *n*th ping,

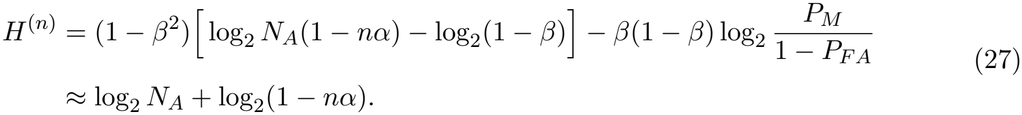

Using log_2_ *y* = ln *y/* ln 2 and ln(1 − *y*) ≈ ™*y*, we obtain the approximate slope of entropy decrease over consecutive pings,

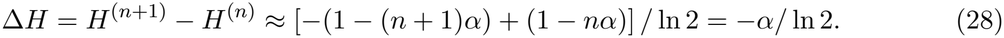

A detailed derivation of the above is provided in Section B in S1 Appendix.

Eqs 25-28 suggest that the rate at which entropy decreases before the first target detection depends strongly on *α*, as shown in Fig 7B. Notably, the actual trajectory of entropy decrease deviates from the analytical approximate when *α* is large, both because the approximation becomes less accurate and because as the number of pings increases, subsequent beam footprints would either lie partially outside of the search space or overlap with previous footprints due to the finite search space, which can be observed in Fig 7A.

### C. Beam aim selection after the first target detection

An interesting observation from the simulations is that once a target detection occurs, the agent may either repeats aiming its beam at the same center cell or shift the beam aim to a neighboring cell such that the new beam footprint partially overlaps with the previous one (Fig 8A). Repeating the same beam aim serves as a confirmation: if the subsequent ping does not yield another detection, the previous detection was likely a false alarm; if it does, the target is likely located within this beam footprint. Alternatively, shifting the beam aim to a neighboring cell allows the agent to further narrow down the target location: if the next ping yields a detection, the target is likely located within the intersection of both beam footprints; if no detection occurs on the second ping, the target is likely within the cells of the first footprint not shared with the second.

**FIG 8:**
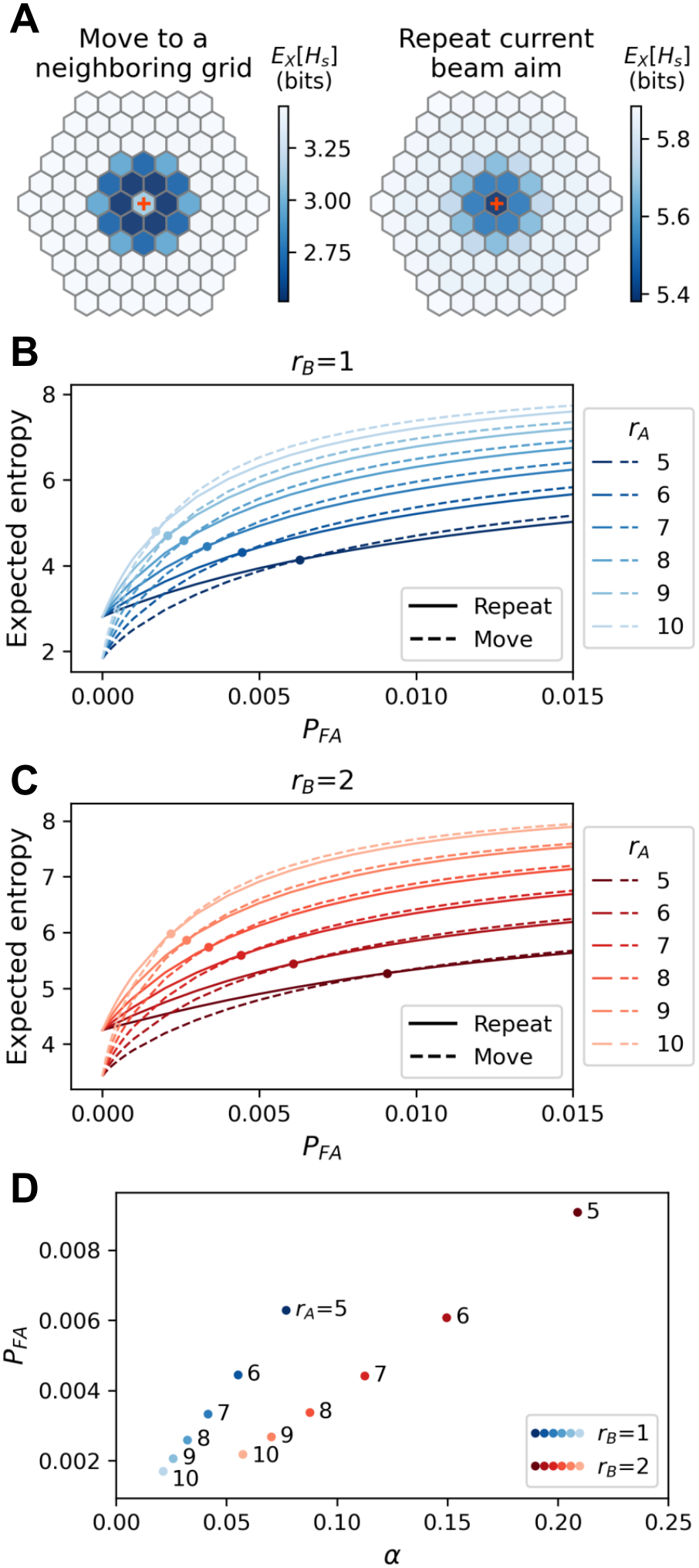
Selection of the next beam aim following a target detection. (A) Expected entropy after detecting a target echo. The agent may choose to move the beam aim to a neighboring cell (left; *P*_*FA*_ = 0.001) or repeat the current beam aim (right; *P*_*FA*_ = 0.02). Plus (+): previous beam aim. (B-C) Changes in expected entropy for shifting the beam aim to a neighboring cell (dashed line) vs. repeating the current beam aim (solid lines) as a function of *P*_*FA*_ for two beam footprint radii across a range of search space sizes. (D) Values of *P*_*FA*_ at which the agent switches from shifting the next beam aim to a neighboring cell to repeating the current beam aim, plotted as a function of *α*. All values and curves shown are computed from the exact expressions numerically rather than using approximate analytical expressions. See Sec. II D for definitions of all quantities.

To better understand the parameters that drive this decision, we derive 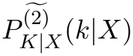 and *P*_*X*|**s**_(*X*|**s**), the key quantities that determine the expected entropy *E*_*X*_[*H*_**s**_(*K*)] after the first target detection, when the agent needs to decide whether to repeat the same beam aim or to shift the beam aim to a neighboring cell. The tilde 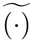 in the superscripts distinguishes this “local” ping indexing, which starts from the first target detection, from the “global” ping sequence of the entire search.

To reduce clutter and preserve the flow in the main text, the full derivations are presented in Section C in S1 Appendix and only the main derivation steps and results are summarized below: The derivation begins with the target location distribution 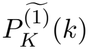 after receiving the first target detection (Section C.1 in S1 Appendix). We then derive the updated target location distribution 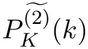 after the immediately subsequent ping for two cases: when the agent repeats the same beam aim (Section C.2 in S1 Appendix) and when it shifts the beam aim to a neighboring cell (Section C.3 in S1 Appendix). These posterior distributions are then used to derive *P*_*X*|**s**_(*X*|**s**).

When the agent repeats the same beam aim 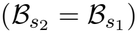, the normalization factors (see Eq 17) of the updated target probability map 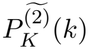 with echo observation *X* = 0 and *X* = 1 are

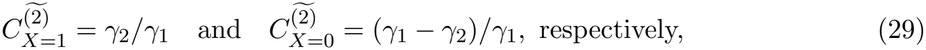

where

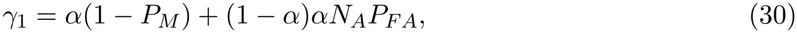

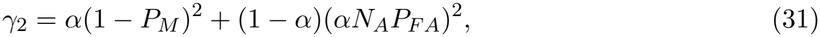

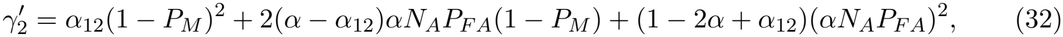

in which *α*_12_ denotes the proportion of the overlapping area between the first and the second beam footprints relative to the entire search space.

Using Eq 1 and the binomial approximation (1 + *y*)^*α*^ ≈ 1 + *αy*, we obtain

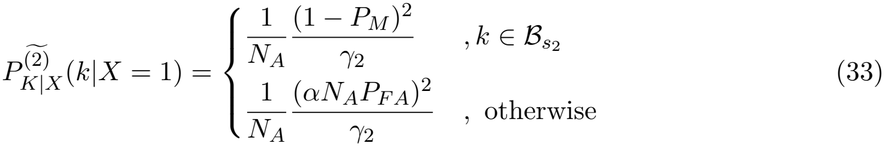

or

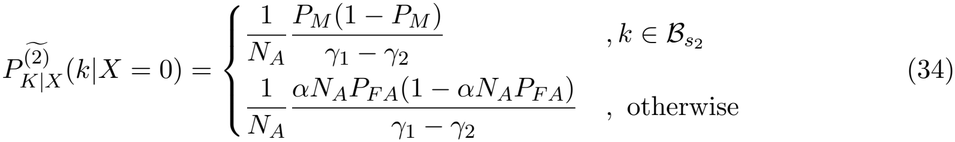

depending on whether a target detection occurs after the subsequent ping. Using Eq 13, the resulting probability of receiving a target detection in the subsequent ping is

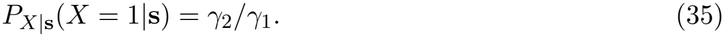

When the agent shifts the next beam aim to a neighboring cell 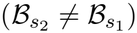,

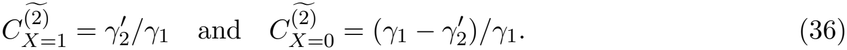

The updated target probability distribution is

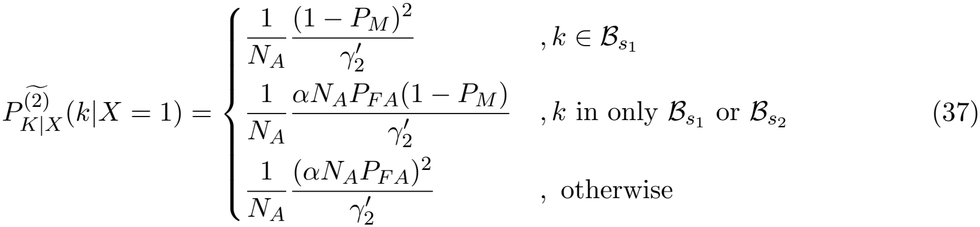

or

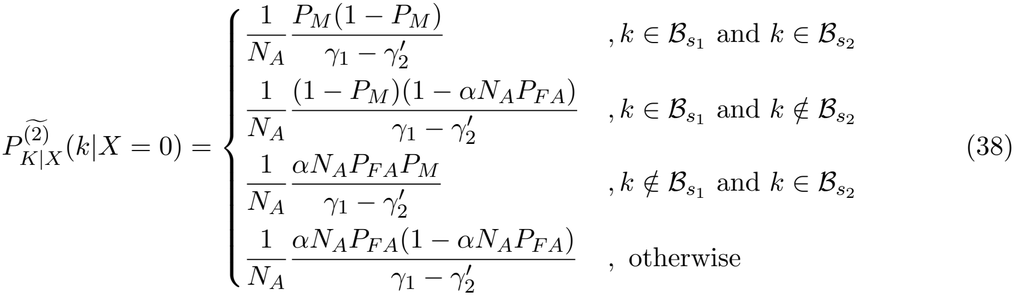

depending on if a target detection occurs after the subsequent ping. Similarly, following Eq 13, the probability of detecting a target in the subsequent ping is

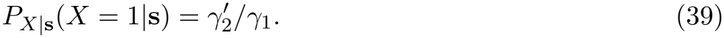

The expected entropy after the subsequent ping, *E*_*X*_[*H*_*s*_(*K*)], can be evaluated from *P*_*X*|**s**_(*X*|**s**) and *H*_**s**_(*K*|*X*) via Eqs 4-5, where *H*_**s**_(*K*|*X*) can be obtained from the updated target probability distribution 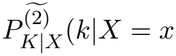. Examination of these quantities reveals that changes in *P*_*FA*_, through the term *αN*_*A*_*P*_*FA*_, influence the probability of detecting a target, and thus the expected entropy, far more strongly than changes in *P*_*M*_. Specifically, 1 − *P*_*M*_ and (1 − *P*_*M*_)^2^ vary only slightly with varying *P*_*M*_, but *αN*_*A*_*P*_*FA*_ and (*αN*_*A*_*P*_*FA*_)^2^ vary strongly with varying *P*_*FA*_ as well as changes of *α* and *N*_*A*_. As an example, Table I shows that a tenfold increase in *P*_*FA*_ substantially alters all quantities contributing to *E*_*X*_[*H*_**s**_(*K*)], whereas a comparable change in *P*_*M*_ produces only minor effects.

Through *αN*_*A*_*P*_*FA*_, the combined factor *αN*_*A*_ (= *N*_*B*_, the beam footprint size) controls the overall trend of expected entropy changes, whereas changes in *P*_*FA*_ produce more gradual variations. Specifically, the smallest increment of the beam footprint radius from *r*_*B*_ = 1 to *r*_*B*_ = 2 nearly triples *N*_*B*_ from 7 to 19, while a large change in background noise would be needed to triple *P*_*FA*_. In Fig 8B-C, the expected entropy curves for repeating the same beam aim (solid lines) vs shifting it to a neighboring cell (dashed lines) cross over at different *P*_*FA*_ values for the two beam footprint sizes. Changes in beam footprint size cause categorical shifts in these curves and, consequently, alter the crossover points. For a given beam footprint, increasing *P*_*FA*_ causes the infotaxis strategy to switch from shifting beam aim to a neighboring cell (lower expected entropy for dashed lines) to repeating the current aim (lower expected entropy for solid lines). This behavior is intuitive: when *P*_*FA*_ is low, the agent is confident that a detection reflects the true target and continues to refine its location; whereas when *P*_*FA*_ is high, verifying the detection by repeating the same aim becomes the safer choice.

**TABLE I:**
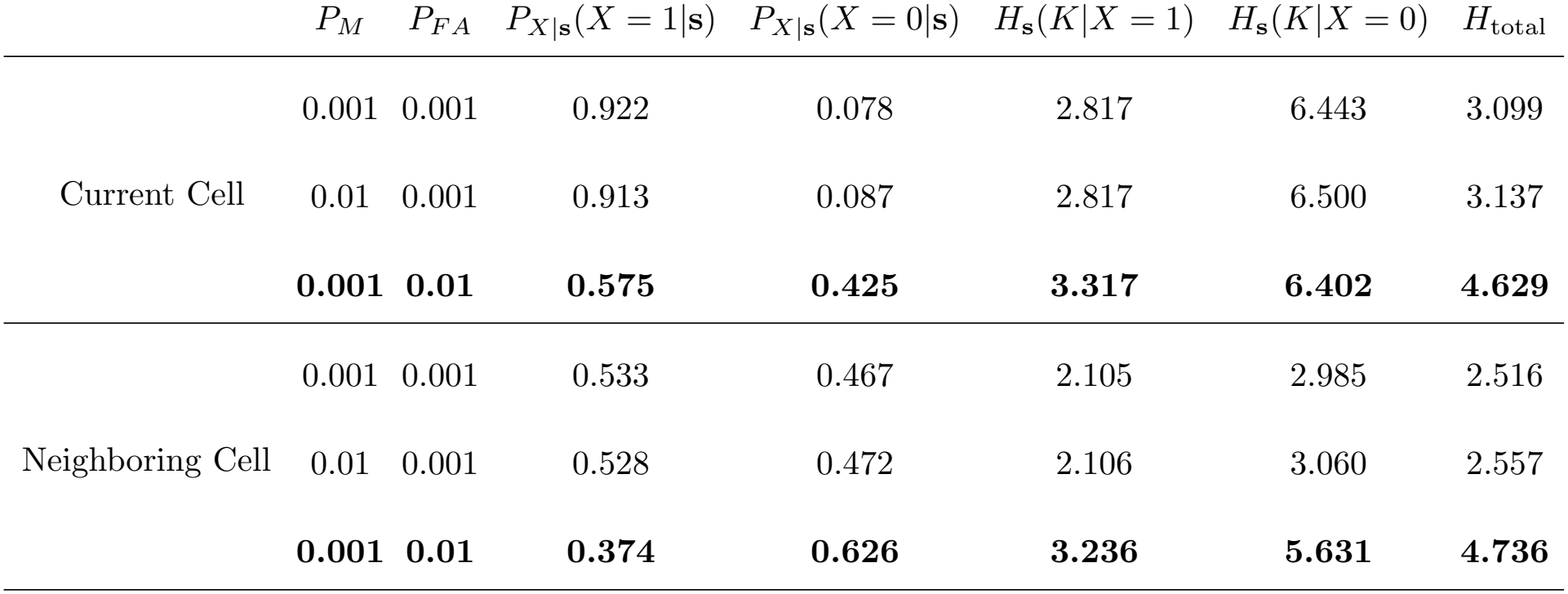
Influence of *P*_*M*_ and *P*_*FA*_ on quantities contributing to the expected entropy. *E*_*X*_[*H*_**s**_(*K*)]. *P*_*X*|**s**_(·|**s**) and *H*_**s**_(*K*|·) are the probability of receiving echo outcome *X* from the next transmission and the entropy of the updated scene given the next beam configuration **s**. “Current Cell” indicates the scenario when the next beam aim remains in the same cell. “Neighboring Cell” indicates the scenario when the next beam aim shifts to a neighboring cell. The calculation here is based on the setup shown in Fig 2 in which *N*_*A*_ = 91 and *N*_*B*_ = 7, resulting in *α* = 0.077. All values are computed from the exact expressions numerically rather than using approximate analytical expressions.

|  | P<sub>M</sub> | P<sub>FA</sub> | P<sub>X\|**s**</sub>(X = 1\|**s**) | P<sub>X\|**s**</sub>(X = 0\|**s**) | H<sub>**s**</sub>(K\|X = 1) | H<sub>**s**</sub>(K\|X = 0) | H<sub>total</sub> |
|---|---|---|---|---|---|---|---|
| Current Cell | 0.001 | 0.001 | 0.922 | 0.078 | 2.817 | 6.443 | 3.099 |
|  | 0.01 | 0.001 | 0.913 | 0.087 | 2.817 | 6.500 | 3.137 |
|  | **0.001** | **0.01** | **0.575** | **0.425** | **3.317** | **6.402** | **4.629** |
| Neighboring Cell | 0.001 | 0.001 | 0.533 | 0.467 | 2.105 | 2.985 | 2.516 |
|  | 0.01 | 0.001 | 0.528 | 0.472 | 2.106 | 3.060 | 2.557 |
|  | **0.001** | **0.01** | **0.374** | **0.626** | **3.236** | **5.631** | **4.736** |

For a fixed search space size *N*_*A*_ (same *r*_*A*_ in Fig 8D), the expected entropy curves cross over at higher *P*_*FA*_ when the relative beam footprint size *α* is larger (larger *r*_*B*_). This is because when a larger portion of the search space is covered by the beam, the agent is more confident that the target lies within the footprint and can proceed to localize it more precisely. Given the same *α*, however, the larger the search space (higher *r*_*A*_ in Fig 8D), the lower the crossover *P*_*FA*_ values. This is because the larger the absolute beam footprint (number of cells covered), the higher the probability that any one of the covered cells would produce a false alarm, making it more advantageous for the agent to repeat the same aim to verify target presence.

Taken together, these findings show that *P*_*FA*_ is a key factor influencing the balance between exploration and exploitation under infotaxis in echolocation. This sensitivity underscores how prob- abilistic representations of sensory uncertainty directly shape the agent’s adaptive search strategies.

### D. Comparison between infotaxis and MAP searches

An infotaxis agent selects the next beam aim by maximizing the expected information gain. In contrast, a MAP agent bases its choice solely on 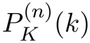 (Eq 16), which summarizes information from all previous echoes. While straightforward to implement, this simple decision rule leads to less effective and less efficient searches: MAP searches often fail to converge to a single cell containing the target (Figs 9 and 4, orange bars on all top panels) and, when successful, generally require more pings to complete (Fig 4, orange histograms on all bottom panels). Below we discuss specific features of MAP search behavior that give rise to these differences.

**FIG 9:**
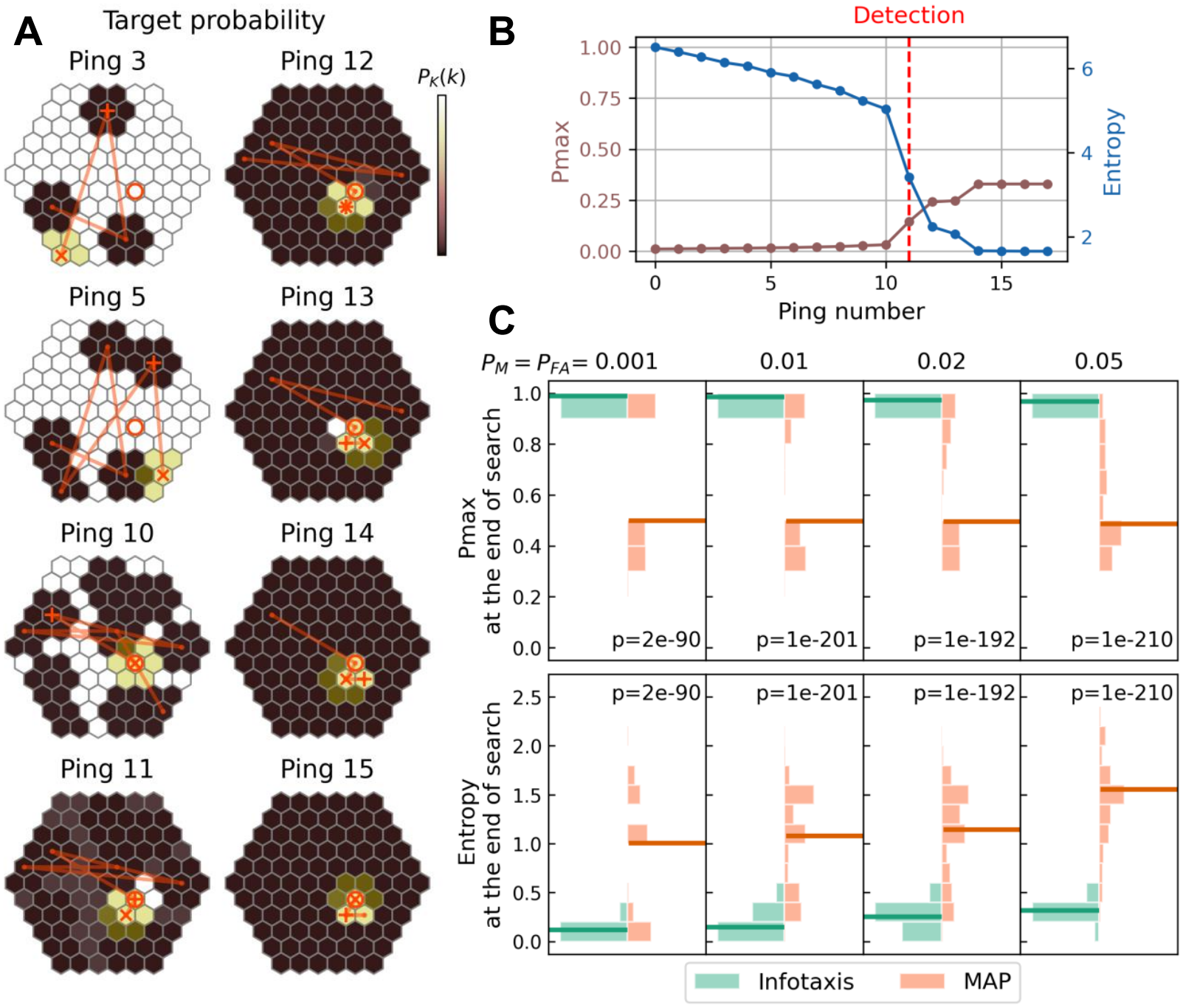
Search dynamics of an echolocating MAP agent. (A) Posterior distribution of target location after receiving echo observations from selected pings. Color represents a relative scale in the value range spanned within the search space. Under MAP, only the relative scale matters because the agent selects the next beam aim from the cells containing the highest posterior probability of target. See Fig 3 caption for definitions of all symbols, Table A in S3 Appendix for all search parameter values, and S6 Video for the full sequence. (B) Variation of maximum target probability in the search space (*P*_*max*_) and entropy across successive pings. In this search, the MAP agent failed to identify a single cell containing the target, and instead converged to three equally probable candidate cells at the end. (C) Distributions of *P*_*max*_ and entropy at the end of the search for infotaxis and MAP agents for a range of sensory uncertainty. For the MAP agent, the ending *P*_*max*_ and entropy depends strongly on the beam footprint size. Note that the beam footprint radius *r*_*B*_ is fixed (= 1) for both the infotaxis and MAP agents, and the search space radius *r*_*A*_ = 5. See Sec. II D for definitions of all quantities and details on the stopping condition.

During the exploration phase prior to the first detection, the MAP agent selects the next beam aim from all unexplored cells, including cells immediately adjacent to those already covered by previous pings (e.g., Fig 9A, ping 6). An infotaxis agent would avoid such choices under moderate *P*_*FA*_, as partially overlapping the new beam footprint with previously explored regions yields less new information.

Once a target detection occurs, the MAP agent consistently selects the next beam aim among cells *within* the region of highest target probability, which are cells covered by the ping yielding the detection but not yet covered by earlier pings (Fig 9A, ping 12 onward). In contrast, the infotaxis agent may aim *outside* this region, moving the beam footprint to further narrow down the target location (Fig 3A, ping 9 onward). As a result, the MAP agent may not be able to identify where the target is *not* located within the high probability region and get “stuck” with several equally probable candidate cells (Fig 9A, ping 14 onward), rendering the search unsuccessful. The larger the beam footprint, the more severe this problem. When such scenario arises, *P*_*max*_ is typically much lower than 1 (Fig 9C top panel and Fig G in S2 Appendix), with the entropy being correspondingly higher than those of successful searches (Fig 9C bottom panel and Fig H in S2 Appendix).

In the face of sensory errors, while both false alarms and misses increase the number of pings required to complete a search, infotaxis consistently outperforms MAP, with the performance gap widening with increasing *P*_*M*_ and *P*_*FA*_ and increasing beam footprint size (Fig 4). When *r*_*A*_ = 5 and *r*_*B*_ = 1 (Fig 4A), the median number of pings rises from 10 vs. 13.5 at *P*_*M*_ = *P*_*FA*_ = 0.001 to 20 vs. 32 pings at *P*_*M*_ = *P*_*FA*_ = 0.05 for the infotaxis and MAP agents, respectively. When the search space is larger (*r*_*A*_ = 10, *r*_*B*_ = 1; Fig 4C), the median increases from 28 vs. 33 pings at *P*_*M*_ = *P*_*FA*_ = 0.001 to 50 vs. 72 pings at *P*_*M*_ = *P*_*FA*_ = 0.05.

When a false alarm occurs, the MAP agent may occasionally select a subsequent beam aim whose footprint substantially overlaps with that of the false alarm, but such selections are random and would become more unlikely as the beam footprint increases. In contrast, the infotaxis agent would systematically revisit and rule out the false alarm region within a few pings (Fig 3D and I). When a miss occurs, even though both the MAP and infotaxis agents must explore the entire search space before recovering from the error, the infotaxis agent accomplishes this exploration more efficiently under its information-guided strategy.

When the agent underestimates or overestimates the sensory uncertainty, MAP remains less robust and less efficient than infotaxis (Figs Q and R in S2 Appendix), despite both agents experi- encing similar performance degradation. This difference is likely due to the MAP agent’s inherent inefficiency in beam aim selection and its frequent inability to further refine the target location once multiple candidate cells remain, as discussed above.

However, because infotaxis requires evaluating the expected information gain for *all* possible subsequent action choices, its decision making process is substantially more computationally de- manding than that of MAP (Fig S in S2 Appendix). Our simulations further showed that this speed difference widens with increasing search space size, as both the number of possible beam aim locations and the computations required to evaluate Eq 5 scale with the search space.

## IV. DISCUSSION

In this paper, we extended infotaxis, a cognitive Bayesian modeling framework, to echolocation, an active sensing modality in which the agent actively moves and emits sound to acquire new infor- mation from echoes (Fig 1). Through theoretical formulation and computational simulations, we examined how the target search behavior of an echolocating infotaxis agent varies with important sonar system characteristics, including beam aim, beam coverage, and sensory uncertainty in the form of probabilities of miss and false alarm in target detection. Despite the simplicity of the setup, the infotaxis algorithm produced a rich repertoire of adaptive behaviors across parameter combinations, revealing how decision rules interact with sonar system properties to shape search strategies and performance.

Our results show that the characteristic infotaxis balance between exploration and exploitation observed in olfactory sensing also emerges under echolocation. In olfactory search, the agent chooses between moving to a neighboring location or remaining stationary to maximize the expected reduction in entropy of the search scene (Vergassola *et al*., 2007). In echolocation search, the agent chooses where to aim its sonar beam, either revisiting cells already covered by previous pings or exploring a previously unobserved area, and can select among different sonar beam footprint sizes (Sec. II B). This trade-off is evident when considering agent behavior under sensory uncertainty: Directing the beam to cover previously unobserved cells allows the agent to rapidly identify a probable target region, but slows detection if the target was missed in earlier echo returns; repeating the same beam aim allows the agent to confirm the validity of a target detection, but does not improve localization (Figs 3A, 3D, 3E and 8). Similarly, using a large beam footprint enables quick survey of the search space, but provides limited information on the precise target location (Fig 3B- 3C). Importantly, accumulation of evidence remains a fundamental property of the echolocation infotaxis model, in which the binary hypothesis test observation model gives rise to a spatially graded information landscape that supports the exploration-exploitation trade-off. Indeed, the posterior distribution of target location continues to update even under consecutive pings with the same beam aim, a scenario analogous to the agent remaining stationary in olfactory sensing.

Our analytical and simulation results also show that the probabilities of false alarm and miss influence the infotaxis agent behavior in distinct ways. In the single target search context, *P*_*FA*_ has a much stronger impact on action selection than *P*_*M*_. When *P*_*FA*_ is high, the uncertainty about whether a detection is true drives the infotaxis agent to repeat the same beam aim after receiving a detection (Fig 8) and to favor beam positions that cover fewer cells (Fig B in S2 Appendix), in order to minimize the likelihood of false alarms. This is because even though *P*_*FA*_ is defined for each individual cell, the agent experiences a combined effective false alarm rate from all cells covered within the beam footprint (see Eq 11-12). The cost of misses in overall search length is far greater than the cost of false alarms: The infotaxis agent can recover from false alarms efficiently within a few pings (Fig 3D and 3I) but cannot recognize a miss until it has explored all other cells in the search space (Fig 3E and 3J). These differences mirror real-world sonar operational consequences, where false alarms prompt immediate actions and missed detections often incur substantially higher costs (Urick, 1983).

Our simulations further showed that when the agent underestimates the sensory uncertainty, search success rate drops rapidly (Fig 5), with the agent often terminating the search prematurely at an incorrect target location (Fig J in S2 Appendix). On the other hand, when the agent overes- timates the sensory uncertainty, while the success rate increases, the number of pings required to complete a search also increases (Fig 6). Together, these results deliver an important message that the efficiency and reliability of infotaxis ultimately depends on the quality of sensory information. As the quality degrades, performance declines accordingly.

The information-based nature of infotaxis becomes particularly apparent when compared with MAP searches. Although both agents account for sensory uncertainty when updating their internal world model (Eq 1), only the infotaxis agent considers this uncertainty when selecting the next set of actions. *P*_*M*_ and *P*_*FA*_ are explicitly incorporated in the infotaxis decision making process (Eq 6), whereas the MAP agent simply treats the most probable target region as true (Eq 16). This difference is also reflected in computational time: The infotaxis agent takes longer to make deliberative decisions than the MAP agent’s reflexive ones (Fig S in S2 Appendix). However, in both biological and engineered systems, neural or computational decision making typically occurs on a millisecond timescale, which is orders of magnitude faster than the execution of motor actions such as beam steering, head turning, or locomotion (Ridgway *et al*., 2014). Thus, the longer deliberation time under infotaxis would have negligible impact on overall operational efficiency. Instead, the infotaxis agent is behaviorally more efficient and reliable, consistently completing searches with fewer pings at the correct target location (Fig 4).

The Bayesian probabilistic framework underlying infotaxis provides a principled avenue to di- rectly integrate sensory and motor capabilities derived from behavioral or neurophysiological ex- periments into a cognitive model that makes information-based decisions (Fig 1B). This integration operates through two key components: the likelihood function and the motor action. The likelihood function defines how the agent interprets echoes given the task goal and the quality of sensory evi- dence, providing a natural entry point for injecting experimental knowledge into the model. For ex- ample, although echoes can be represented by multidimensional temporal, spectral, and statistical features, how an echolocating agent perceives and uses these features is highly context-dependent. An agent may attend to fine spectral cues tied to position within the beam when localizing a target (Arditi *et al*., 2015; Kloepper *et al*., 2018; Müller *et al*., 2017), but emphasize broad spectral pat- terns when classifying targets (Accomando *et al*., 2020, 2022). Echo interpretation is also shaped by environmental noise and prior experience of the experimental subject. The Bayesian framework in infotaxis allows these factors to be incorporated into task-, environment-, and even individual- dependent likelihood functions for more realistic modeling of echolocation behavior. However, we note that successive echo observations are assumed independent in the likelihood functions used here (Sec. II B). In realistic echolocation scenarios, consecutive echoes may be temporally correlated because the acoustic scene often changes gradually across pings, and because auditory processing may integrate information over time. Such correlations would reduce the effective information gain from each ping and slow the posterior updates. Therefore, a temporally correlated observation model that incorporates recent echo history, either empirically or through physics-based models, will be important for using the infotaxis model to interpret experimental data.

Motor actions define the space of behavioral choices available to an agent to achieve its task goal. These actions correspond to measurable observables in behavioral experiments, and thus form a crucial bridge between model and data. In echolocation, motor actions include adjustments to acoustic parameters, such as the spectral and temporal characteristics of the emitted echolocation signals, and to movement parameters, including conformational changes of the sound production and reception apparatus [e.g., shape adjustments of the mouth, noseleaf, ears (bats), melon, and the associated air sacs (toothed whales)], rotational movements (e.g., head turning, body reorientation), and locomotion (Moss *et al*., 2023). These adjustments are often coupled and can be constrained by physical and physiological limitations to ensure modeled behavioral outputs remain biologically realistic.

Infotaxis provides a versatile framework for interpreting experimentally observed echolocation behaviors, inferring the underlying cognitive processes, and generating testable hypotheses for further model refinement. By systematically varying model parameters, simulated motor action sequences can be quantitatively compared with experimental data either via summary statistics or on a ping-by-ping basis (Calhoun *et al*., 2014; Najemnik and Geisler, 2009; Pang *et al*., 2018; Yang *et al*., 2016). Changes in model parameterization can also enable numerical experiments to inform the design of behavioral experiments. For example, our simulations showed that the infotaxis agent selects an increasingly small beam footprint to precisely locate the target in the search space (Fig 3B-3C and 3G-3H). This progressive narrowing of the beam mirrors how the projected beam footprint would change for an echolocator approaching a target screen while maintaining a constant beamwidth, and is reminiscent of the narrower beamwidths bats use to precisely locate a landing platform (Eitan *et al*., 2022). However, bats and toothed whales typically broaden their sonar beams just before intercepting mobile prey, likely to keep the prey within the sensing volume in case of evasive maneuvers (e.g., Matsuta *et al*., 2013; Wisniewska *et al*., 2015). While it is unclear how echolocating animals hunting for stationary prey adjust their beam aim and beamwidth at fine scale (Geipel *et al*., 2019), would incorporating potential target motion into the model alter the infotaxis choices? Beyond sensory control, the cost of movement has also been proposed as a factor that balances information gain in shaping animal trajectories during tracking and localization tasks (Chen *et al*., 2020; Liu *et al*., 2024; Loisy and Eloy, 2022). Paired numerical and behavioral experiments could test whether similar trade-offs occur in echolocation, a distal sensing modality in which adjustments of beam aim, beam coverage, and transmission rate likely incur lower energetic costs than in proximal sensing modalities such as tactile sensing, and electrolocation, which has a substantially smaller sensing volume than echolocation (Nelson and MacIver, 2006). In our simulations, restricting beam movements without explicitly including movement cost in the decision rule had minimal effect on search efficiency (Fig A in S2 Appendix). Even though we were not able to identify existing behavioral datasets suitable for direct com- parison with the single target search model presented here, we have conducted controlled robotic experiments showing successful infotaxis-based sonar target localization in a one-dimensional search space under varying beamwidth conditions (Mboya *et al*., 2026) and mismatched sensory uncer- tainty (Mudassir *et al*., under review). These studies provide initial validation of echolocation infotaxis on a robotic platform and motivate future tests using animal behavioral experiments. Such model-data comparisons will require mapping the abstract model parameters presented here to specific experimental contexts. For example, beam footprint size is species-specific and often varies dynamically depending on task priorities (e.g., Jensen *et al*., 2015; Motoi *et al*., 2017), whereas sensory uncertainty is further tied to intrinsic target scattering properties, which influence the animal’s ability to detect or discriminate targets (e.g., Wei *et al*., 2021).

The echolocation infotaxis model presented in this paper differs in setup and objectives from deep RL models of echolocating bats (Mohan and Vanderelst, 2020; Nguyen and Vanderelst, 2022, 2025), which capture similar cognitive and sensorimotor aspects of decision making in echolocation. To investigate intrinsic properties of infotaxis search in echolocation, our model is intentionally ab- stract: the complexity and uncertainty of auditory scene analysis and target echo detection are encapsulated by *P*_*M*_ and *P*_*FA*_, whose influence on action selection can be explicitly traced. This simplified observation model was chosen deliberately to create a tractable framework in which the resulting decision behavior could be analyzed analytically and computationally. In contrast, the deep RL models operate directly on simulated echo time series and learn action policies through trial-and-error to maximize cumulative reward. The infotaxis agent maintains an explicit proba- bilistic world model tied to the task goal (e.g., a distribution over target location in search tasks or target category in discrimination tasks), whereas in RL-based approaches, task-relevant informa- tion is encoded within learned policy or value representations. Both infotaxis and RL approaches generally require reformulation when the task context changes. RL approaches typically require retraining after such reformulation, whereas infotaxis action selection is determined directly from the probabilistic model and objective function. Selecting actions in infotaxis can become com- putationally expensive when the motor action space is large, as each decision involves evaluating the expected entropy over *all* possible action combinations. In comparison, although RL train- ing can be time- and resource-intensive, once trained, action selection is computationally efficient. Taken together, these differences highlight different trade-offs: infotaxis provides a directly analyz- able probabilistic formulation of action selection, while RL enables rapid action selection through learned policies. We postulate that training a surrogate neural network model to approximate infotaxis action selection could substantially reduce its run-time computational cost. The demon- strated success of neural networks in capturing structured internal control during bat navigation (Teshima *et al*., 2025) supports the feasibility of this hybrid approach.

In summary, we developed an infotaxis model for echolocation that integrates sensory uncer- tainty, motor action, and information-based reasoning in a single interpretable framework. Im- plemented in a simple target search scenario, the cognitive infotaxis agent proved more reliable and efficient than a reflexive MAP agent, though its performance declined as sensory informa- tion degraded. This work lays the foundation for extending infotaxis modeling to more complex echolocation and active sonar contexts, such as multi-target search among clutter, and provide interpretable linkages between behavioral observables, sensory mechanisms, and decision making processes. We propose that infotaxis offers a powerful framework for iterative model refinement and experimental testing that can help understand sonar-guided autonomy in natural and engineered systems.

## Supporting information

S1 Appendix (all supporting texts)

S2 Appendix (all supporting figures)

S3 Appendix (all supporting tables)

## ACKNOWLEDGMENTS

We thank Adrienne Fairhall of the University of Washington for introducing W.-J. L. to the seminal infotaxis paper.

## AUTHOR CONTRIBUTIONS

W.-J.L.: Conceptualization, Formal Analysis, Funding Acquisition, Investigation, Methodology, Project administration, Resources, Software, Visualization, Writing – Original Draft Preparation, Writing – Review & Editing. J.R.B.: Funding Acquisition, Investigation, Methodology, Project administration, Visualization, Writing – Review & Editing. P.L.T.: Funding Acquisition, Project administration, Writing – Review & Editing.

## DATA AVAILABILITY

All code and scripts for the model, simulation, and figure generation are available in the GitHub repository: https://github.com/uw-echospace/infotaxis-search-single-target.

## APPENDIX: LIST OF SYMBOLS

*A*: Search space: set of possible target locations
*ℬ*_**s**_: Set of cells covered by the beam footprint
*E*_*X*_ [*H*_**s**_(*K*)]: Expected entropy after action **s** over possible observations *X*
*H*(*K*): Entropy of the posterior of target location
*H*_**s**_(*K*|*X*): Entropy of the posterior of target location after action *s* and observation *X*
*K*: Random variable representing target location
*K*: Specific candidate target location (cell)
*ℒ*: Likelihood
*N*_*A*_: Number of cells in the search space
*N*_*B*_: Number of cells covered by the beam footprint
*N*: Ping index
*P*_*FA*_: Probability of false alarm
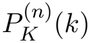: Posterior probability of target location after ping *n*
*P*_*M*_: Probability of miss
*P*_*max*_: Maximum posterior probability of target location 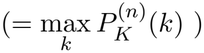
*p*_*th*_: Search termination threshold for *P*_*max*_ *r*_*A*_ Radius of the search space
*r*_*B*_: Radius of the beam footprint
*S*: Set of possible sensing actions
S: Agent action; consisting of beam aim and beam footprint size in single target search
*X*: Echo observation (*X* = 1 detection, *X* = 0 no detection)
*α*: Relative proportion of the beam footprint within the search space (= *N*_*B*_*/N*_*A*_)
*θ*_*B*_: Beam aim

## List of Supporting Information Files

**S1 Appendix**. Supporting derivations and analyses, including the effects of beampattern- modulated sensory uncertainty.

**S2 Appendix**. Supporting figures.

**S3 Appendix**. Supporting tables containing model parameters.

**S1 Video**. An infotaxis search with a fixed beam footprint size *r*_*B*_ = 1. Selected pings are shown in Fig 3A.

**S2 Video**. An infotaxis search with two beam footprint sizes *r*_*B*_ = {1, 2}. Selected pings are shown in Fig 3B.

**S3 Video**. An infotaxis search with three beam footprint sizes *r*_*B*_ = {1, 2, 3}. Selected pings are shown in Fig 3C.

**S4 Video**. An infotaxis search with a false alarm. Selected pings are shown in Fig 3D.

**S5 Video**. An infotaxis search with a miss. Selected pings are shown in Fig 3E.

**S6 Video**. A MAP search. Selected pings are shown in Fig 9A.

**S7 Video**. An infotaxis search with beam aim movement restriction *r*_*M*_ = 3. Selected pings are shown in panel A of Fig A in S2 Appendix.

**S8 Video**. An infotaxis search with one miss occurring early in the search. Selected pings are shown in panel A of Fig C in S2 Appendix.

**S9 Video**. An infotaxis search with one miss occurring later in the search. Selected pings are shown in panel B of Fig C in S2 Appendix.

**S10 Video**. An infotaxis search when the agent underestimates the sensory uncertainty. Se- lected pings are shown in panel A of Fig J in S2 Appendix.

