## Supplementary material for "Modeling echolocation as an active pursuit of information via infotaxis": S1 Appendix (all supporting texts)

### S1 APPENDIX: SUPPORTING TEXT

#### Section A : Single target search with beampattern-modulated $P_M$ and $P_{FA}$

To conduct a preliminary investigation on the effects of incorporating directional characteristics of echolocation, we devised a simple beampattern model in which $P_M$ and $P_{FA}$ vary spatially depending on a cell's distance from the beam aim. In echolocating animals, sensing reliability generally vary across direction, with measurements near the beam edge expected to exhibit greater uncertainty than those near the beam center. Therefore, we modulate these probabilities by a multiplicative factor $B$, modeled as an inverted Gaussian-shaped curve:

$$B = \lambda_B \left[1 - \exp\left(-\frac{r_c^2}{2{\sigma_B}^2}\right)\right] + 1, \tag{S1A-1}$$

where $\lambda_B$ controls the magnitude of modulation, $\sigma_B$ controls the width of modulation, and $r_c$ is the distance of the cell from the beam aim. This modulation elevates $P_M$ and $P_{FA}$ toward the edge of the beam.

Simulations show that infotaxis searches with beampattern-modulated sensory uncertainty generally become slower and have moderately lower success rates, with the effects being more pronounced at higher baseline $P_M$ and $P_{FA}$ (Fig T in S2 Appendix). These results are intuitive: higher sensory uncertainty at the edge of the beam effectively reduces the beam footprint size, which can lengthen the initial exploration phase until the first target detection. On the other hand, the beampattern modulation also causes the posterior $P_K^{(n)}(k)$ to peak at the center of the beam after a target detection. Once a detection is received, the spatially varying posterior and the infotaxis rule work in concert to quickly narrow down the target location.

Interestingly, the beampattern-modulated sensory uncertainty affects MAP searches much more severely, resulting in both lower success rates and substantially more pings required to complete a search (Fig U in S2 Appendix). As in the infotaxis case, higher sensory uncertainty near the beam edge effectively reduces the beam footprint size. However, detailed ping-by-ping inspection of the search sequences (not shown) indicates that the spatially varying posterior $P_K^{(n)}(k)$ can cause the MAP beam aim to become "stuck" at a cell neighboring the true target location. This leads to slower posterior updates and correspondingly slower changes in $P_{max}$, often preventing the search from satisfying either of the stopping criteria. Paradoxically, when baseline sensory uncertainty is higher, the occasional false alarms or misses can prompt the agent to shift beam aim to a different cell, resolving the "stuck" situation, leading to a higher overall success rate.

We note that these results are preliminary based on a simplistic beampattern model. More detailed analyses across different parameter value ranges and spatial structures of sensory uncertainty within the beam would be needed to fully understand the model search behavior and to support comparison with experimental data.

**Section B : Detailed derivation for entropy decrease in the exploration phase**

From Sec. IIIB,

$$P_K^{(1)}(k) \approx \begin{cases} \dfrac{1}{N_A(1-\alpha)}\dfrac{P_M}{1-P_{FA}}\left[1-\dfrac{\alpha}{1-\alpha}\dfrac{P_M}{1-P_{FA}}\right], & k \in \mathcal{B}_{s_1} \\ \dfrac{1}{N_A(1-\alpha)}\left[1-\dfrac{\alpha}{1-\alpha}\dfrac{P_M}{1-P_{FA}}\right], & \text{otherwise.} \end{cases}$$

$$P_K^{(2)}(k) \approx \begin{cases} \dfrac{1}{N_A(1-2\alpha)}\dfrac{P_M}{1-P_{FA}}\left[1-\dfrac{2\alpha}{1-2\alpha}\dfrac{P_M}{1-P_{FA}}\right], & k \in \mathcal{B}_{s_1}\cup\mathcal{B}_{s_2} \\ \dfrac{1}{N_A(1-2\alpha)}\left[1-\dfrac{2\alpha}{1-2\alpha}\dfrac{P_M}{1-P_{FA}}\right], & \text{otherwise.} \end{cases} \quad \text{(S1B-1)}$$

$$\vdots$$

$$P_K^{(n)}(k) \approx \begin{cases} \dfrac{1}{N_A(1-n\alpha)}\dfrac{P_M}{1-P_{FA}}\left[1-\dfrac{n\alpha}{1-n\alpha}\dfrac{P_M}{1-P_{FA}}\right], & k \in \bigcup_{m=1}^{n}\mathcal{B}_{s_m} \\ \dfrac{1}{N_A(1-n\alpha)}\left[1-\dfrac{n\alpha}{1-n\alpha}\dfrac{P_M}{1-P_{FA}}\right], & \text{otherwise} \end{cases}$$

The entropy of the updated scene after transmission $n$ is

$$H^{(n)} = -\sum_{k\in\bigcup_{m=1}^{n}\mathcal{B}_{s_m}} P_K^{(n)}(k)\log_2 P_K^{(n)}(k) - \sum_{k\notin\bigcup_{m=1}^{n}\mathcal{B}_{s_m}} P_K^{(n)}(k)\log_2 P_K^{(n)}(k) \quad \text{(S1B-2)}$$

Since consecutive beam footprints would not overlap under the infotaxis selection, the first term sums over $n\alpha N_A$ cells, and the second term sums over $(1-n\alpha)N_A$ cells:

$$\begin{aligned} \sum_{k\in\bigcup_{m=1}^{n}\mathcal{B}_{s_m}} P_K^{(n)}(k)\log_2 P_K^{(n)}(k) &= \frac{n\alpha N_A}{N_A(1-n\alpha)}\frac{P_M}{1-P_{FA}}\left[1-\frac{n\alpha}{1-n\alpha}\frac{P_M}{1-P_{FA}}\right] \\ &\times \log_2\left[\frac{1}{N_A(1-n\alpha)}\frac{P_M}{1-P_{FA}}\left[1-\frac{n\alpha}{1-n\alpha}\frac{P_M}{1-P_{FA}}\right]\right] \end{aligned} \quad \text{(S1B-3)}$$

$$\begin{aligned} \sum_{k\notin\bigcup_{m=1}^{n}\mathcal{B}_{s_m}} P_K^{(n)}(k)\log_2 P_K^{(n)}(k) &= \frac{N_A(1-n\alpha)}{N_A(1-n\alpha)}\left[1-\frac{n\alpha}{1-n\alpha}\frac{P_M}{1-P_{FA}}\right] \\ &\times \log_2\left[\frac{1}{N_A(1-n\alpha)}\left[1-\frac{n\alpha}{1-n\alpha}\frac{P_M}{1-P_{FA}}\right]\right] \end{aligned} \quad \text{(S1B-4)}$$

Let $\beta = \frac{n\alpha}{1-n\alpha}\frac{P_M}{1-P_{FA}}$,

$$\begin{aligned} H^{(n)} &= \beta(1-\beta)\left[\log_2 N_A(1-n\alpha) - \log_2\frac{P_M}{1-P_{FA}} - \log_2(1-\beta)\right] \\ &+ (1-\beta)\left[\log_2 N_A(1-n\alpha) - \log_2(1-\beta)\right] \\ &= (1-\beta^2)\left[\log_2 N_A(1-n\alpha) - \log_2(1-\beta)\right] - \beta(1-\beta)\log_2\frac{P_M}{1-P_{FA}} \end{aligned} \quad \text{(S1B-5)}$$

When $P_M$ and $\alpha$ are small and thus $\beta$ and $\beta^2$ are both small,

$$H^{(n)} \approx \log_2 N_A(1-n\alpha) = \log_2(N_A - nN_B) = \log_2\left|\mathcal{A}\setminus\bigcup_{m=1}^{n}\mathcal{B}_{s_m}\right|, \quad \text{(S1B-6)}$$

which is the entropy for a uniform distribution over the set of cells in the search space $\mathcal{A}$ not yet observed.

#### Section C : Detailed derivation for quantities related to beam aim selection after the first target detection

##### C.1 Probability distribution of target location after receiving the first detection

In the derivation below, superscripts $\widetilde{(0)}$, $\widetilde{(1)}$, and $\widetilde{(2)}$ indicate quantities associated with the ping before the first target detection, the first ping that yields a target detection, and the subsequent ping, respectively. Following Eq 17, the distribution of target probability after receiving a target detection is

$$P_K^{\widetilde{(1)}}(k) = \frac{P_K^{\widetilde{(0)}}(k)\mathcal{L}(k|X=1;S=s_1)}{\mathcal{C}_{X=1}^{(1)}} \tag{S1C-1}$$

where

$$\mathcal{L}(k|X=1;S=s_1)\begin{cases} 1-P_M(1-P_{FA})^{N_B-1} & \text{, for } k\in\mathcal{B}_{s_1} \\ 1-(1-P_{FA})^{N_B} & \text{, otherwise}\end{cases} \tag{S1C-2}$$

and

$$\begin{aligned} C_{X=1}^{\widetilde{(1)}} &= \sum_{k\in\mathcal{B}_{s_1}} P_K^{\widetilde{(0)}}(k)\mathcal{L}(k|X=1) + \sum_{k\notin\mathcal{B}_{s_1}} P_K^{\widetilde{(0)}}(k)\mathcal{L}(k|X=1) \\ &= P_K^{\widetilde{(0)}}(k)\Big[\alpha N_A\big[1-P_M(1-P_{FA})^{N_B-1}\big]+(1-\alpha)N_A\big[1-(1-P_{FA})^{N_B}\big]\Big]\end{aligned} \tag{S1C-3}$$

Let $N_B=\alpha N_A$ and use the binomial approximation $(1+y)^a\approx 1+ay$,

$$\begin{aligned} 1-P_M(1-P_{FA})^{N_B-1} &\approx 1-P_M\Big[1-(N_B-1)P_{FA}\Big] = 1-P_M+P_{FA}(N_B-1)\approx 1-P_M \\ 1-(1-P_{FA})^{N_B} &\approx 1-(1-N_BP_{FA}) = \alpha N_A P_{FA}\end{aligned} \tag{S1C-4}$$

We then obtain

$$\mathcal{C}_{X=1}^{\widetilde{(1)}} \approx P_K^{\widetilde{(0)}}(k)N_A\big[\alpha(1-P_M)+(1-\alpha)\alpha N_A P_{FA}\big] = P_K^{\widetilde{(0)}}(k)N_A\gamma_1, \tag{S1C-5}$$

where

$$\gamma_1 = \alpha(1-P_M)+(1-\alpha)\alpha N_A P_{FA}. \tag{S1C-6}$$

Then,

$$\begin{aligned} P_K^{\widetilde{(1)}}(k) &= \begin{cases} \dfrac{P_K^{(0)}(k)(1-P_M)}{\mathcal{C}_{X=1}^{(1)}} & \text{, for } k\in\mathcal{B}_{s_1} \\ \dfrac{P_K^{(0)}(k)(\alpha N_A P_{FA})}{\mathcal{C}_{X=1}^{(1)}} & \text{, otherwise}\end{cases} \\ &= \begin{cases} \dfrac{1}{N_A}\dfrac{1-P_M}{\gamma_1} & \text{, for } k\in\mathcal{B}_{s_1} \\ \dfrac{1}{N_A}\dfrac{\alpha N_A P_{FA}}{\gamma_1} & \text{, otherwise}\end{cases}\end{aligned} \tag{S1C-7}$$

##### C.2 If the subsequent ping repeats the same beam aim

Now consider the scenario when the agent points the next beam aim at the same cell as in the first transmission.

If $X = 1$, using the approximations in Eq S1C-4 and recall $N_B = \alpha N_A$,

$$
\begin{aligned}
\widetilde{P_K^{(2)}}(k) &= \begin{cases} \dfrac{\widetilde{P_K^{(1)}}(k)\Big[1 - P_M(1-P_{FA})^{N_B-1}\Big]}{\widetilde{C_{X=1}^{(2)}}} & \text{, for } k \in \mathcal{B}_{s_1} = \mathcal{B}_{s_2} \\ \dfrac{\widetilde{P_K^{(1)}}(k)\Big[1-(1-P_{FA})^{N_B}\Big]}{\widetilde{C_{X=1}^{(2)}}} & \text{, otherwise} \end{cases} \\
&\approx \begin{cases} \dfrac{\widetilde{P_K^{(1)}}(k)(1-P_M)}{\widetilde{C_{X=1}^{(2)}}} & \text{, for } k \in \mathcal{B}_{s_1} = \mathcal{B}_{s_2} \\ \dfrac{\widetilde{P_K^{(1)}}(k)(\alpha N_A P_{FA})}{\widetilde{C_{X=1}^{(2)}}} & \text{, otherwise} \end{cases}
\end{aligned} \tag{S1C-8}
$$

where

$$
\begin{aligned}
\widetilde{C_{X=1}^{(2)}} &= \sum_{k \in \mathcal{B}_{s_1} = \mathcal{B}_{s_2}} \widetilde{P_K^{(1)}}(k) + \sum_{k \notin \mathcal{B}_{s_1} = \mathcal{B}_{s_2}} \widetilde{P_K^{(1)}}(k) \\
&= \alpha N_A \Big( \frac{1}{N_A} \frac{1-P_M}{\gamma_1} (1-P_M) \Big) + (1-\alpha) N_A \Big( \frac{1}{N_A} \frac{\alpha N_A P_{FA}}{\gamma_1} (\alpha N_A P_{FA}) \Big) \\
&= \frac{1}{\gamma_1} \Big[ \alpha (1-P_M)^2 + (1-\alpha)(\alpha N_A P_{FA})^2 \Big]
\end{aligned} \tag{S1C-9}
$$

If $X = 0$, similarly using the the approximations in Eq S1C-4 and additionally $1 - \alpha N_A P_{FA} + P_{FA} \approx 1$

$$
\begin{aligned}
\widetilde{P_K^{(2)}}(k) &= \begin{cases} \dfrac{\widetilde{P_K^{(1)}}(k) P_M (1-P_{FA})^{N_B-1}}{\widetilde{C_{X=0}^{(2)}}} & \text{, for } k \in \mathcal{B}_{s_1} = \mathcal{B}_{s_2} \\ \dfrac{\widetilde{P_K^{(1)}}(k)(1-P_{FA})^{N_B}}{\widetilde{C_{X=0}^{(2)}}} & \text{, otherwise} \end{cases} \\
&\approx \begin{cases} \dfrac{\widetilde{P_K^{(1)}}(k) P_M}{\widetilde{C_{X=0}^{(2)}}} & \text{, for } k \in \mathcal{B}_{s_1} = \mathcal{B}_{s_2} \\ \dfrac{\widetilde{P_K^{(1)}}(k)(1-\alpha N_A P_{FA})}{\widetilde{C_{X=0}^{(2)}}} & \text{, otherwise} \end{cases}
\end{aligned} \tag{S1C-10}
$$

where

$$
\begin{aligned}
C^{\widetilde{(2)}}_{X=0} &= \alpha N_A\Big(\frac{1}{N_A}\frac{1-P_M}{\gamma_1}\Big)P_M + (1-\alpha)N_A\Big(\frac{1}{N_A}\frac{\alpha N_A P_{FA}}{\gamma_1}\Big)(1-\alpha N_A P_{FA}) \\
&= \frac{1}{\gamma_1}\Big[\alpha P_M(1-P_M) + (1-\alpha)\alpha N_A P_{FA}(1-\alpha N_A P_{FA})\Big] \\
&= \frac{1}{\gamma_1}\Big[\alpha P_M(1-P_M) + (1-\alpha)\alpha N_A P_{FA} - (1-\alpha)(\alpha N_A P_{FA})^2\Big]
\end{aligned}
\tag{S1C-11}
$$

Let $\gamma_2 = \alpha(1-P_M)^2 + (1-\alpha)(\alpha N_A P_{FA})^2$, we then obtain

$$
\begin{aligned}
C^{\widetilde{(2)}}_{X=1} &= \frac{\gamma_2}{\gamma_1} \\
C^{\widetilde{(2)}}_{X=0} &= \frac{\gamma_1-\gamma_2}{\gamma_1}
\end{aligned}
\tag{S1C-12}
$$

The probability of receiving an echo observation $X = x$ from the subsequent ping is

$$
P(X = x|s_2) = \sum_{k\in\mathcal{B}_{s_1}} P(X = x|k; s_2) + \sum_{k\notin\mathcal{B}_{s_1}} P(X = x|k; s_2) = C^{\widetilde{(2)}}_{X=x}, \tag{S1C-13}
$$

where $s_1 = s_2$ and $\mathcal{B}_{s_1} = \mathcal{B}_{s_2}$.

##### C.3 If the subsequent beam aim shifts to a neighboring cell

Now consider the scenario when the beam aim of the second ping is aimed at a neighboring cell. Following the same procedure as the above, we partition the search space into four different sections

- $\mathcal{A}_1$: cells covered by both ping 1 and ping 2 beam footprints
- $\mathcal{A}_2$: cells covered by only ping 1 beam footprint but not ping 2
- $\mathcal{A}_3$: cells covered by only ping 2 beam footprint but not ping 1
- $\mathcal{A}_4$: cells not covered by either ping 1 or ping 2 beam footprint

and compute the update target probability map $P^{\widetilde{(2)}}_K(k)$ and the probability of receiving an echo or no echo after the second ping, $P(X = 1)$ and $P(X = 0)$.

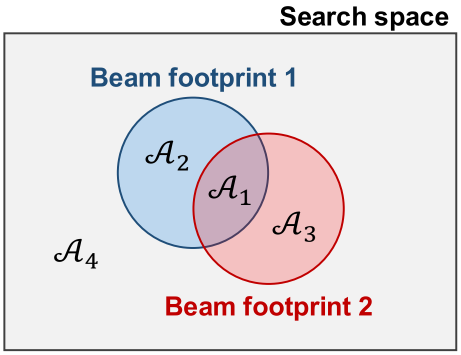

If $X=1$, let $\alpha_{12}$ be the proportion of cells covered by the beam footprints of both ping 1 and ping 2, we can obtain

$$\widetilde{P_K^{(2)}}(k)=\begin{cases}\dfrac{\frac{1}{N_A}(1-P_M)^2}{\widetilde{C_{X=1}^{(2)}}} & ,k\in\mathcal{A}_1\\ \dfrac{\frac{1}{N_A}(\alpha N_A P_{FA})(1-P_M)}{\widetilde{C_{X=1}^{(2)}}} & ,k\in\mathcal{A}_2 \text{ or } k\in\mathcal{A}_3\\ \dfrac{\frac{1}{N_A}(\alpha N_A P_{FA})^2}{\widetilde{C_{X=1}^{(2)}}} & ,k\in\mathcal{A}_4\end{cases} \tag{S1C-14}$$

where

$$\begin{aligned}\widetilde{C_{X=1}^{(2)}}&=\sum_{k\in\mathcal{A}_1}\widetilde{P_K^{(1)}}(k)+\sum_{k\in\mathcal{A}_2}\widetilde{P_K^{(1)}}(k)+\sum_{k\in\mathcal{A}_3}\widetilde{P_K^{(1)}}(k)+\sum_{k\in\mathcal{A}_4}\widetilde{P_K^{(1)}}(k)\\ &=\alpha_{12}N_A\Big(\frac{1}{N_A}\frac{1-P_M}{\gamma_1}\Big)(1-P_M)\\ &\quad+(\alpha-\alpha_{12})N_A\Big(\frac{1}{N_A}\frac{1-P_M}{\gamma_1}\Big)(\alpha N_A P_{FA})\\ &\quad+(\alpha-\alpha_{12})N_A\Big(\frac{1}{N_A}\frac{\alpha N_A P_{FA}}{\gamma_1}\Big)(1-P_M)\\ &\quad+(1-2\alpha+\alpha_{12})N_A\Big(\frac{1}{N_A}\frac{\alpha N_A P_{FA}}{\gamma_1}\Big)(\alpha N_A P_{FA})\\ &=\frac{1}{\gamma_1}\Big[\alpha_{12}(1-P_M)^2+2(\alpha-\alpha_{12})\alpha N_A P_{FA}(1-P_M)+(1-2\alpha+\alpha_{12})(\alpha N_A P_{FA})^2\Big]\end{aligned} \tag{S1C-15}$$

Let

$$\gamma_2'=\alpha_{12}(1-P_M)^2+2(\alpha-\alpha_{12})\alpha N_A P_{FA}(1-P_M)+(1-2\alpha+\alpha_{12})(\alpha N_A P_{FA})^2 \tag{S1C-16}$$

then

$$\widetilde{C_{X=1}^{(2)}}=\frac{\gamma_2'}{\gamma_1} \tag{S1C-17}$$

If $X=0$,

$$\widetilde{P_k^{(2)}}(k)=\begin{cases}\dfrac{\frac{1}{N_A}(1-P_M)P_M}{\widetilde{C_{X=0}^{(2)}}} & ,k\in\mathcal{A}_1\\ \dfrac{\frac{1}{N_A}(1-P_M)(1-\alpha N_A P_{FA})}{\widetilde{C_{X=0}^{(2)}}} & ,k\in\mathcal{A}_2\\ \dfrac{\frac{1}{N_A}(\alpha N_A P_{FA})P_M}{\widetilde{C_{X=0}^{(2)}}} & ,k\in\mathcal{A}_3\\ \dfrac{\frac{1}{N_A}(\alpha N_A P_{FA})(1-\alpha N_A P_{FA})}{\widetilde{C_{X=0}^{(2)}}} & ,k\in\mathcal{A}_4\end{cases} \tag{S1C-18}$$

and

$$
\begin{aligned}
C_{X=1}^{\widetilde{(2)}} &= \alpha_{12} N_A \Big(\frac{1}{N_A}\frac{1-P_M}{\gamma_1}\Big) P_M \\
&+ (\alpha - \alpha_{12}) N_A \Big(\frac{1}{N_A}\frac{1-P_M}{\gamma_1}\Big)(1-\alpha N_A P_{FA}) \\
&+ (\alpha - \alpha_{12}) N_A \Big(\frac{1}{N_A}\frac{\alpha N_A P_{FA}}{\gamma_1}\Big) P_M \\
&+ (1 - 2\alpha + \alpha_{12}) N_A \Big(\frac{1}{N_A}\frac{\alpha N_A P_{FA}}{\gamma_1}\Big)(1-\alpha N_A P_{FA}) \\
&= \frac{1}{\gamma_1}\Big[\alpha_{12}(1-P_M)P_M \\
&\qquad + (\alpha - \alpha_{12})(1-P_M)(1-\alpha N_A P_{FA}) \\
&\qquad + (\alpha - \alpha_{12})\alpha N_A P_{FA} P_M \\
&\qquad + (1 - 2\alpha + \alpha_{12})\alpha N_A P_{FA}(1-\alpha N_A P_{FA})\Big] \\
&= \frac{\gamma_1 - \gamma_2'}{\gamma_1}
\end{aligned}
\tag{S1C-19}
$$

The probability of receiving an echo observation $X = x$ from the subsequent ping is

$$
\begin{aligned}
P(X = x|s_2) &= \sum_{k\in\mathcal{A}_1} P(X = x|k; s_2) + \sum_{k\in\mathcal{A}_2} P(X = x|k; s_2) \\
&\quad + \sum_{k\in\mathcal{A}_3} P(X = x|k; s_2) + \sum_{k\in\mathcal{A}_4} P(X = x|k; s_2) \\
&= C_{X=x}^{\widetilde{(2)}}
\end{aligned}
\tag{S1C-20}
$$

where $s_1 \neq s_2$ and $\mathcal{B}_{s_1} \neq \mathcal{B}_{s_2}$.
