## Supplementary material for "Modeling echolocation as an active pursuit of information via infotaxis": S2 Appendix (all supporting figures)

### S2 APPENDIX: SUPPORTING FIGURES

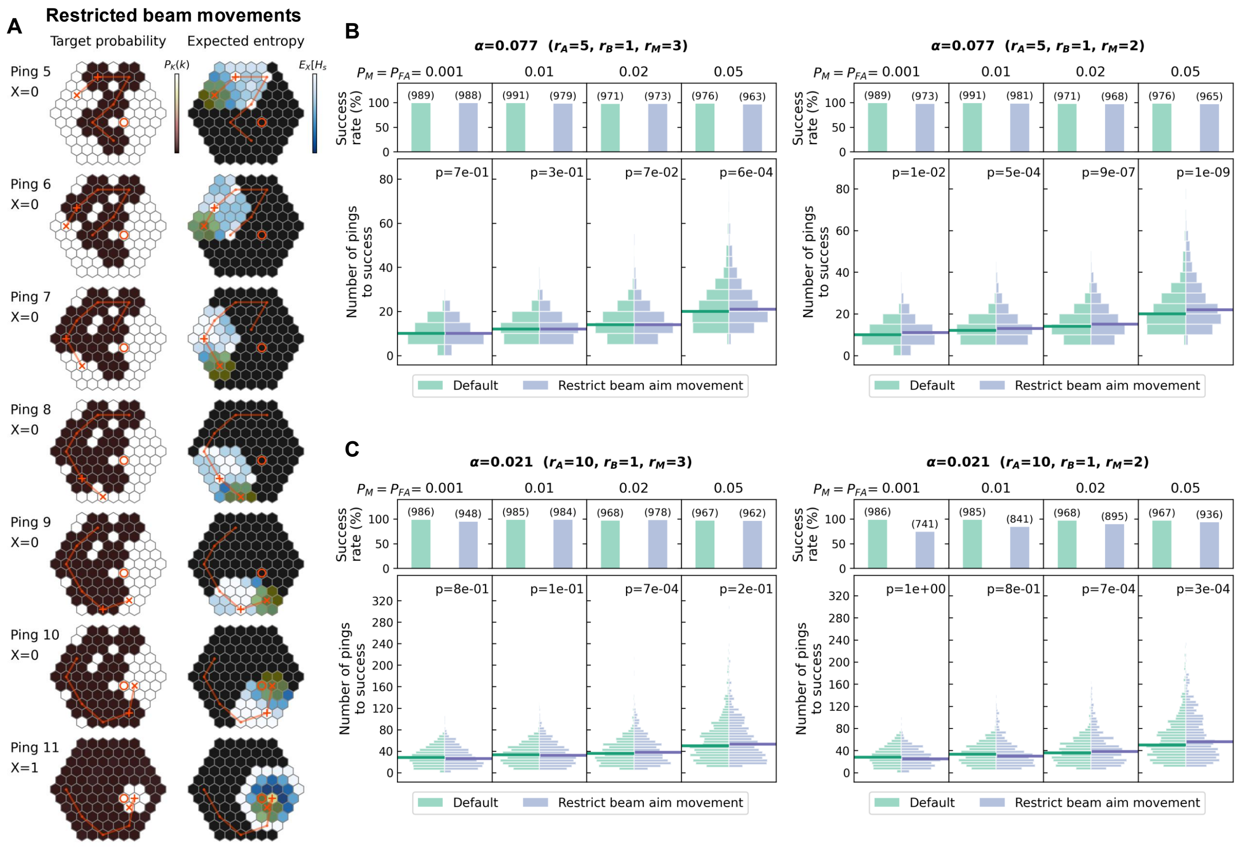


FIG A: **Infotaxis search with beam aim movement restriction.** (A) Probability distribution of target location after selected pings. Restricting beam aim movement produces smoother search trajectories. Black cells in the expected entropy plots (right column) indicate those not considered for beam aim selection. Color represents a relative scale in the value range spanned in the search space. Under infotaxis, only the relative scale matters because the agent selects the next set of actions that gives the lowest expected entropy among all choices. See Fig 3 caption for definitions of all symbols, Table B in S3 Appendix for all search parameter values, and S7 Video for the full sequence. (B-C) Comparison of the number of pings required to complete a search without (green on the left) and with (purple on the right) beam aim movement restriction under two search space radii $r_A$ and two beam footprint size $r_B$. $r_M$ in each panel title indicates the radius of restricted beam aim movement. Searches with smaller $r_M$ take longer to complete. The reported $p$-values are from Mann-Whitney U tests under the alternative hypothesis that searches without beam aim movement restriction requires fewer pings to complete a search. See Sec. IID for definitions of all quantities and details of the stopping condition.

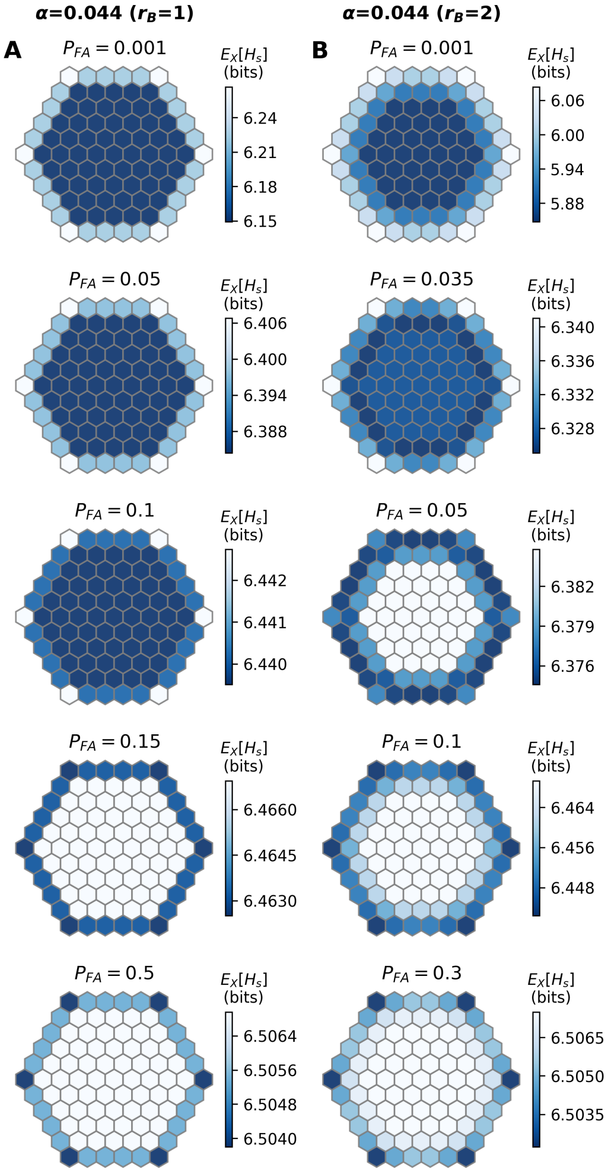


FIG B: **Expected entropy at the beginning of a search.** Expected entropy computed for two beam footprint radii $r_B = 1$ (A) and $r_B = 2$ (B) at selected $P_{FA}$ values. The $P_{FA}$ value at which the minimum expected entropy shifts to cells near the search space boundary is smaller for larger beam footprints.

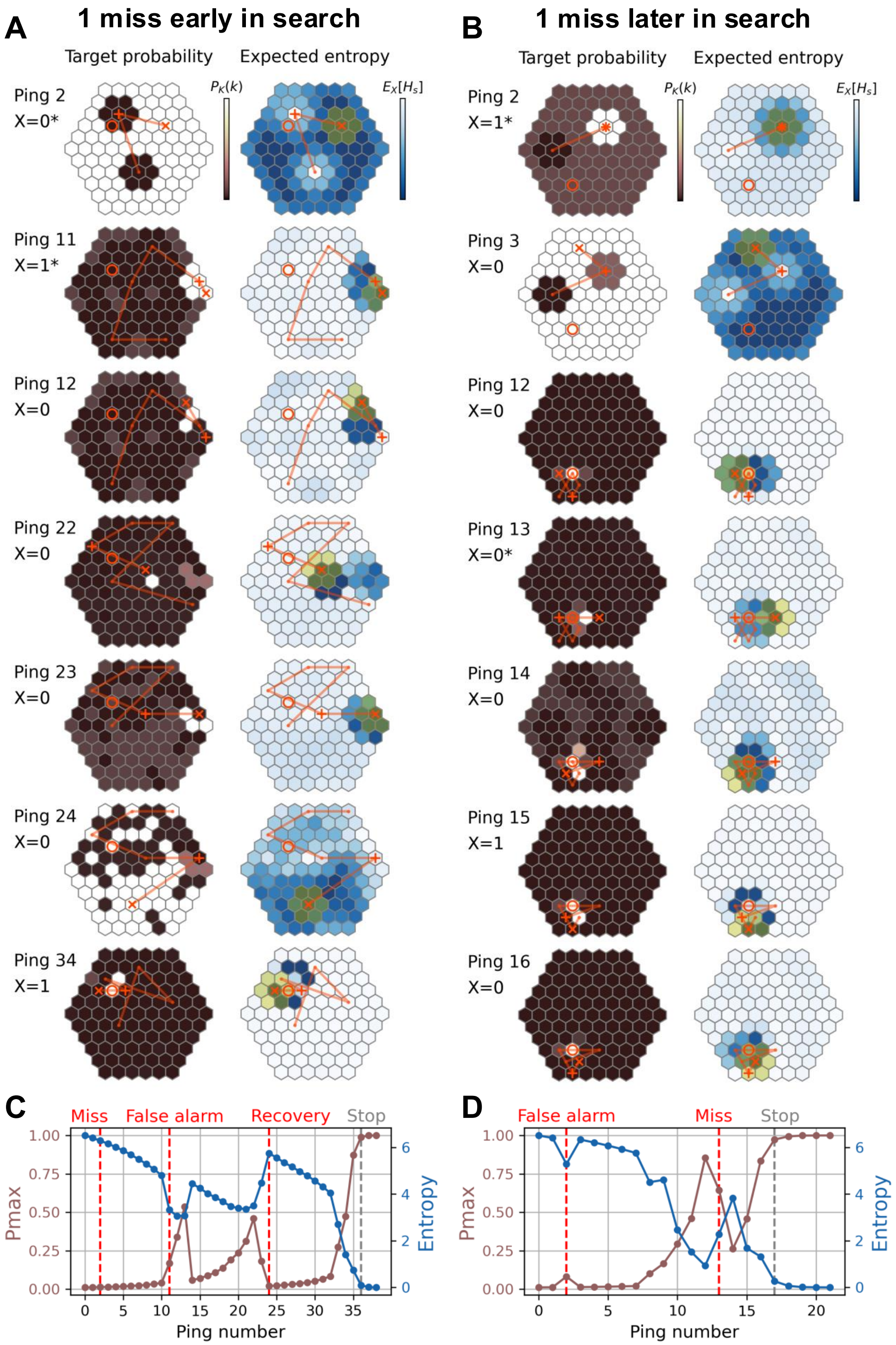


FIG C: **Example infotaxis searches with one miss and one false alarm.** (A, C) An example search with one miss occurring early (ping 2) in the search, which lengthens the search substantially, even though the impact of false alarm is mitigated quickly. (B, D) An example search with one miss occurring later (ping 13) in the search, causing a transient rise in the entropy curve. Color represents a relative scale in the value range spanned within the search space. Under infotaxis, only the relative scale matters because the agent selects the next set of actions that gives the lowest expected entropy among all choices. See Fig 3 caption for definitions of all symbols, Table B in S3 Appendix for all search parameter values, and S8–S9 Videos for the full sequences.

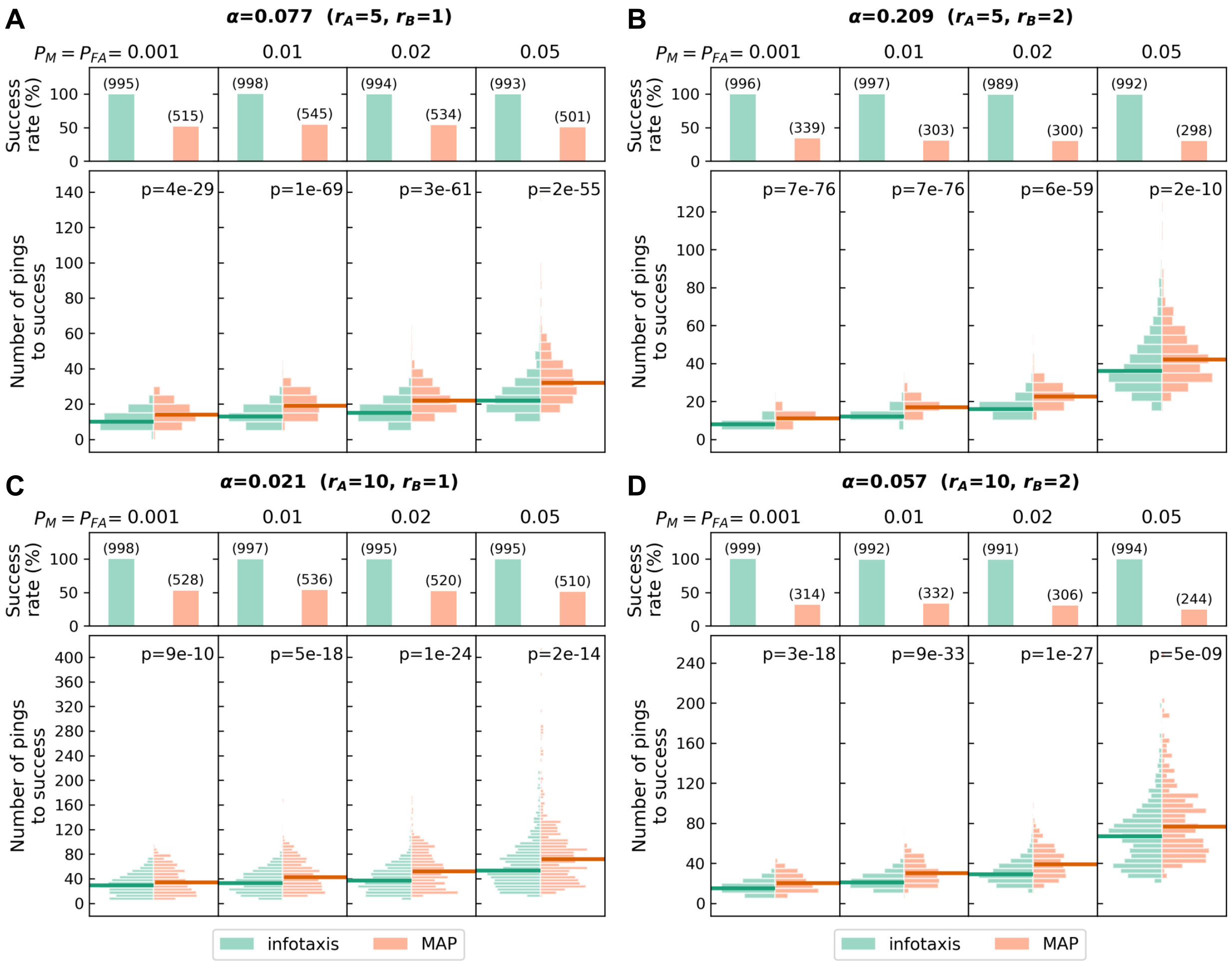


FIG D: **Performance of infotaxis and MAP searches with the stopping condition: (1) $p_{th} = 0.99$ or (2) the change of $P_{max}$ across three consecutive pings is smaller than $10^{-5}$.** The search parameter combinations in panels A-D are identical to those in Fig 4, except for $p_{th} = 0.95$ in Fig 4.

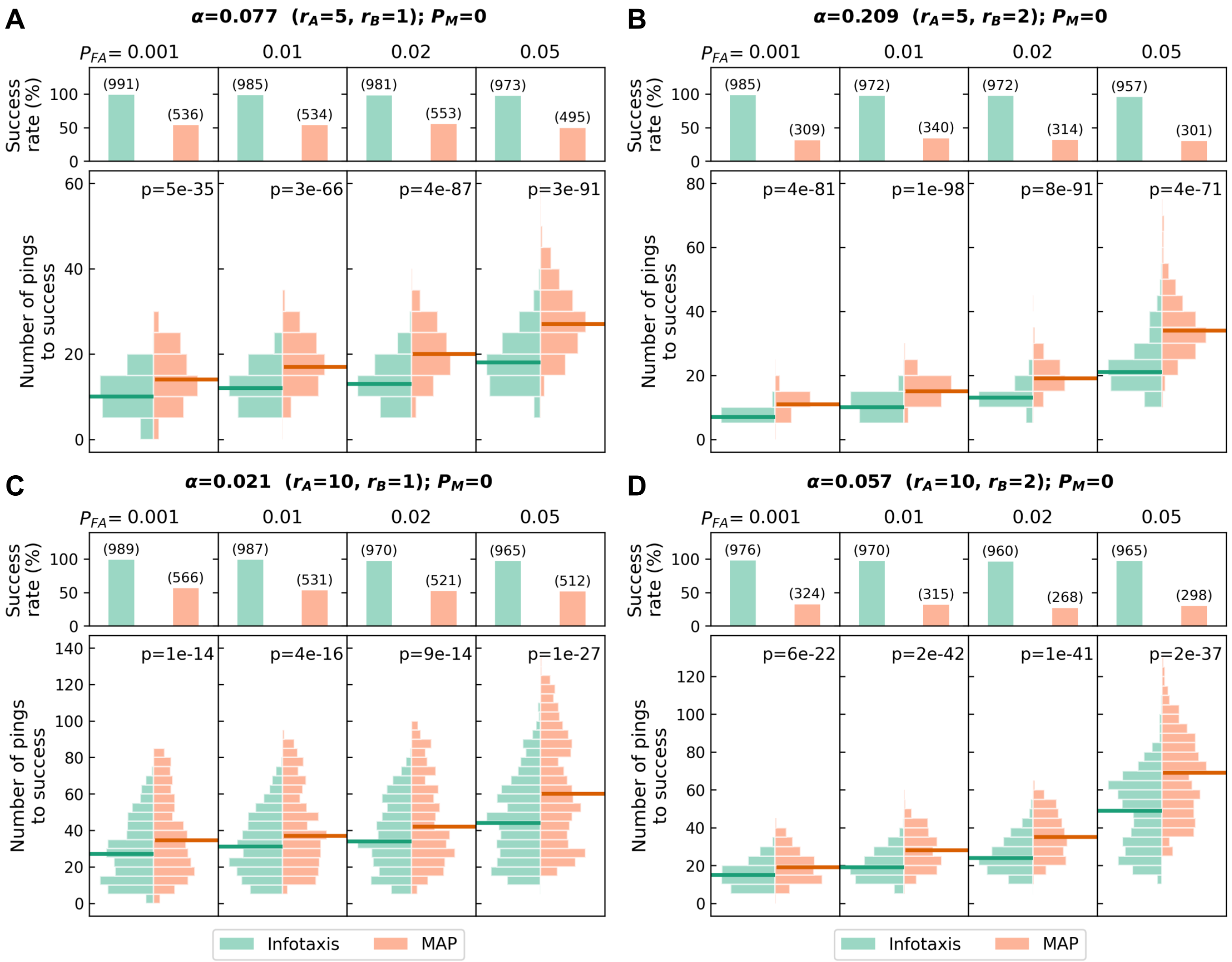


FIG E: **Comparison of infotaxis and MAP searches with varying $P_{FA}$ while fixing $P_M = 0$.** Success rate (top) and number of echolocation pings required (bottom) to locate the target under different search space and beam parameters. Elevated $P_{FA}$ increases the overall number of pings required to complete a search. See Sec. IID for definitions of all quantities and details on the stopping condition.

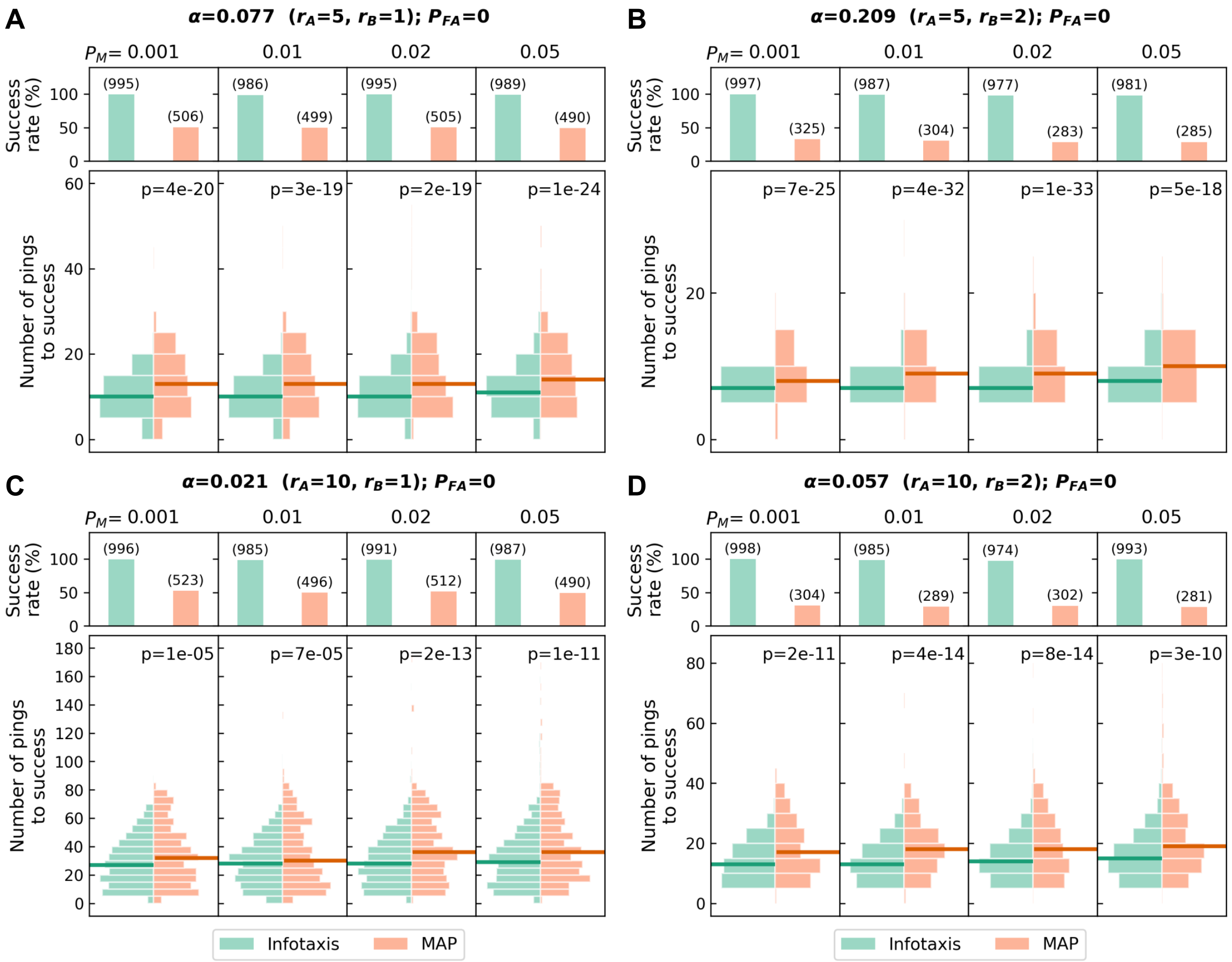


FIG F: **Comparison of infotaxis and MAP searches with varying $P_M$ while fixing $P_{FA} = 0$.** Success rate (top) and number of pings required (bottom) to locate the target under different search space and beam parameters. Elevated $P_M$ increases the number of outlier runs that require a large number of pings to complete a search. See Sec. IID for definitions of all quantities and details on the stopping condition.

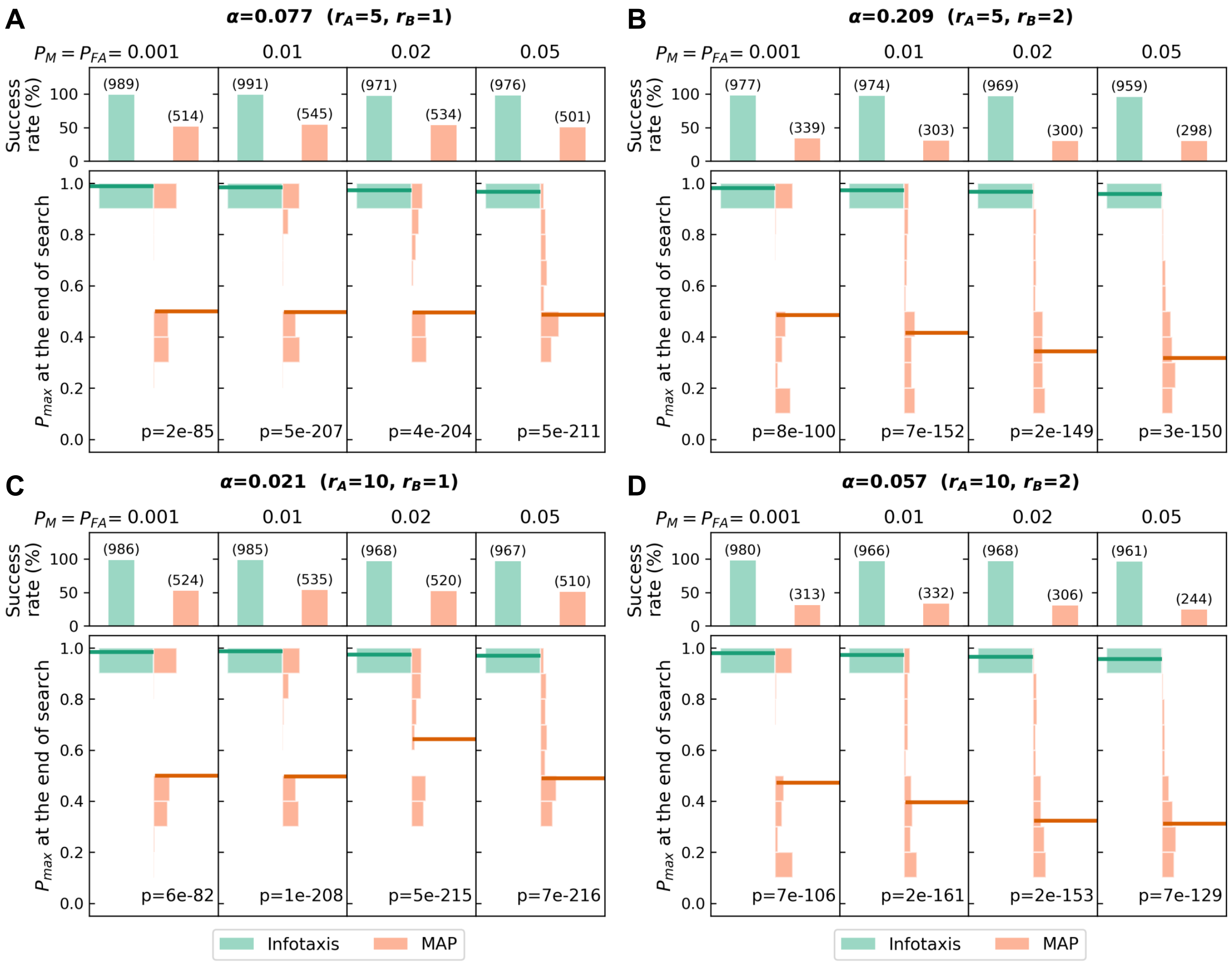


FIG G: **Comparison of the distributions of $P_{max}$ at the end of the search for infotaxis and MAP.** The data plotted are from the same simulation runs shown in Fig 4.

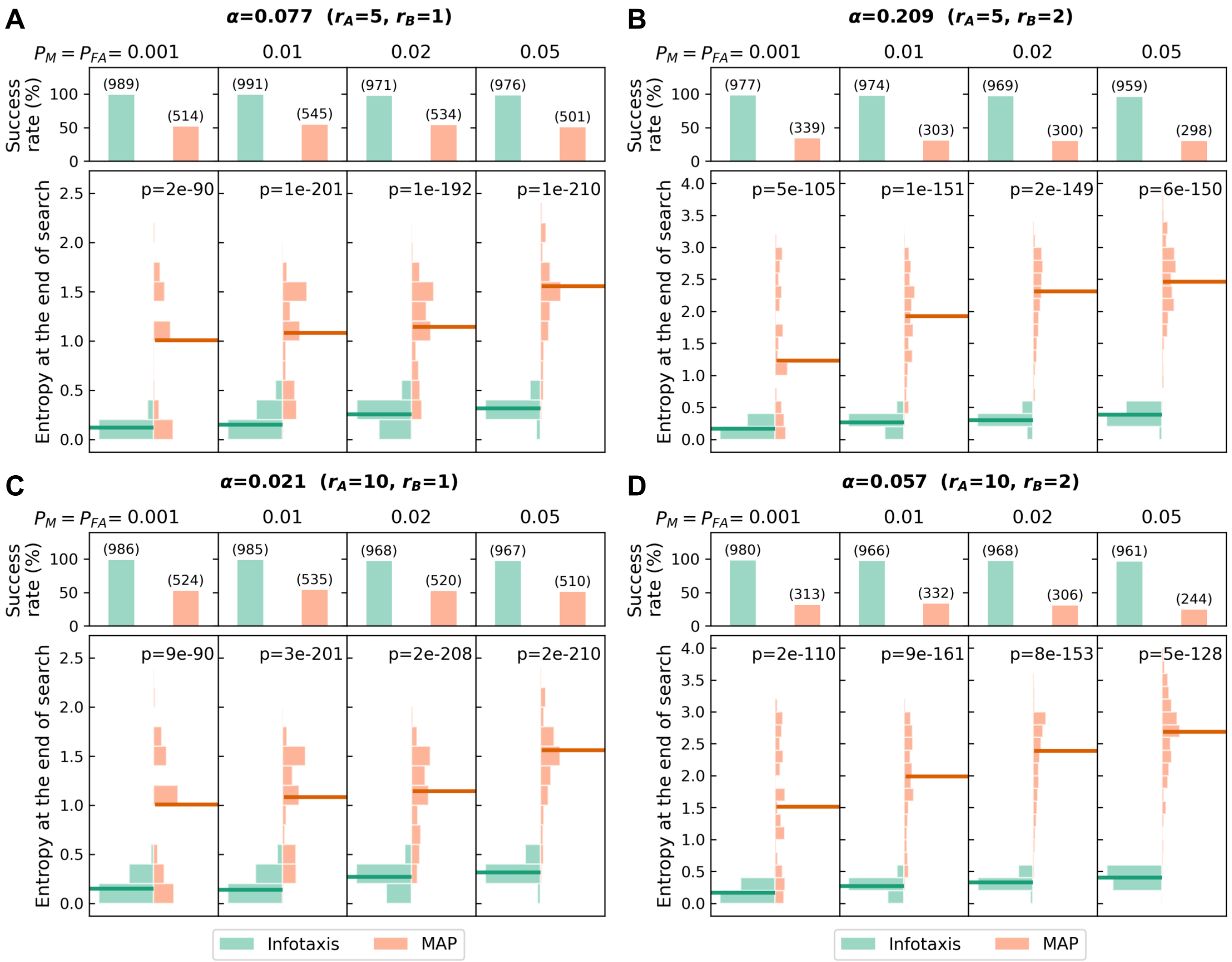


FIG H: **Comparison of the distributions of entropy $H[K]$ at the end of the search for infotaxis and MAP.** The data plotted are from the same simulation runs shown in Fig 4.

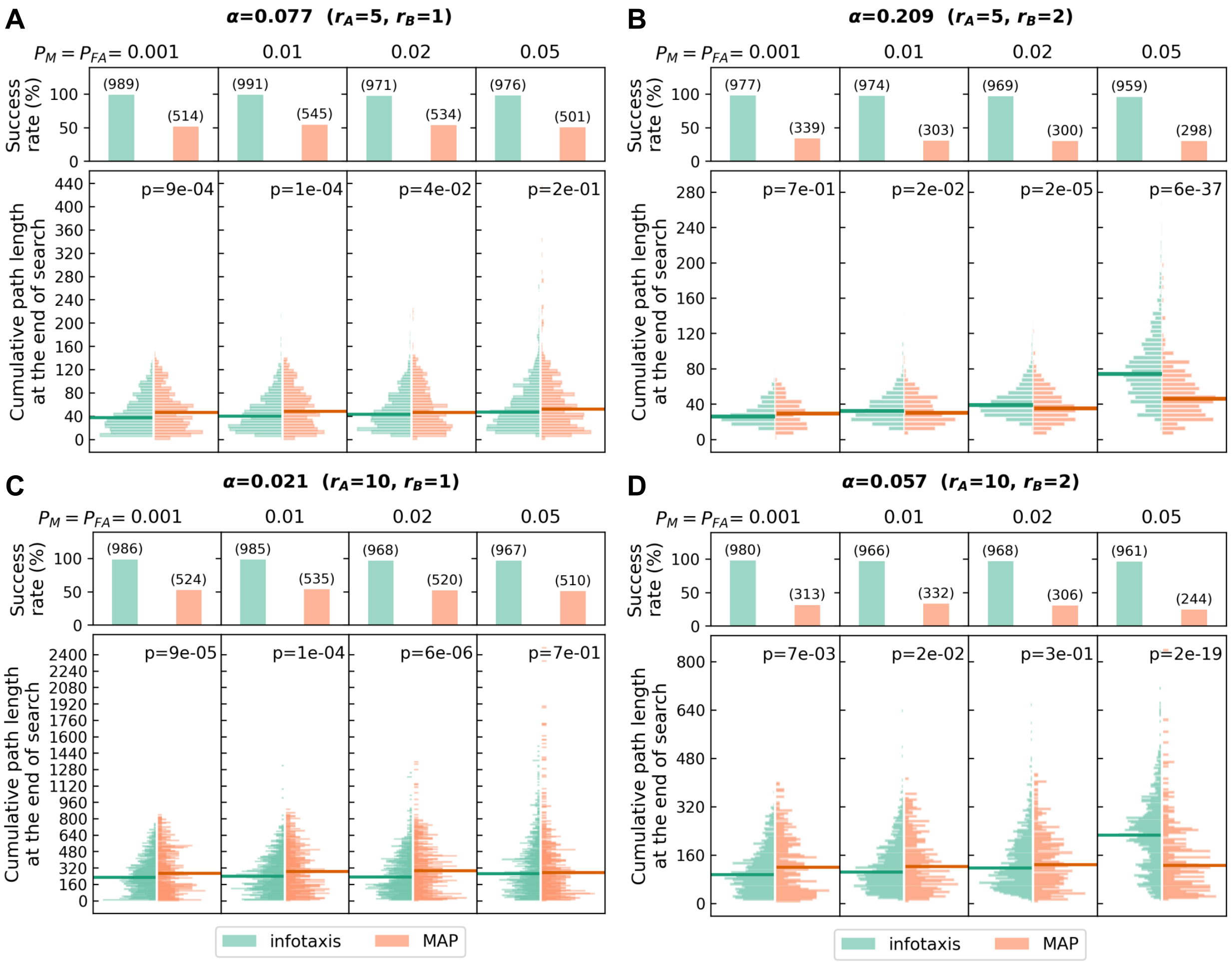


FIG I: **Comparison of the distributions of cumulative beam aim path length until the end of the search for infotaxis and MAP.** The data plotted are from the same simulation runs shown in Fig 4.

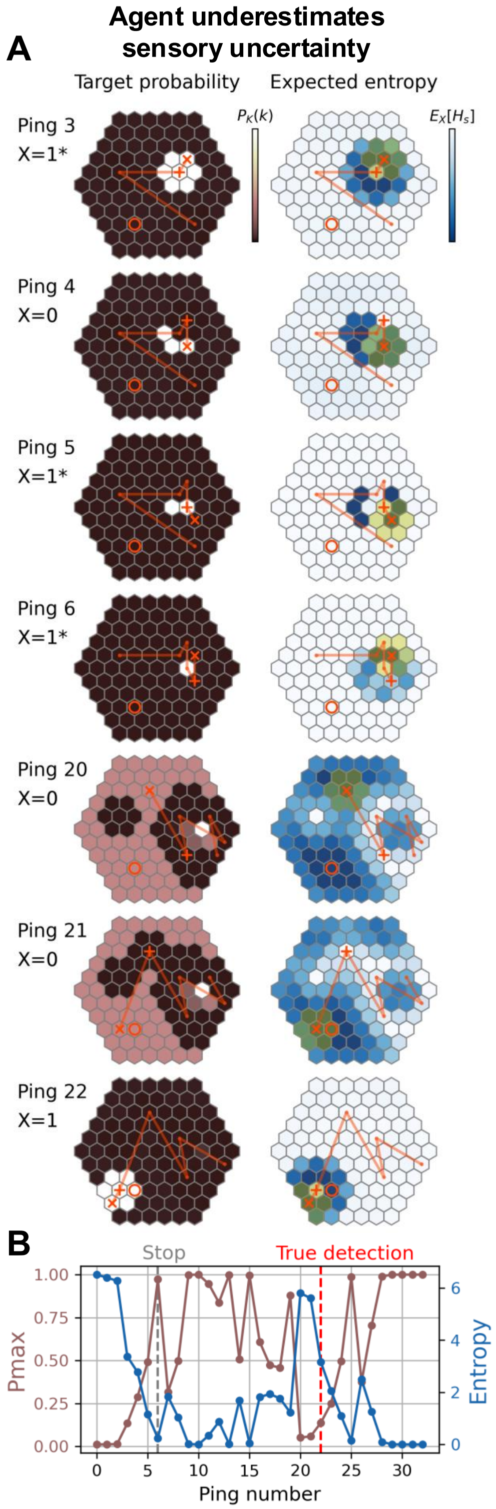


FIG J: **An example infotaxis search in which the agent underestimates the sensory uncertainty.** Due to the assumed lower sensory uncertainty, $P_{max}$ exceeds the threshold $p_{th} = 0.95$ too early, causing the agent to terminate the search prematurely at an incorrect location. Color represents a relative scale in the value range spanned within the search space. Under infotaxis, only the relative scale matters because the agent selects the next set of actions that gives the lowest expected entropy among all choices. See Fig 3 caption for definitions of all symbols, Table B in S3 Appendix for all search parameter values, and S10 Video for the full sequence.

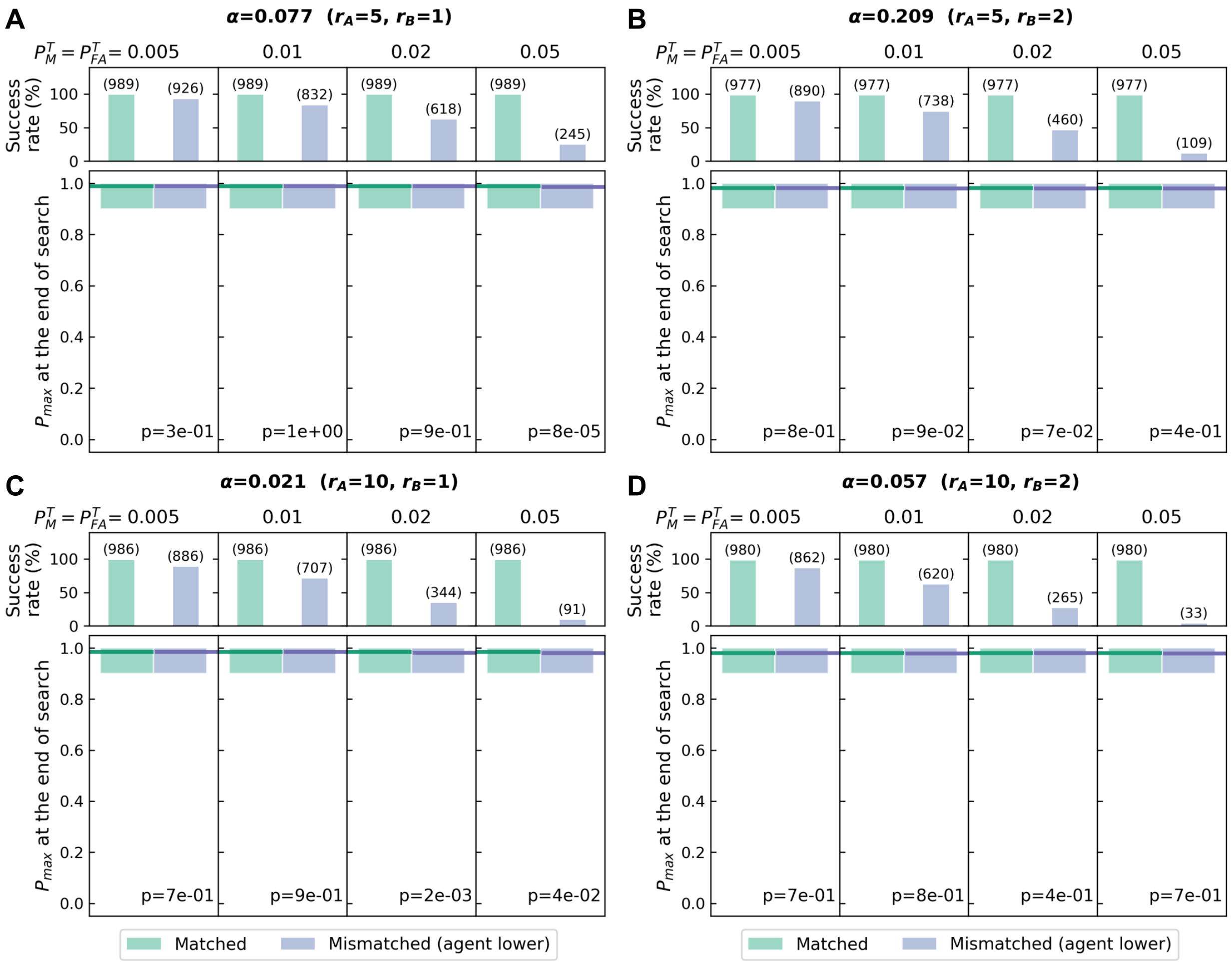


FIG K: **Comparison of the distributions of $P_{max}$ at the end of the search when the infotaxis agent underestimates the sensory uncertainty.** The data plotted are from the same simulation runs shown in Fig 5.

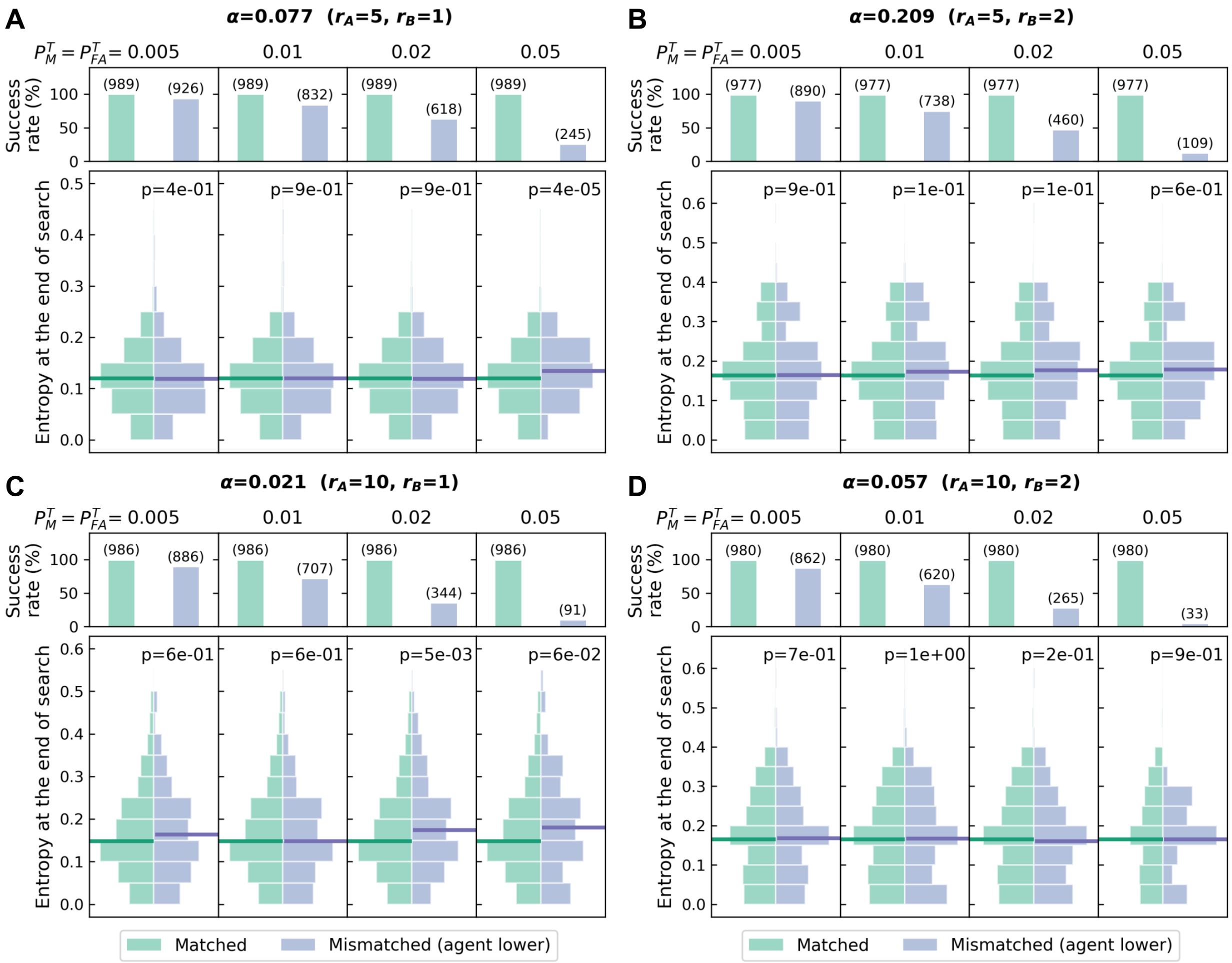


FIG L: **Comparison of the distributions of entropy $H[K]$ at the end of the search when the infotaxis agent underestimates the sensory uncertainty.** The data plotted are from the same simulation runs shown in Fig 5.

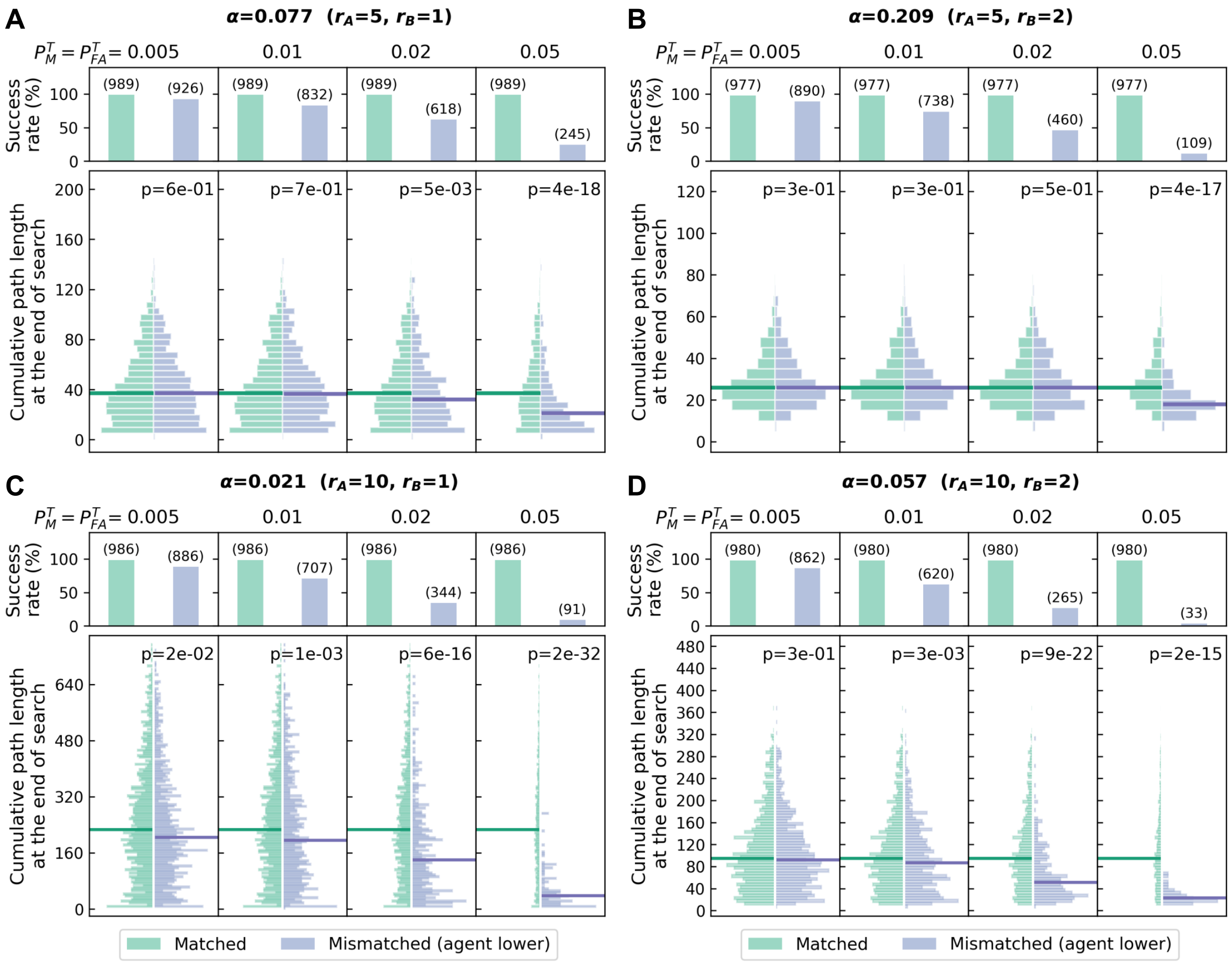


FIG M: **Comparison of the distributions of cumulative beam aim path length until the end of the search when the infotaxis agent underestimates the sensory uncertainty.** The data plotted are from the same simulation runs shown in Fig 5.

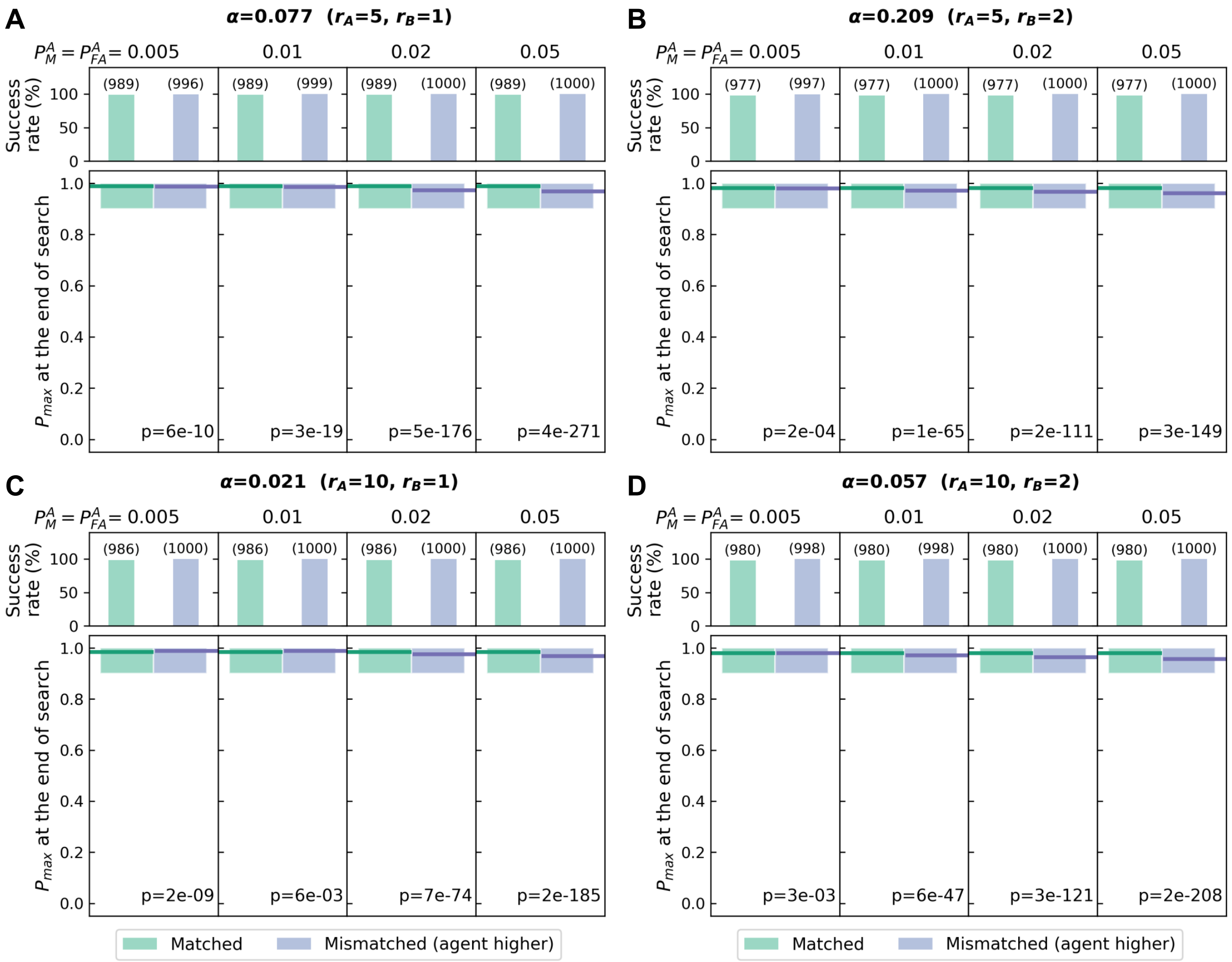


FIG N: **Comparison of the distributions of $P_{max}$ at the end of the search when the infotaxis agent overestimates the sensory uncertainty.** The data plotted are from the same simulation runs shown in Fig 6.

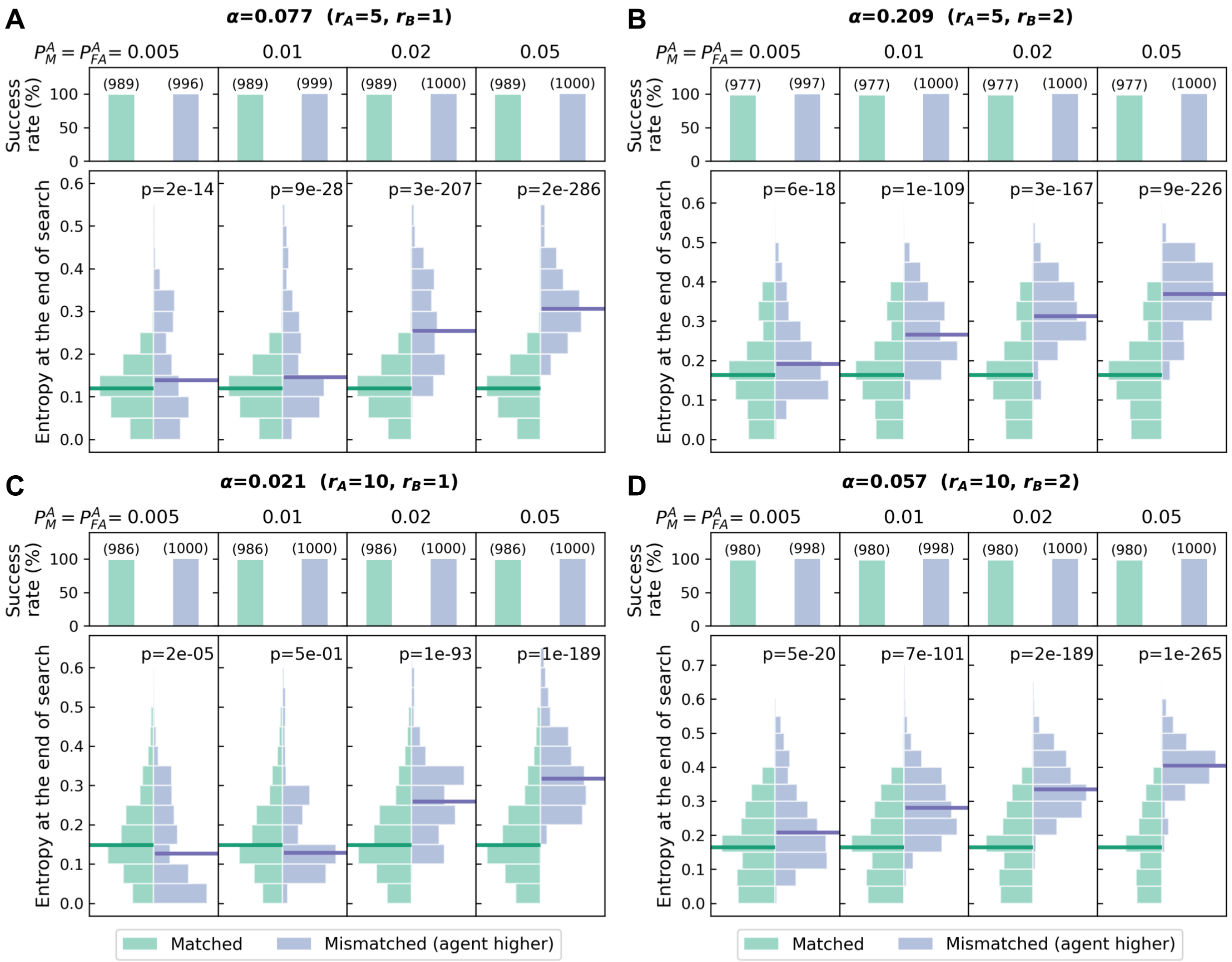


FIG O: **Distribution of entropy $H[K]$ at the end of the search when the infotaxis agent overestimates sensory uncertainty.** The data plotted are from the same simulation runs shown in Fig 6.

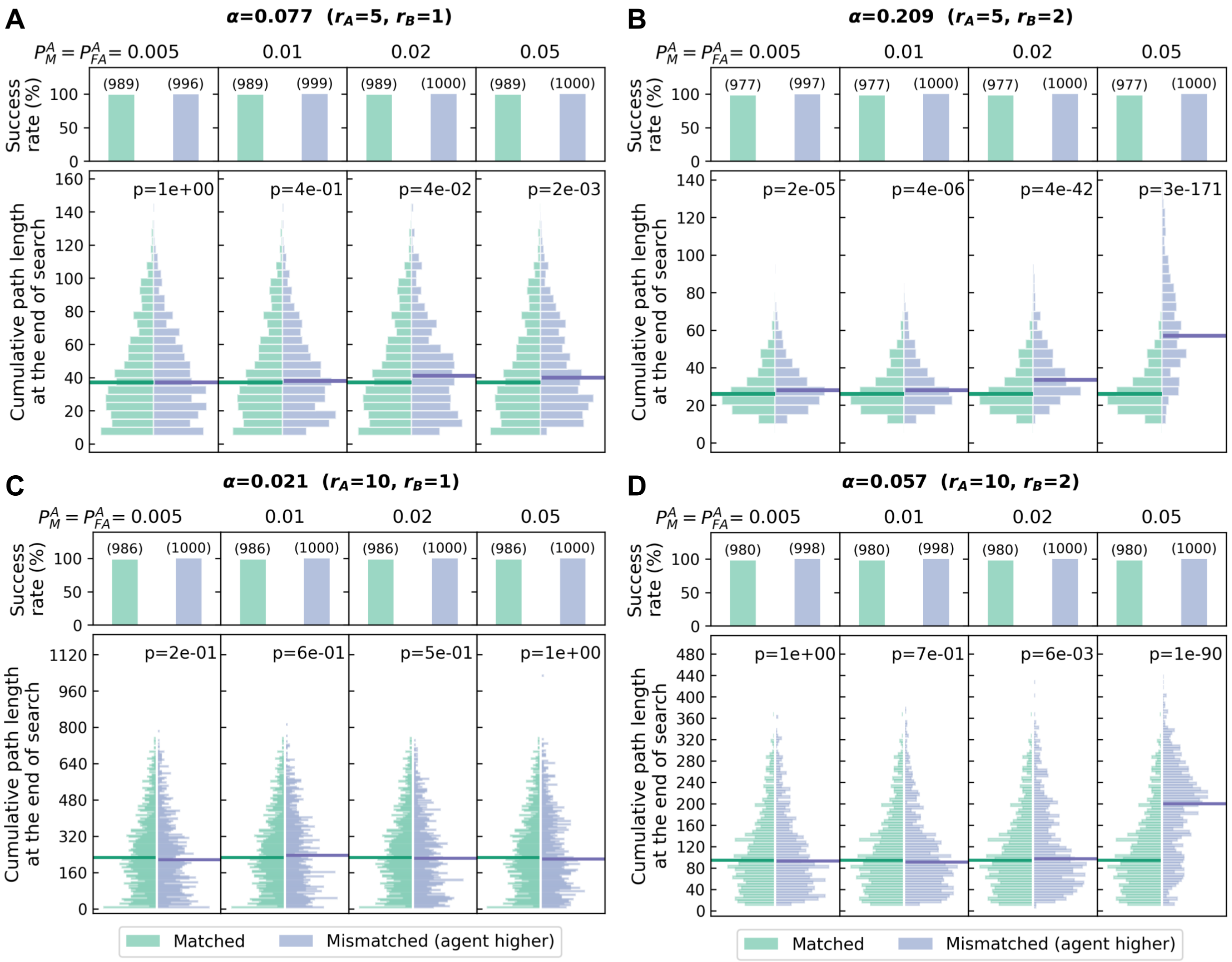


FIG P: **Comparison of the distributions of cumulative beam aim path length until the end of the search when the infotaxis agent overestimates sensory uncertainty.** The data plotted are from the same simulation runs shown in Fig 6.

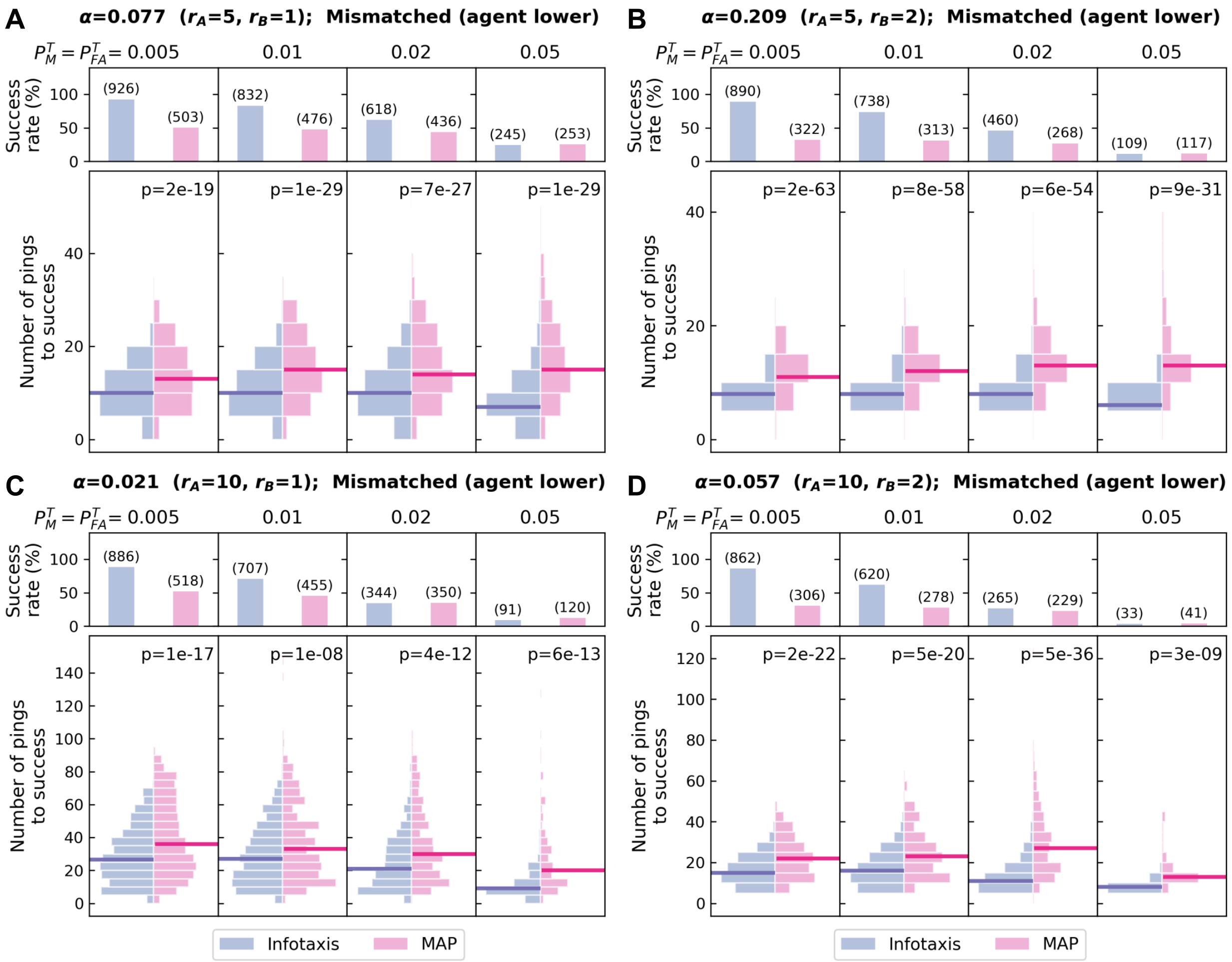


FIG Q: **Performance comparison of infotaxis and MAP searches when the agent underestimates sensory uncertainty.** All other plot and annotation details are identical to those of Fig 4. See Sec. IID for definitions of all quantities and details on the stopping condition.

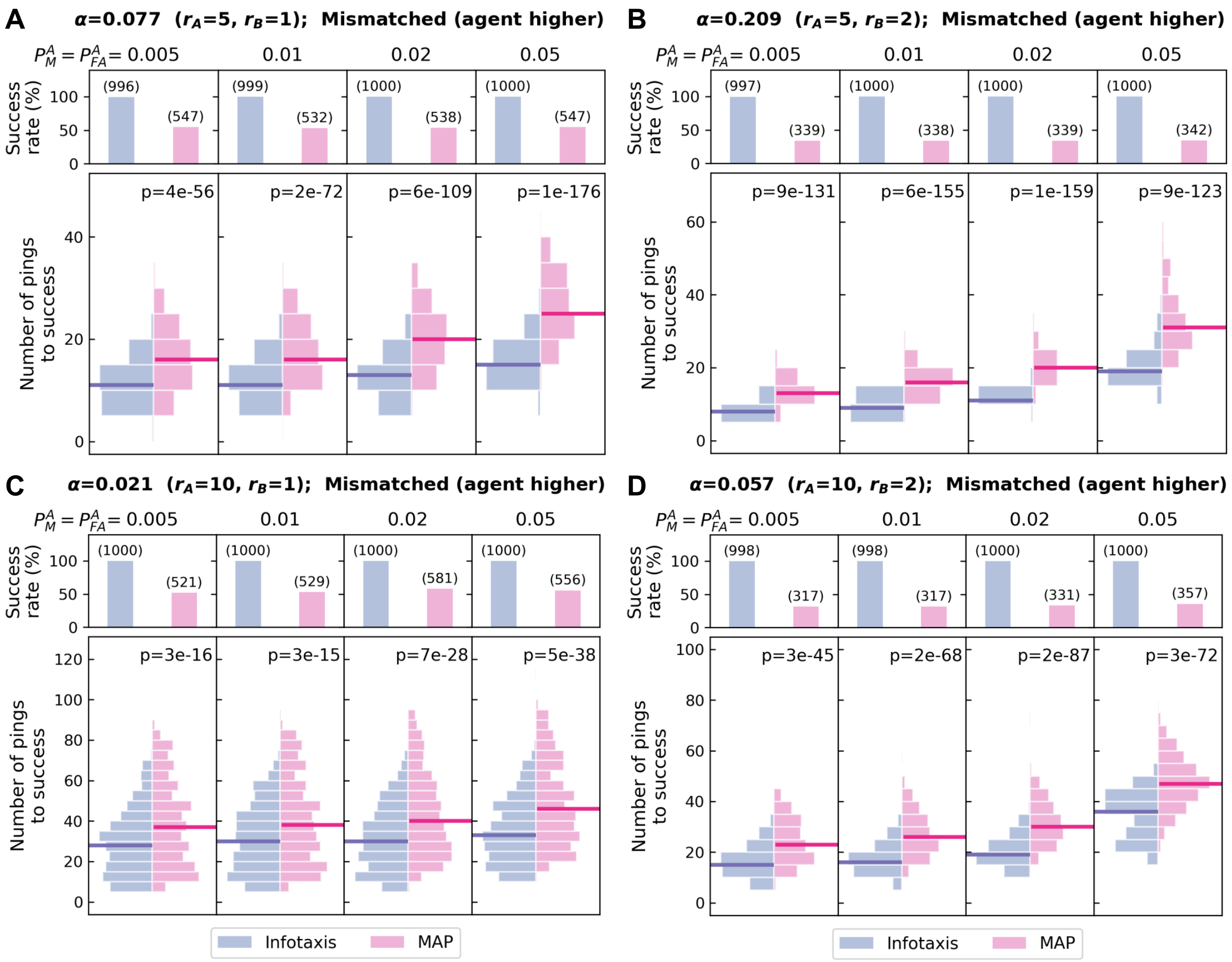


FIG R: **Performance comparison of infotaxis and MAP searches when the agent overestimates sensory uncertainty.** All other plot and annotation details are identical to those of Fig 4. See Sec. IID for definitions of all quantities and details on the stopping condition.

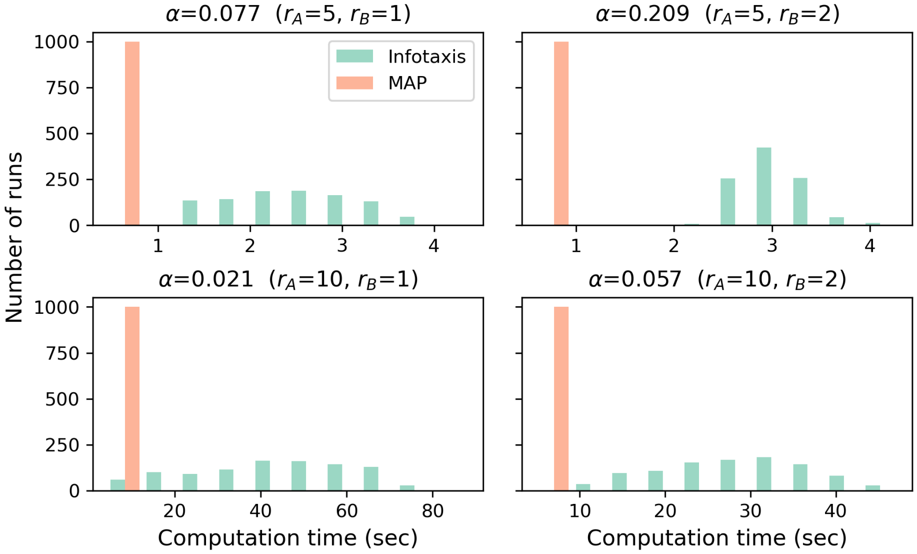


FIG S: **Comparison of computation time for infotaxis and MAP searches.** Aggregated computation time taken by each infotaxis and MAP search measured by the Python function `time.process_time()`, which is the sum of the system and user CPU time without time elapsed during sleep. $P_M = P_{FA} = 0.001$ in these simulations. See Sec. IID for definitions of all quantities and details on the stopping condition, and Table B in S3 Appendix for all search parameter values.

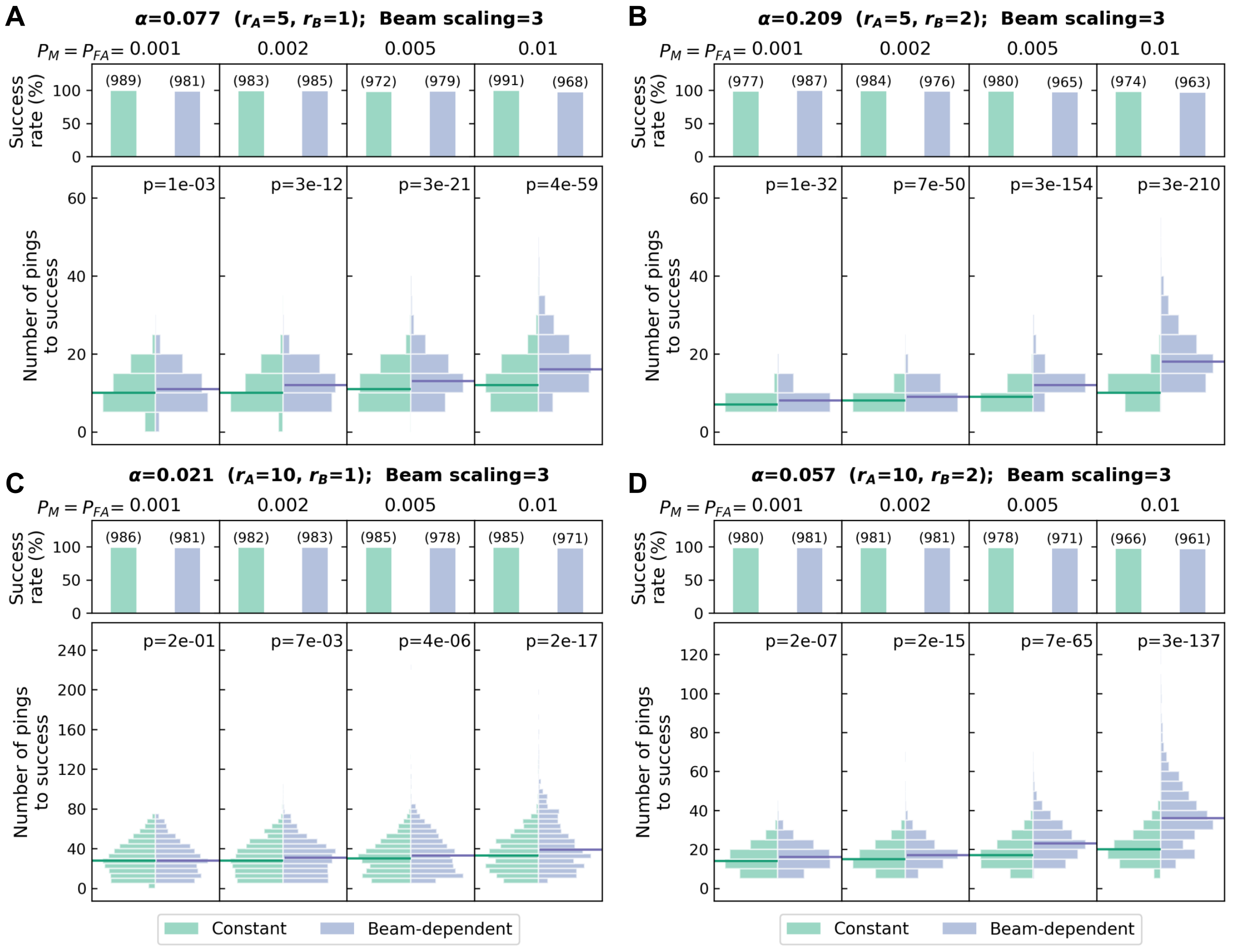


FIG T: **Effects of using beampattern-modulated $P_M$ and $P_{FA}$ on infotaxis search.** The beampattern magnitude modulation is $\lambda_B = 3$, with the modulation width controlled by $\sigma_B = 0.5$ and $\sigma_B = 1$ for $r_B = 1$ and $r_B = 2$, respectively (see Section A in S1 Appendix for detail). The reported $p$-values are from Mann-Whitney U tests under the alternative hypothesis that the number of pings required to locate the target are the same. All other plot and annotation details are identical to those of Fig 4.

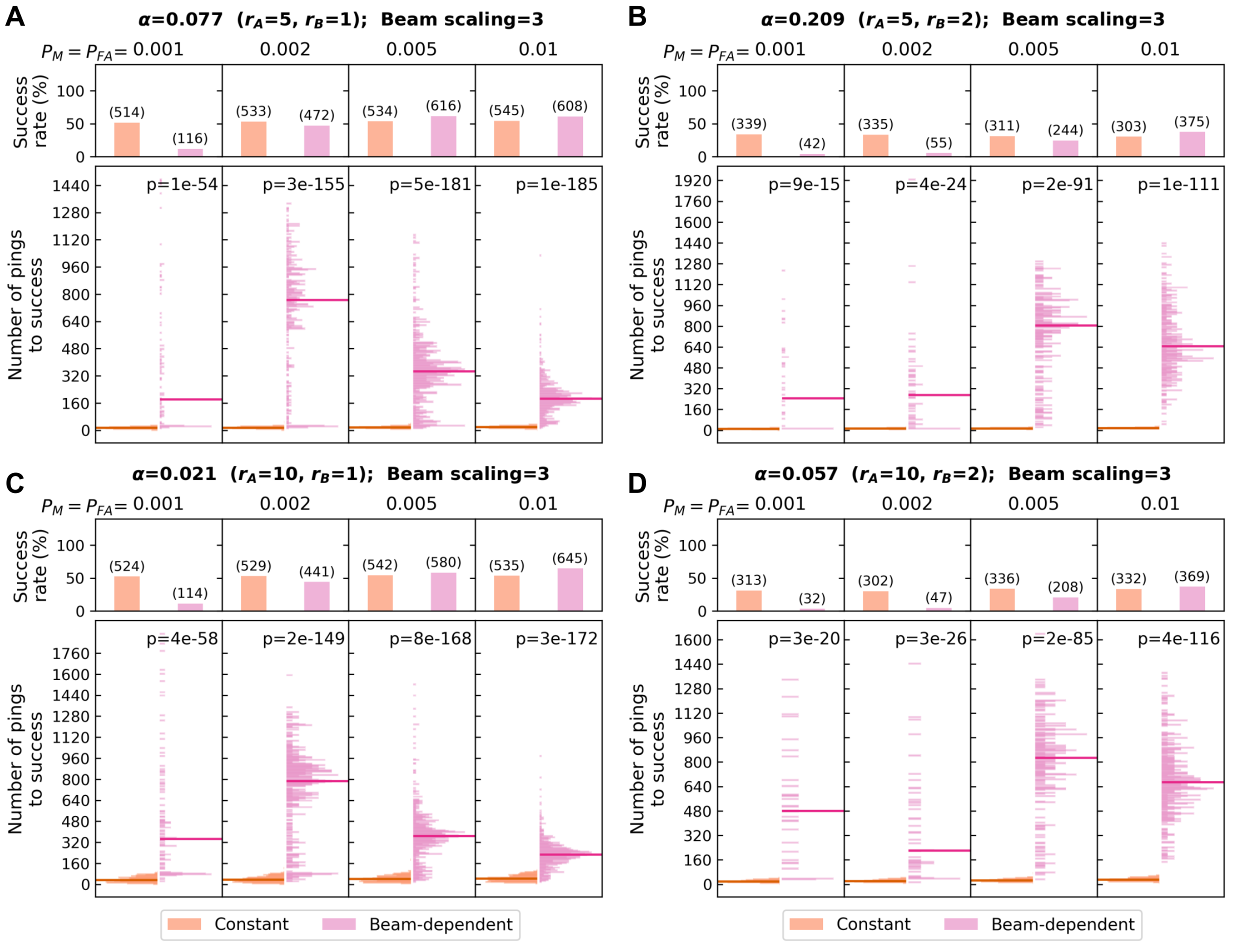


FIG U: **Effects of using beampattern-modulated $P_M$ and $P_{FA}$ on MAP search.** The beampattern magnitude modulation is $\lambda_B = 3$, with the modulation width controlled by $\sigma_B = 0.5$ and $\sigma_B = 1$ for $r_B = 1$ and $r_B = 2$, respectively (see Section A in S1 Appendix for detail). Note that a search is considered failed when the maximum ping number of 2000 is reached and neither of the stopping criteria are met. The reported $p$-values are from Mann-Whitney U tests under the alternative hypothesis that the number of pings required to locate the target are the same. All other plot and annotation details are identical to those of Fig 4.
