## Supplementary material for "Modeling echolocation as an active pursuit of information via infotaxis": S3 Appendix (all supporting tables)

### S3 APPENDIX: SUPPORTING TABLES

TABLE A: Model parameters used for all main text figures. When $P_M^T$ and $P_{FA}^T$ are not noted separately, they share the same values as $P_M^A$ and $P_M^A$.

| **Figure** | $r_A$ | $N_A$ | $r_B$ | $N_B$ | $P_M^A = P_{FA}^A$ | $P_M^T = P_{FA}^T$ |
|---|---|---|---|---|---|---|
| Fig 2 | 5 | 91 | 1 | 7 | 0.005 | — |
| Fig 3A/3F | 5 | 91 | 1 | 7 | 0.005 | — |
| Fig 3B/3G | 8 | 217 | {1, 2} | {7, 19} | 0.001 | — |
| Fig 3C/3H | 8 | 217 | {1, 2, 3} | {7, 19, 37} | 0.001 | — |
| Fig 3D/3I | 5 | 91 | 1 | 7 | 0.001 | — |
| Fig 3E/3J | 5 | 91 | 1 | 7 | 0.001 | — |
| Fig 4 | {5, 10} | {91, 331} | {1, 2} | {7, 19} | {0.001, 0.01, 0.02, 0.05} | — |
| Fig 5 | {5, 10} | {91, 331} | {1, 2} | {7, 19} | 0.001 | {0.005, 0.01, 0.02, 0.05} |
| Fig 6 | {5, 10} | {91, 331} | {1, 2} | {7, 19} | {0.005, 0.01, 0.02, 0.05} | 0.001 |
| Fig 7A | 5 | 91 | 1 | 7 | 0.005 | — |
| Fig 7B | {5, 6, 7, 8, 9, 10, 11, 12, 13, 14, 15} | {91, 127, 169, 217, 271, 331, 397, 469, 547, 631, 721} | 1 | 7 | 0 | — |
| Fig 8A | 5 | 91 | 1 | 7 | {0.001, 0.02} | — |
| Fig 8B/8C | {5, 6, 7, 8, 9, 10} | {91, 127, 169, 217, 271, 331} | {1, 2} | {7, 19} | $P_M^A$=0.001, $P_{FA}^A$ sweep | — |
| Fig 9A/9B | 5 | 91 | 1 | 7 | 0.005 | — |
| Fig 9C | 5 | 91 | 1 | 7 | {0.001, 0.01, 0.02, 0.05} | — |

TABLE B: Model parameters used for all supporting figures in S2 Appendix. When $P_M^T$ and $P_{FA}^T$ are not noted separately, they share the same values as $P_M^A$ and $P_{FA}^A$.

| **Figure** | $r_A$ | $N_A$ | $r_B$ | $N_B$ | $P_M^A$ | $P_{FA}^A$ | $r_M$ | $N_M$ | $P_M^T = P_{FA}^T$ |
|---|---|---|---|---|---|---|---|---|---|
| Panel A of Fig A | 5 | 91 | 1 | 7 | 0.001 | $= P_M^A$ | 3 | 37 | — |
| Panel B of Fig A | 5 | 91 | 1 | 7 | {0.001, 0.01, 0.02, 0.05} | $= P_M^A$ | {3,2} | {37, 19} | — |
| Panel C of Fig A | 10 | 331 | 1 | 7 | {0.001, 0.01, 0.02, 0.05} | $= P_M^A$ | {3,2} | {37, 19 } | — |
| Panel A of Fig B | 5 | 91 | 1 | 7 | 0 | {0.001, 0.05, 0.1, 0.15, 0.5} | — | — | — |
| Panel B of Fig B | 5 | 91 | 2 | 19 | 0 | {0.001, 0.035, 0.05, 0.1, 0.3} | — | — | — |
| Panel A of Fig C | 5 | 91 | 1 | 7 | 0.01 | $= P_M^A$ | — | — | — |
| Panel B of Fig C | 5 | 91 | 1 | 7 | 0.01 | $= P_M^A$ | — | — | — |
| Figs D | {5, 10} | {91, 331} | {1, 2} | {7, 19} | {0.001, 0.01, 0.02, 0.05} | $= P_M^A$ | — | — | — |
| Fig E | {5, 10} | {91, 331} | {1, 2} | {7, 19} | 0 | {0.001, 0.01, 0.02, 0.05} | — | — | — |
| Fig F | {5, 10} | {91, 331} | {1, 2} | {7, 19} | {0.001, 0.01, 0.02, 0.05} | 0 | — | — | — |
| Figs G/H/I | {5, 10} | {91, 331} | {1, 2} | {7, 19} | {0.001, 0.01, 0.02, 0.05} | $= P_M^A$ | — | — | — |
| Fig J | {5, 10} | {91, 331} | {1, 2} | {7, 19} | 0.001 | $= P_M^A$ | — | — | 0.05 |
| Figs K/L/M | {5, 10} | {91, 331} | {1, 2} | {7, 19} | 0.001 | $= P_M^A$ | — | — | {0.005, 0.01, 0.02, 0.05} |
| Figs N/O/P | {5, 10} | {91, 331} | {1, 2} | {7, 19} | {0.005, 0.01, 0.02, 0.05} | $= P_M^A$ | — | — | 0.001 |
| Fig Q | {5, 10} | {91, 331} | {1, 2} | {7, 19} | 0.001 | $= P_M^A$ | — | — | {0.005, 0.01, 0.02, 0.05} |
| Fig R | {5, 10} | {91, 331} | {1, 2} | {7, 19} | {0.005, 0.01, 0.02, 0.05} | $= P_M^A$ | — | — | 0.001 |
| Fig S | {5, 10} | {91, 331} | {1, 2} | {7, 19} | 0.001 | $= P_M^A$ | — | — | — |
| Figs T/U | {5, 10} | {91, 331} | {1, 2} | {7, 19} | {0.001, 0.01, 0.02, 0.05} | $= P_M^A$ | — | — | — |
